# Synthetic cell-based materials extract positional information from morphogen gradients

**DOI:** 10.1101/2021.04.25.441320

**Authors:** Aurore Dupin, Lukas Aufinger, Igor Styazhkin, Florian Rothfischer, Benedikt Kaufmann, Sascha Schwarz, Nikolas Galensowske, Hauke Clausen-Schaumann, Friedrich C. Simmel

**Affiliations:** Physics Department – E14, TU Munich, 85748 Garching, Germany; Center for NanoScience – CeNS, Schellingstraße 4, 80799 Munich, Germany; Center for Applied Tissue Engineering and Regenerative Medicine – CANTER, Munich University of Applied Sciences, Lothstrasse 34, 80335 Munich, Germany; Chair of Applied Mechanics, TU Munich, 85748 Garching, Germany

## Abstract

Dynamic biomaterials composed of synthetic cellular structures have the potential to adapt and functionally differentiate guided by physical and chemical cues from their environment. Inspired by developing biological systems, which efficiently extract positional information from chemical morphogen gradients in the presence of environmental uncertainties, we here investigate the analogous question: how well can a synthetic cell determine its position within a synthetic multicellular structure? In order to calculate positional information in such systems, we created and analyzed a large number of replicas of synthetic cellular assemblies, which were composed of emulsion droplets connected via lipid bilayer membranes. The droplets contained cell-free two-node feedback gene circuits that responded to gradients of a genetic inducer acting as a morphogen. We found that in our system, simple anterior-posterior differentiation is possible, but positional information is limited by gene expression noise, and is also critically affected by the temporal evolution of the morphogen gradient and the life-time of the cell-free expression system contained in the synthetic cells. Using a 3D printing approach, we demonstrate morphogen-based differentiation also in larger tissue-like assemblies.

---

Biological development - the generation of a complex, differentiated organism starting from a single cell - is a striking example of self-organization in biology that has inspired chemists, materials scientists, molecular programmers and synthetic biologists alike to envision autonomously developing, self-differentiating and self-sustaining biomimetic systems ^1-11^. Newly available techniques such as 3D printing of soft materials have opened up the possibility to automate and standardize the assembly of novel materials, where properties are defined across scales by combining top-down specification via additive manufacturing with bottom-up pattern formation via molecular self-organization. In this context, researchers have recently begun to create artificial multicellular structures ^12-14^, which may form the basis of biomaterials capable of differentiating into functionally distinct regions, based on external chemical and physical cues.

A common mechanism for biological patterning utilizes the information supplied by morphogen gradients that are interpreted by gene regulatory circuits to infer the position of cells within the developing organism ^15^. In a similar way, synthetic morphogen gradients might be utilized in the context of synthetic cell-based materials that host engineered gene circuits for pattern formation. To reproducibly manufacture such materials, it will be important to understand the potential and limitations of morphogen-based self-organization in such systems ^16^. As position determines the fate of the synthetic cells and the future organization of the biomaterial – together with its desired functionality -, developmental processes have to take place robustly in the presence of stochastic variations in both external and internal parameters, e.g., the size of the artificial tissue, the gradient profile, or gene expression noise ^17^.

The ability of a genetic circuit to determine position within a system – in the presence of noise - can be precisely quantified by calculating the “positional information” (PI) based on the “mutual information” between gene expression levels and position within the organism ^18, 19^. In the present work we apply the PI concept in a synthetic context by characterizing the response of synthetic multicellular systems to the presence of a morphogen gradient formed by a diffusible genetic inducer.

To this end, permeable tissues of synthetic cells are equipped with different kinds of cell-free gene circuits – comprised of different combinations of genetic repressors - that, depending on the circuit topology, respond to this gradient in different ways. Backed up by numerical modelling, we find that the extracted PI depends on the circuit’s response function relative to the morphogen gradient and the temporal evolution and lifetime of the cell-free gene expression reactions in the presence of gene expression noise. Quantitation of positional information is thus shown to provide a rational approach for the characterization and engineering of developmental processes in synthetic cell-based biomaterials. To explore the route towards the automated and standardized manufacturing of differentiated synthetic tissues, we also apply our results obtained from 1D differentiation studies to investigate the evolution of a developmental circuit within a synthetic multicellular structure fabricated with a custom-made multipurpose bioprinting platform.

A summary of our approach is shown in Figure 1. Our experimental system consists of linear assemblies of water-in-oil droplets serving as prototypical synthetic cellular compartments. Multicellular systems are formed by bringing lipid monolayer-enclosed compartments into contact with each other using a micromanipulator, which creates a permeable lipid bilayer between adjacent cells as previously described (**Fig. 1a**) ^20-22^. Using this technique, we generated assemblies of six droplets each, in which one of the terminal cells contained a genetic inducer (IPTG) acting as the morphogen and the other cells contained a cell-free expression system and a genetic circuit responding to the morphogen. Diffusion of the inducer from the “organizer” or “sender” cell into the array creates a morphogen gradient across the “receiver” cells that elicits a position-dependent gene expression response (**Fig. 1a, SI video**).

**Figure 1.**
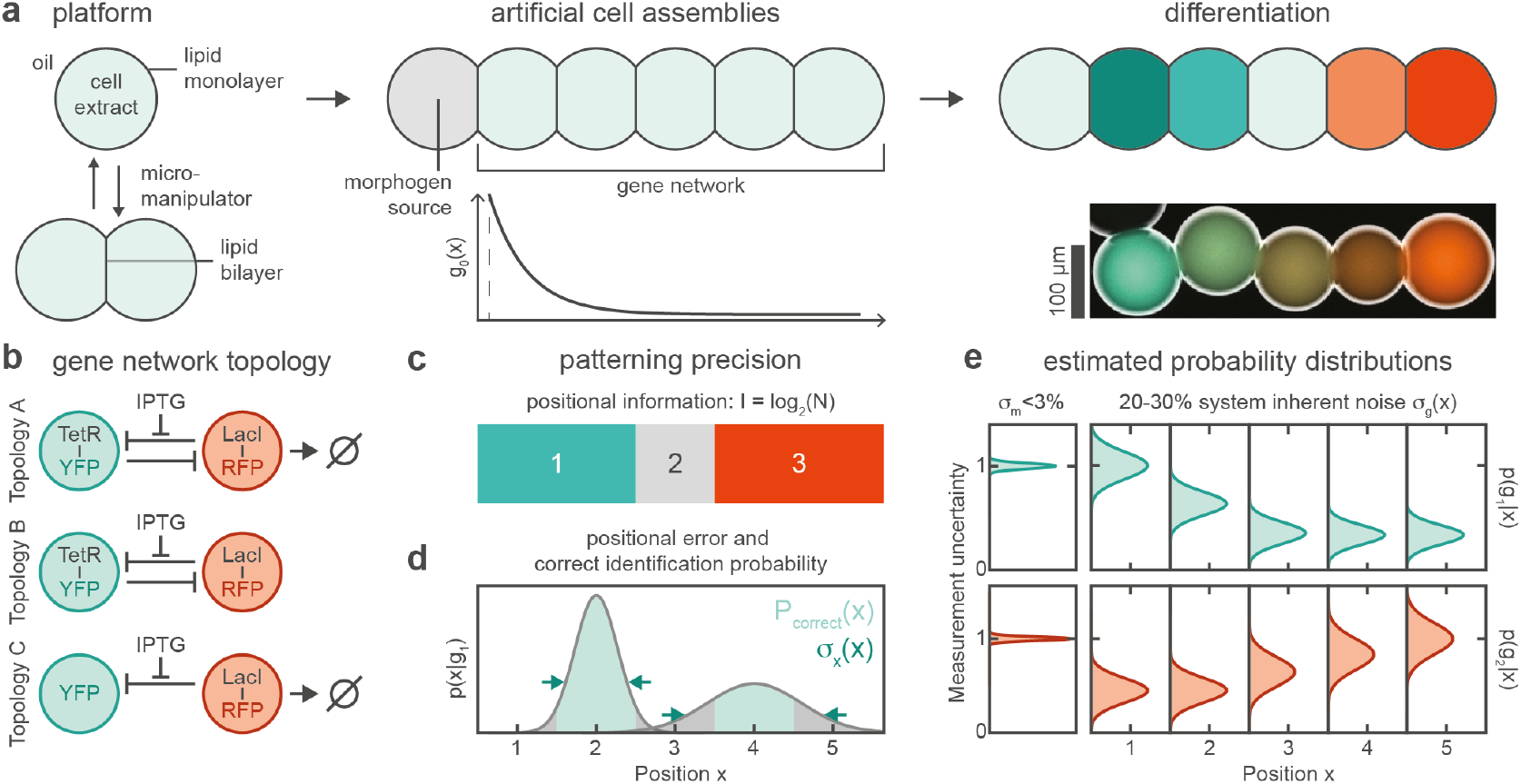
Investigation of morphogen-based differentiation in synthetic cell assemblies. The response of gene networks to morphogen gradients is used to quantify positional information and positional error, to assess the accuracy of patterning in this system. **a**, Artificial cell assemblies consist of nanoliter-sized water-in-oil droplets where oil-dispersed lipids form a monolayer at the water-oil interface. Droplets are then brought in contact using a micromanipulator to form bilayer interfaces. Assemblies consist of a droplet containing the morphogen (the inducer IPTG in our case) and serving as a source, and identical receivers containing the gene network. The morphogen diffuses from its source and forms a dynamic gradient along the main axis of the system as indicated. The induction of the gene network in a gradient of the morphogen results in a differentiation of the artificial cells (microscopy image: red and blue represent two fluorescent reporters of gene activity, cf. **SI video** for a time lapse). **b**, Three gene network topologies investigated in this work that involve the transcriptional repressors LacI (fluorescence co-expressed with RFP) and TetR (co-expressed with YFP). Topology A: mutual repression with degradation of one of the repressors, Topology B: mutual repression without degradation, Topology C: Repression of YFP by LacI-RFP with degradation of the repressor. **c**, The capability of a circuit to differentiate distinct regions in the presence of noise can be measured by the positional information. **d**, The local uncertainty of a position estimate based on a measurement of the gene expression levels can be quantified by the positional error and the deduced probability that a position estimate is correct. **e**, To estimate the positional error and positional information the expression of the tagged repressors is measured in each droplet with high accuracy for a large collection of droplet assemblies, resulting in position dependent distributions of gene expression levels.

We investigated three simple gene circuit topologies (A, B, C) for the readout of the morphogen gradient (**Fig. 1b**). Circuits A and B comprised two transcriptional repressors (LacI and TetR) mutually repressing each other’s expression, a circuit motif often found also in biological developmental circuits ^23^. The repressors were transcriptionally fused to fluorescent proteins (mScarlet-I, an RFP variant, and YFP respectively) for readout. Furthermore, in feedback circuit A, LacI was destabilized using a degradation tag ^24^. Circuit C, where LacI-RFP with degradation tag simply represses the expression of YFP, serves as a control without feedback via a second repressor.

Due to variabilities during assembly of the array, environmental fluctuations, and variations in gene expression, both morphogen gradient and gene expression levels differ from assembly to assembly. To systematically quantify the effect of these uncertainties on pattern generation, we calculated the positional information, which is defined as the mutual information 𝒥({g_*i*_}; *x*) between a set of gene expression levels {g_*i*_}, *i =* 1,…, *I*, and the spatial position *x*, i.e.,^18, 19^

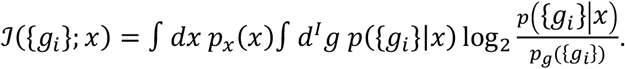

Here *p*({g_*i*_}|*x*) denotes the probability distribution of the gene expression levels for each position *x*, while *p*_*g*_({g_*i*_}) and *p*_*x*_(*x*) are the corresponding marginal distributions. Intuitively, PI measures the base-2-logarithm of the number of differentiated regions in the system (**Fig. 1c**).

We also evaluate how well the position can be determined at a specific point by quantifying the positional error (PE) *σ*_*x*_ (*x*). PE can be estimated using the Fisher information 𝒥 (*x*) = *∫ d*^*I*^ g *p*({g_*i*_}|*x*) . (*∂*_*x*_ log *p*({g_*i*_}|*x*))^2^, which provides a lower bound for the PE via the Cramer-Rao inequality 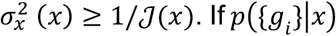 is Gaussian, PE can be estimated from the gradients in the mean and variance of the gene expression levels as

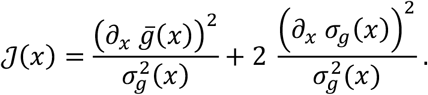

Because PE diverges in regions with flat gene expression profiles (**Supplementary Figure 10**), we instead display the probability with which the position of a droplet within the array can be inferred correctly *p*_*corr*_(*σ*_*x*_ (*x*)) ∈ [0,1] (**Fig. 1d, Supplementary Figure 10**).

In order to apply these concepts to our experimental data, we hence have to infer probability distributions *p*({g_*i*_}|*x*) for each position *x* in the assembly (**Fig. 1e**), and marginalize this distribution over *x* to obtain *p*_*g*_({g_*i*_}). *p*_*x*_ (*x*) = 1/5 in our system is uniform (**Supplementary Section 2.8, Supplementary Figure 8**).

A detailed description of our data analysis pipeline to estimate *p*({g_*i*_}|*x*) is given in **Supplementary Section 2.8**. Briefly, we analyzed large numbers (*N* = 9-37, **Supplementary Table 3**) of nominally identical linear arrays of synthetic cells for each of our three circuit topologies and different inducer concentrations. To ensure that the *measured variability* allows statements about the *positional variability* of gene expression, we developed an optimized image processing routine to correct for imaging and segmentation artifacts, and normalized the fluorescence signals with a co-encapsulated reference dye to reduce the *experimental variability* of the fluorescence measurements sufficiently (coefficient of variation C.V. < 3%, compared to 20-30% in gene expression profiles).

Next, we tracked the positions of the cells over time to obtain fluorescence traces for the YFP and RFP channels (as proxies for the gene expression levels 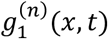 and 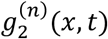for each assembly (*n =* 1,…, *N*) and at each receiver position (*x =* 1,…, 5) (**Figure 2a-c**). This resulted in experimental probability distributions for the gene expression levels at each position, i.e., *p*(g_1_|*x, t*), *p*(g_2_|*x, t*), and the joint distribution*p*({g_1_, g_2_}|*x, t*) (**Supplementary Figure 8**). We then corrected for finite sampling and binning bias as described in detail the Supplementary Information.

**Figure 2.**
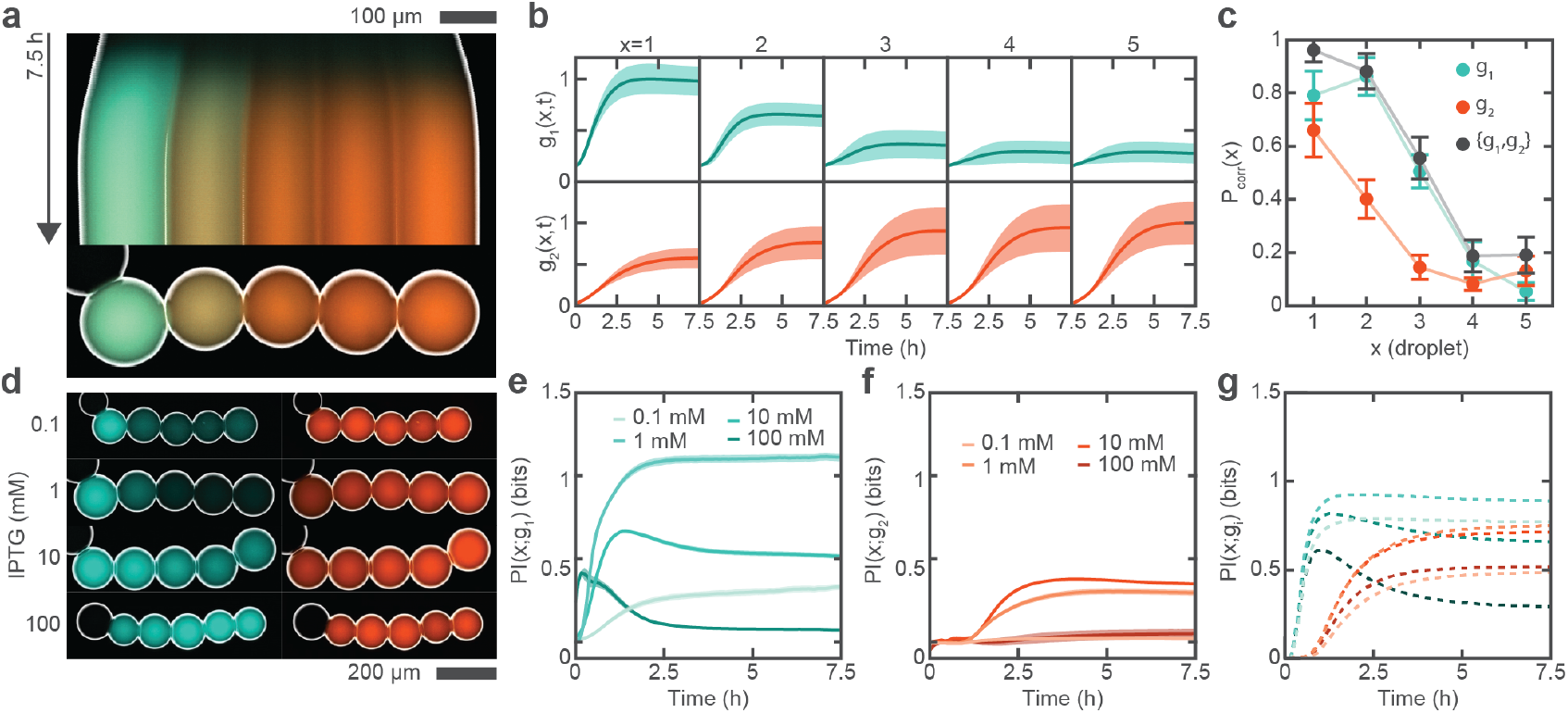
Experimental observation of gene expression gradients and determination of positional information. **a**, Kymograph of an overlay of two fluorescence channels and an inverted brightfield image showing the evolution of the two genes expressed in a single assembly for topology A at 1 mM IPTG. The IPTG sender droplet is on the left. **b**, Temporal evolution of normalized fluorescence intensity of the two genes (g_1_, TetR-YFP: blue, top, g_2_, LacI-RFP: red, bottom) for the full dataset as in **a** grouped by droplet position relative to the sender droplet. Solid lines represent the mean and shaded area represents the standard deviation for N=27 samples. **c**, Probability to correctly predict the position of a droplet based on measuring the concentration of the genes for the data in **b** after 7.5 hours. **d**, Example images for fully developed assemblies (after 7.5 hours) for varying morphogen concentrations. **e**,**f**, Temporal evolution of the estimated positional information in **e** the TetR-YFP and **f** the LacI-RFP gradient at different morphogen concentrations. The corresponding simulation **g** reproduces the experimental observations that i) the onset of PI evolution in the RFP gradient is delayed by about 1 hour and ii) that there is an optimum morphogen concentration above which PI in the YFP gradient is lost after an initial transient increase (colors correspond to **e** and **f**). Error bars in and shaded areas c,e,f are statistical uncertainties of the PI and PE estimates as described in SI Section 2.8.6 and 2.8.7.

Our key experimental observations are presented in **Figure 2**, which shows a typical kymograph for the optimized topology A at an optimal morphogen concentration of 1 mM. When the morphogen-containing compartment is connected to the assembly, a resulting morphogen gradient initially establishes rapidly in the first droplets, while increasing more slowly in the remote droplets, before assuming a relatively steady profile (**Supplementary Figure 19**). TetR-YFP expression (blue) is switched on in the nearest droplets whereas LacI-RFP (red) expression dominates in the more remote droplets.

Whether a synthetic cell can reliably “infer” its position within the array, will depend on the width of the distribution *p*({g_*i*_}|*x, t*) (**Supplementary Figure 8**), and how it compares to the distribution of the neighboring cells. From the average time traces of the full data set shown in **Fig. 2b**, it is apparent that YFP fluorescence levels differ substantially depending on the position, and the relatively small width of the distributions may allow to distinguish between several different positions. By contrast, the spatial variation of the mean RFP levels is less pronounced and the variability for each droplet position is considerable, providing less information on the position of the compartments.

In order to quantify this impression, we first calculated *p*_*corr*_ (*x*) to measure the local positional accuracy. Indeed, *p*_*corr*_ (*x*) for the first three droplets is relatively high, indicating that these droplets can, in principle, very well determine their position with respect to the organizer droplet (**Fig. 2c**). On the contrary, *p*_*corr*_ (*x*) for the two remote droplets is very low. Thus the local *p*_*corr*_ (*x*) measure indicates that the circuits are performing well for the first three droplets, but the uncertainty in the determination of the two remote droplets does not contribute to the global PI measure. Also, we observe that *p*_*corr*_ (*x*) is generally lower for the RFP gradient than for the YFP gradient and that the joint information about both gradients does not significantly improve on the information compared to the YFP gradient alone.

We next analyzed data for different, non-optimal inducer concentrations **(Fig. 2d)** and calculated the temporal evolution of positional information contained in the YFP and RFP levels (**Fig. 2e, f**). As expected, the system has zero positional information initially, but then PI-YFP rises to its maximum value (1.08-1.15 bit in the best cases across all topologies), within ≈ 2 hours (**Fig. 2e**), indicating that a distinction of more than two regions within the 5 compartments is possible. Interestingly, we observe an optimum inducer concentration, above which generation of PI-YFP is transient, peaking at around 1 hour and then decreasing again. This phenomenon can be attributed to the transient nature of the diffusing IPTG gradient, which initially only induces the first droplets, but then floods the whole system.

The positional information contained in the RFP expression levels rises with a delay of ≈ 1h and is generally less than in the YFP levels (**Fig. 2f**). The same delay is also observed in the temporal evolution of *p*_*corr*_ (*x*). The delay in PI-RFP is caused by the cascaded dynamics of LacI-RFP in the circuit: in order to reduce the LacI-RFP level in the droplets close to the IPTG source, TetR-YFP has to be produced first, LacI-RFP production has to be stopped and already present LacI-RFP has to be degraded (which only occurs in topology A). Positional information calculated from the *joint* probability distributions for both TetR-YFP and LacI-RFP gave only a slight improvement over the PI contained in the YFP levels alone (**Supplementary Figure 9**). Taken together, these results support the hypothesis that PI is transferred from TetR-YFP to LacI-RFP (and is therefore necessarily lower and delayed), rather than generated jointly.

We attempted to rationalize these results by computationally modelling the circuits’ responses to morphogen gradients (**Supplementary Section 3**), reaching good qualitative agreement of the resulting simulated PI time traces (**Fig. 2g)** with the experimental data. We also utilized computational modelling to assess the influence of key parameters on the positional information such as circuit dynamics, induction thresholds, and different types of noise. As diffusion of the morphogen through our assemblies is limited by its permeation through the lipid bilayers (**Fig. 3c, Supplementary Figures 13 and 18**), each compartment can be regarded homogeneous and positions can thus be considered discrete. In our model, we therefore only considered slow permeation of the morphogen through the interface bilayers, and calculated the circuit responses in each droplet by solving the corresponding ordinary differential equations (**Supplementary Section 3**).

**Figure 3.**
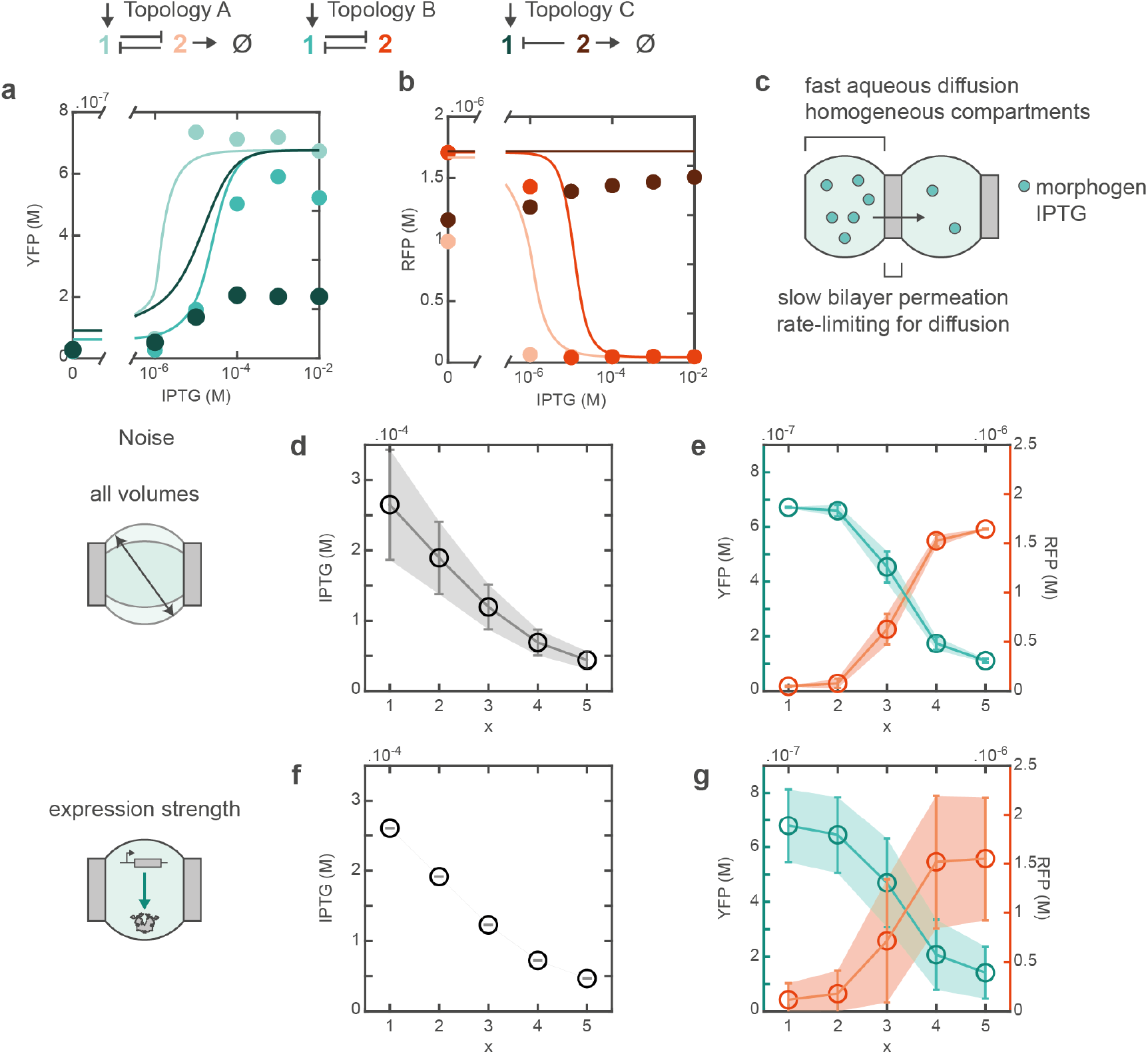
Bulk response and noise sources. **a**,**b**, Bulk titrations and simulated dose-response curves for the 3 circuit topologies for **a**, the YFP readout and **b**, the RFP readout. Simulations were performed with the same set of parameters obtained from bulk titrations of the individual nodes (Supplementary Figure 13). Hence, the change in the apparent circuit *K*_*d*_ can be purely attributed to the differences in circuit topology. **c**, Diffusion of IPTG through DIB networks is membrane limited, resulting in a dynamic morphogen gradient in a discrete space. **d-g**, Two sources of noise are considered to explain the observed variability in the gene expression gradients. First, geometrical noise, as caused for instance by variations in droplet volumes and consequently bilayer areas, leads to variability in the morphogen gradient **d**, but does not strongly propagate to a variability in the gene expression profiles **e**. On the contrary, variability in gene expression strength does not affect the morphogen gradient **f**, but leads to a variability of protein profiles **g** closely reproducing the observed variability in the gene expression profiles. All error bars are standard deviations from 500 simulations, where the droplet volumes, or gene expression strengths were drawn from a normal distribution with CV of 35 % and 20 %, respectively (**Supplementary Section 3.4**).

Relevant model parameters were measured experimentally by characterizing the circuit responses in bulk experiments, in which we varied the concentration of the inducer morphogen IPTG (**Fig. 3a, b**). With increasing amounts of IPTG, TetR and YFP production is activated in all circuits as expected. In addition, for the feedback circuits (A and B) LacI and RFP production is reduced for high [IPTG], whereas in the absence of feedback (topology C) the RFP level is unaffected by IPTG. The transfer functions of the three topologies differ as predicted by the model: the feedback circuits resulted in steeper regulation/transfer functions than the simple repression circuit (Hill coefficients of n = 2.3 and 1.4, as opposed to n = 0.76). The presence of a degradation tag (topology A) rendered the circuit responsive to lower concentrations of IPTG (Hill *K*_*d*_ = 1.51 μM, as opposed to Hill *K_d_* = 26.2 μM for topology B, **Supplementary Figure 16**). The steepness and threshold of these transfer functions affect how the circuit interprets the morphogen gradients and how much PI can be extracted.

We next used a Monte Carlo approach (**Supplementary Section 3.4**) to identify relevant noise sources and quantify their effect on the PI. We found that the two principal noise sources are the variation of droplet sizes together with their spatial arrangement, and gene expression noise. Other parameters, such as the volume of the morphogen-containing compartment, interfacial tension, or leaky protein expression did not significantly affect the system’s response (**Supplementary Figures 14 and 20**).

The overall variability of droplet volumes in the experiments is about 30-40%, and even though we balanced the osmolarity of the contents of the cells to avoid osmotic swelling or shrinking, cells still changed size and sometimes rearranged over time (**Supplementary Figure 11**). In the model, we systematically varied the volume of all droplets within and between assemblies. This resulted in some variation in the inducer concentrations in the individual droplets (**Fig. 3d**), but had only a minor effect on the final expression levels (**Fig. 3e**), and the resulting PI values were well above those measured experimentally (**Supplementary Figure 19**).

By contrast, variations in gene expression strength (with a constant inducer gradient (**Fig. 3f**)) were found to have a comparatively large impact on the variability of the expression levels (**Fig. 3g**), and predicted PI values close to those observed experimentally (**Supplementary Figure 19**). The apparent noise in gene expression strength is consistent with previous observations ^20, 25^ and could have various sources: time of manufacture, temperature variations, as well as partitioning effects, which have been observed even for large droplet volumes ^26^.

The experimental values obtained for the PI and its temporal evolution can be well understood through simulations with our model (**Fig. 4**). Both experimentally and in the model, we find a strong dependence of the PI on the “organizer” morphogen concentration, with a maximum PI for YFP of above 1 for topology A at [IPTG] = 1 mM. As expected, topology C – in which RFP expression is not regulated - has a PI of ≈ 0 bit for RFP. Important features of the kinetics of the PI levels are also well captured by our model (**Fig. 2g**), such as the initial rise in PI YFP followed with 1 hour delay by the rise in PI RFP, and an earlier peak of PI YFP for higher inducer concentrations. In contrast to our experiments, the positional information in the RFP and YFP expression levels is comparable (**Fig. 4a**, topology A), indicating that our model predicts a stronger correlation between YFP and RFP expression than observed experimentally.

**Figure 4.**
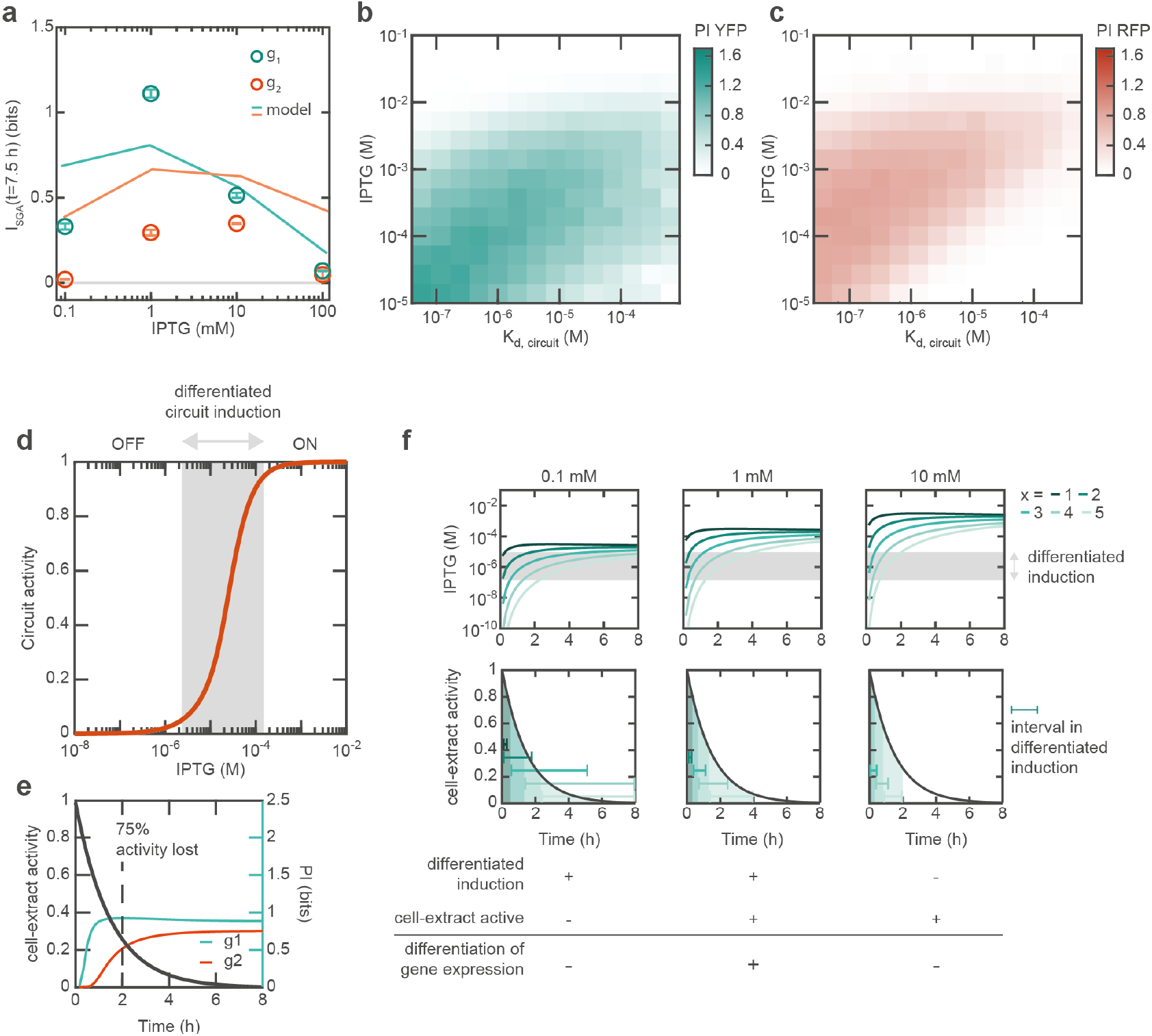
Circuit sensitivity and reaction kinetics. **a**, Comparison of the experimentally estimated and simulated final PI for circuit topology A and varying sender morphogen concentrations. The optimum PI is reached at about 1 mM IPTG. Same data and error bars as in Figure 2e-g. **b**,**c**, Simulated PI in the YFP (**b**) and RFP (**c**) gradient, respectively, for varying sender morphogen concentration and circuit *K*_*d*_. PI is maximal when the circuit *K*_*d*_ matches the morphogen gradient and the maximum PI increases with increasing sensitivity. **d**, Circuit activity (topology B, YFP) against inducer morphogen concentration, indicating three regions: circuit OFF (below 5 % activity), circuit ON (above 95 % activity), and the region where the circuit induction can be differentiated (shaded area, between 5 % and 95 % activity). **e**, Simulated temporal evolution of PI overlaid with the presumed reaction activity. **f**, The diffusion kinetics of the morphogen gradient (left panel: 0.1 mM, center panel: 1 mM, right panel: 10 mM) creates a time window during which differentiated induction of the circuit can occur. Maximizing this time window for each receiver with respect to the finite reaction lifetime allows to generate higher PI.

Our computational model further suggests that positional information is optimal for an inducer gradient matching the threshold of induction of the genetic circuit, and that higher PI can be reached for more sensitive circuits induced with correspondingly lower morphogen concentrations (**Fig. 4b, c, Fig S21)**. Indeed, differentiation of the protein expression between compartments is optimal when the morphogen profile spans the response region of the circuit, determined by the threshold and steepness of the transfer function (**Fig. 4d**). When the morphogen concentration is too low or too high (e.g. [IPTG] = 10 mM), all compartments are similarly induced and the positional information is low (**Fig. 4f**, right panel). The morphogen profile also explains the transient peak of PI for high inducer concentrations: initially, gene expression is only induced in the first droplets and the system is more differentiated, but soon all droplets are fully induced and the total PI of the system decreases.

Crucially, the cell-free expression reaction only has a finite lifetime on the order of a few hours, which we phenomenologically account for in the model with an exponential decay function (**Supplementary Section 3.2.3**). Whether a protein gradient is established depends on whether the IPTG concentration rises above the induction threshold within the lifetime of the cell-free expression system and how much activity is left at this point (**Fig. 4e**). When the morphogen profile reaches the circuit’s threshold too late (e.g. [IPTG] = 0.1 mM), most of the expression activity is lost and differentiated gene expression no longer occurs (**Fig. 4f**, left panel).

We finally studied the dynamics of the circuit in the context of more complex, 3D printed assemblies of synthetic cells (**Fig. 5**). To this end we developed a custom multipurpose bioprinting platform (**Fig. 5a, b, Supplementary Section 2.6**), which could be used to deposit simple, well-defined assemblies of emulsion droplets (**Fig. 5c**), or more complex, tissue-like assemblies (**Fig. 5d, e**) extending in three dimensions (for a detailed description of the bioprinter see the Supplementary Information).

**Figure 5.**
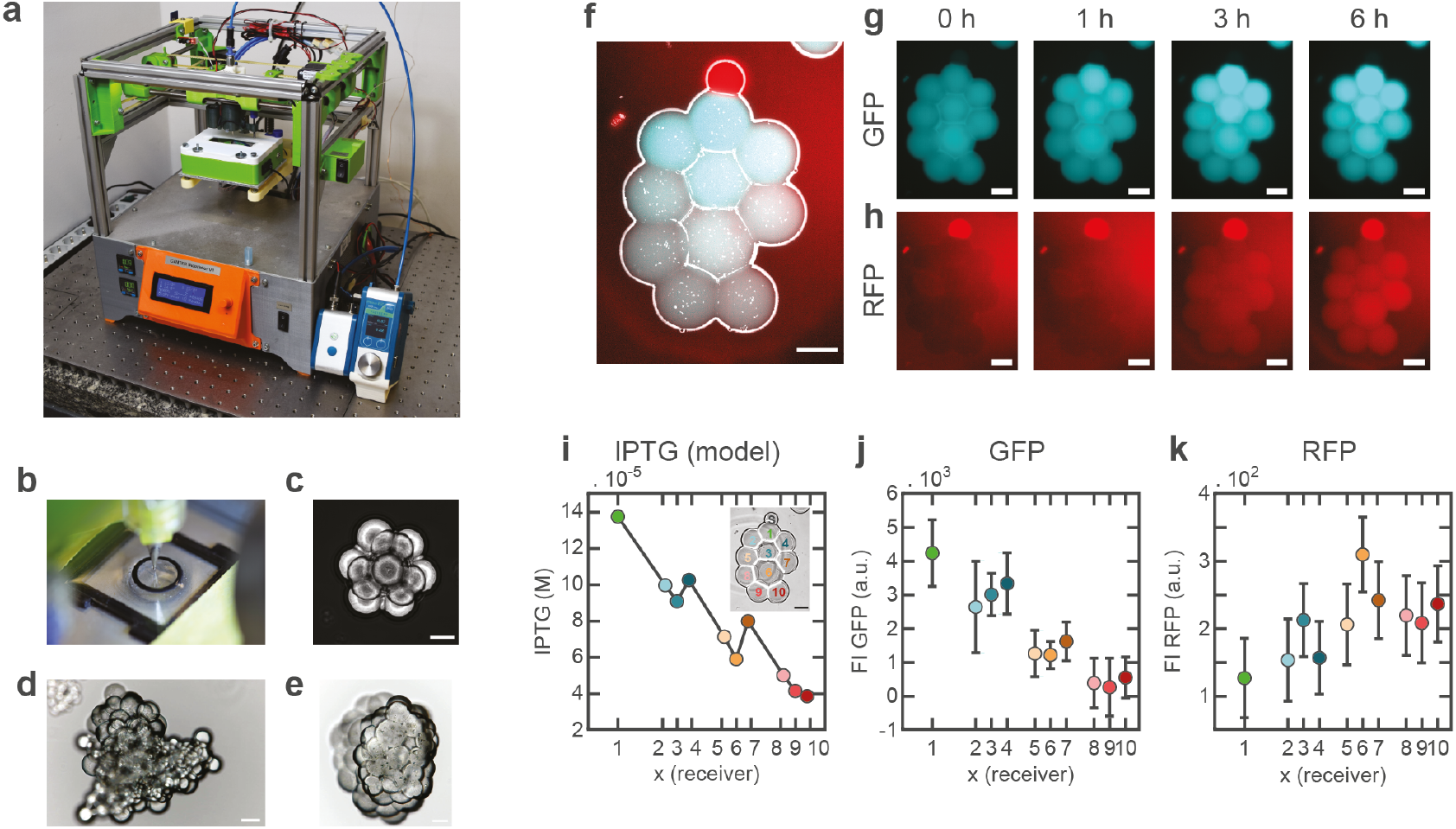
3D printing of assemblies and circuit implementation. **a**, Automated assembly of the networks is implemented with a custom-built bioprinting platform. **b**, Experiment chamber and printing nozzle. **c**, Well-defined but small 3D structures can be reproducibly printed, such as pyramid with a 7 droplets base and close-packing of the droplets. Scale bar: 100 μm. **d**,**e**, Printing of larger tissue-like assemblies. Scale bar: 100 μm. **f**, The two-node feedback gene circuit is implemented in a larger printed 2D assembly. Overlay of inverted brightfield image and fluorescence images (blue: TetR-GFP and red: LacI-RFP, sender droplet containing 1 mM IPTG at the top of the assembly is marked with a red dye). Scale bar: 100 μm. **g**,**h**, Fluorescence images of GFP reporter (**g**) and RFP reporter (**h**) of the assembly in **f** at 0, 1, 3 and 6 h. Scale bar: 100 μm. **i**, Simulated IPTG concentration at 6 hours in the assembly’s geometry. **j**,**k**, GFP and RFP fluorescence at 6 h in the assembly. Technical mean and standard deviation from data extraction are indicated by circles and error bars.

As shown in **Fig. 5f**, an “organizer cell” is initially loaded with IPTG as for the linear assemblies discussed above. Upon diffusion into the assembly, GFP (used as an alternative reporter for TetR) expression is activated in the proximal droplets (**Fig. 5g, j**), while RFP is generated in the distal droplets (**Fig. 5h, k**), clearly generating a division into two regions.

As shown in **Fig. 5i-k**, an additional structure appears to emerge in the 3D assembly, which is caused by the sphere packing geometry of the droplets. Gene expression levels vary in “shells” around the organizer cell, within which the receiver droplets approximately have the same distance from the source. This is consistent with the predicted morphogen levels in each droplet at 6 h.

As it was challenging to generate a large number of identical 3D assemblies with our current setup, a systematic analysis of the positional information in this context could not be carried out. Nevertheless, the step-like change in expression levels in the “shells” seen in **Fig. 5j** suggests that even more positional information could be extracted in the 3D context than in our 1D chains.

We have shown that the concepts of positional information and positional error – originally used for the analysis of developmental processes in biology – can be fruitfully applied also in the artificial context of synthetic cellular assemblies, which is motivated by the vision of dynamic, “living” biomaterials composed of synthetic cells that interact with their environment and are capable of context-dependent cellular specialization or differentiation. Efficient use of positional information will be important in cases where the materials have to autonomously make decisions in the presence of uncertainties generated during their production and self-assembly.

In the specific case considered here, we analyzed three simple gene circuit topologies based on a standard feedback motif containing two mutually repressing transcriptional repressors, which were encapsulated in linear assemblies of five cell-like compartments connected by permeable lipid bilayer membranes. We found that the circuits indeed responded to externally generated “morphogen” gradients, which were created by genetic inducers emanating from an “organizer” cell. From this gradient, in the best case the circuits could derive PI of ≈ 1.2 bits, which allows distinction of two to three regions within the five-cell assembly. In order to infer the position of all five cells, a correspondingly higher PI of log^2^ 5 = 2.3 bits would be required. The use of the PI and PE concept also allowed us to precisely quantify which of the circuits could best transfer information from one morphogen layer to the next (from TetR to LacI), which otherwise would have been based on rather vague visual impressions.

Several factors presently limit the extraction of positional information from the gradient – next to the inherent variability in gene expression itself, the temporal evolution of the gradient and of the cell-free expression system itself play a role. Importantly, in our closed system containing a finite reservoir of inducer molecules acting as morphogens, the gradient steadily evolves during the “differentiation” process (**Supplementary Figure 19**). As observed experimentally, the inducer concentration has to be chosen properly to observe any spatial variation at all. Our computational model shows how the interplay of circuit response (characterized by the induction threshold *K*_*d*_) and diffusion dynamics affects the total positional information (**Fig. 4 and Supplementary Figure 21**). If the concentration is too high (**Fig. 4f**), the inducers quickly rise above the induction threshold in all droplets, preventing any spatial differentiation. If it is too low, the concentrations are not sufficient to elicit any response. The spatiotemporal evolution of the morphogen is then sampled by the circuits for a finite amount of time. Cell-extract activity decays exponentially over time (**Fig. 4e**), and after 2 hours 75 % of the protein expression activity is lost. Thus, if diffusion is too slow, the inducers will reach some of the droplets too late (i.e., after their “death”, **Fig. 4f** left-most panel). In the ideal case, the inducer concentration in all droplets spans the sensitive transition region of the circuit response function for longer periods of time, allowing for a differentiated induction of the response in each droplet (**Fig. 4f** center panel, [IPTG] = 1 mM).

The deterioration of the cell-free system and relatively slow gene expression dynamics also affect the transfer of positional information from one transcription factor to another (i.e., from TetR to LacI in our case). Our system is not able to complete a full cycle of the feedback circuit employed, and therefore the gradient information extracted from the IPTG gradient by TetR-YFP cannot be transferred to the LacI expression level, which it influences.

This situation is not unlike that of developing organisms, where cell fate must be determined in the time period during which morphogen gradients are stable and provide high positional information. In the case of the vertebrate neural tube for example, the combined antiparallel gradients of morphogenetic proteins can only allow precise patterning during 10 hours before they diverge ^17^. Layers of patterning circuits, wherein a morphogen gradient induces the differentiated expression of a protein which itself becomes a morphogen for other proteins allows for robust, step-wise patterning of an organism. Contrary to our system, production-diffusion-degradation mechanisms often stabilize morphogen gradients in living systems. In the case of *Drosophila*, in the first 90 min of embryonic development a roughly exponential gradient of Bcd/Bcd-mRNA is established along the anterior-posterior axis, which stays stable for ≈ 1 hour (between cell cycles 10 and 14), during which the spatial differentiation of the next layer of morphogens (caudal and hunchback) is achieved ^16^. The stability of this gradient has been recognized as one of the key factors for the astonishingly high positional information extracted in this biological system.

In addition to a stabilized morphogen gradient, an efficient use of the positional information contained in the gradient would require a cell-free expression system with an extended lifetime. To this end, replenishment of its resources via some sort of external supply system would be necessary, possibly coupled to metabolic activity (such as ATP production/regeneration, or even self-regeneration). The finite time window generated by gradient stability and lifetime of the expression system could be used more efficiently by speeding up the regulatory processes involved in the differentiation process. Realization of these improvements could be quite challenging in the context of the closed emulsion-based compartments studied here. Open bioreactors (e.g., permeable structures immersed in “broth” or combined with microfluidic supplies), could thus be promising for “self-differentiating” biomaterials, in which multiple layers of developmental genes would carry out a synthetic morphogenetic program. Further improvements in 3D printing technologies will also facilitate the automated generation of extended, tissue-like structures with an artificial “vasculature”, which would help to keep these “living” biomaterials out of equilibrium over extended periods of time.

## Author contributions

A.D. purified LacI and TetR, conducted preliminary experiments and analysis, conducted the droplet experiments, conceptualized the project. L.A. analyzed the droplet experiments, developed the analysis procedure, and calculated the positional information. I.S. purified LacI and TetR and determined kinetics and thermodynamic constants. I.S., F.R. and A.D. cloned the plasmids. F.R. conducted preliminary experiments. B. K., S. S., N. G. and H. C.-S. conceptualized and developed the multipurpose bioprinting platform and relevant components for this specific experiment. F.C.S co-conceptualized the project, and contributed to the analysis. A.D., L.A., and F.C.S co-wrote the paper.

## Acknowledgements

We gratefully acknowledge financial support for this project by the European Research Council (grant agreement no. 694410 - AEDNA). B. K., S. S., N. G. and H. C.-S acknowledge financial support from the Bavarian State Ministry for Science and Arts thought the research focus “Center for Applied Tissue Engineering and Regenerative Medicine (CANTER)”. We would like to thank Elisabeth Falgenhauer for purifying LacI, TetR and mTurquoise2, Stefanie Sudhop for co-developing the bioprinting platform, and Ulrich Gerland for useful discussions.

## Notes

### Competing Interest Statement

The authors have declared no competing interest.

