## Supplementary Information for "Synthetic cell-based materials extract positional information from morphogen gradients"

Aurore Dupin[1]*, Lukas Aufinger[1]*, Igor Styazhkin[1], Florian Rothfischer[1], Benedikt Kaufmann[2,3], Sascha Schwarz[2,3,4], Nikolas Galensowske[3], Hauke Clausen-Schaumann[2,3], Friedrich C. Simmel[1#]

[1] *Physics Department – E14, TU Munich, 85748 Garching, Germany*

[2]*Center for NanoScience – CeNS, Schellingstraße 4, 80799 Munich, Germany*

[3]*Center for Applied Tissue Engineering and Regenerative Medicine – CANTER, Munich University of Applied Sciences, Lothstrasse 34, 80335 Munich, Germany*

[4]*Chair of Applied Mechanics, TU Munich, 85748 Garching, Germany*

[#]

2.8.7 Calculation of Positional Error ... 23
3 Models ... 25
3.1 Diffusion in droplet assemblies ... 25
3.1.1 General diffusion equation ... 25
3.1.2 Geometrical parameters ... 25
3.2 Protein expression, repression and induction ... 27
3.2.1 Protein expression ... 27
3.2.2 Repression and induction ... 28
3.2.3 Protein degradation and life-time of cell-extract ... 28
3.2.4 Protein and fluorescence intensity ... 29
3.3 Reaction-diffusion model ... 29
3.4 Noise ... 30
4 Supplementary experiments ... 30
4.1 Toxicity of reference dye ... 30
4.2 Fitting of model parameters ... 31
4.2.1 Repression constants ... 31
4.2.2 Permeability of bilayers to IPTG ... 31
4.3 Simulations of bulk response ... 32
4.4 Characterization in droplets ... 35
4.5 Simulations of reaction-diffusion circuit ... 36
5 Sequences ... 40
6 References ... 43

### 1 Materials

Sigma-Aldrich, Germany:
isopropyl b-D-1-thiogalactoside IPTG #I6758, hexadecane #296317, silicone oil AR 20 #10836, $\alpha$-Hemolysin #H9395, anhydrotetracycline (aTc) #37919, Atto488 #41051, lysozyme #L6876.

Carl Roth, Germany:
LB Medium #X968, glycerol #3783, nuclease-free water (nf H2O) #T143, chloroform #3313, ethanol #P076.

Avanti Polar Lipids (USA):

DPhPC (4ME 16:0 PC) #850356, DOPC (18:1 ($\Delta$9-Cis) PC) #850375, DOPG (18:1 ($\Delta$9-Cis) PG) #840475, cholesterol (20$\alpha$-hydroxycholesterol) #700156.

For DNA, single-stranded oligos were purchased from Biomers (Germany) or Eurofins (Germany), and double-stranded gBlocks were purchased from IDT (USA).

LacI, TetR and mTurquoise2 were purified in our lab according to standard His-tag nickel purification protocols (with the following specific buffers for LacI[1], TetR[2]). For mTurquoise2, the following buffers were used: lysis and wash buffer 50 mM Tris, 500 mM NaCl, 25 mM imidazole, 1 mM DTT, pH 8.0, elution buffer 50 mM Tris, 500 mM NaCl, 250 mM imidazole, 1 mM DTT, pH 8.0, storage buffer 50 mM Tris, 100 mM NaCl, 2 % v/v DMSO, 15 % glycerol, 1 mM DTT, pH 7.5).

All other chemicals were purchased from Sigma-Aldrich, Germany.

### 2 Methods

#### 2.1 Plasmid cloning and purification

All plasmids were cloned using standard strategies and enzymes from New-England Biolabs (NEB, USA). We alternatively used digestion/ligation, Golden Gate assembly[3] or PCR with 5'-phosphated primers followed by ligation.

The plasmids were transformed into the *E. coli* bacterial strain Turbo (NEB, USA, #C2984I). Constructs were cloned in the plasmid backbones pSB1A3 and pSB1C3. Bacteria were stored in glycerol stocks: an overnight culture in LB medium was mixed with glycerol to a final concentration of 25 % v/v glycerol and kept at – 80 °C. To ensure that the glycerol stocks contained a monoclonal population with the correct plasmid, an overnight culture was grown in LB medium from the glycerol stock, Mini-prepped (QIAprep Spin Miniprep Kit, Qiagen, Netherlands) and sequenced (Sanger sequencing, LightRun, GATC Biotech, Germany). Plasmids were purified prior to use in cell-extract by Midiprep (NucleoBond Xtra Midi, Macherey-Nagel, Germany).

#### 2.2 Cell extract preparation

The E. coli cell extract was prepared according to the protocol by Sun *et al.*[4]. Shortly, a mid-log phase culture of BL21(DE3) Rosetta2 was spun, washed, resuspended at 1 g/mL and incubated with 1 mg/mL lysozyme on ice for 30 min. It was then lysed by sonication in 4 mL aliquots at 30 kHz, 10% amplitude, 20 cycles, 10 s/cycle. The extract was incubated at 37 °C for 80 min to allow the digestion of genomic DNA, and was then dialyzed for 3 h at 4 °C with a cut-off of 10 kDa (Slide-A-Lyzer Dialysis Cassettes, Thermo Fisher Scientific, USA). Protein concentration was estimated to be 30 mg/mL with a BCA essay (Thermo Fisher Scientific, USA, #23225). In the buffer, instead of 3-phosphoglyceric acid (3-PGA), phosphoenolpyruvate (PEP) was utilized as an energy source[5]. The buffer was composed of 50 mM HEPES pH 8, 1.5 mM ATP and GTP, 0.9 mM CTP and UTP, 0.2 mg.ml$^{-1}$ tRNA, 0.26 mM coenzyme A, 0.33 mM NAD, 0.75 mM cAMP, 68 µM folinic acid, 1 mM spermidine, 30 mM PEP, 1.25 mM leucine, 1.5 mM other amino acids, 1.5 mM DTT, 3.5 % PEG-8000, 80 mM K-glutamate and 4 mM Mg-glutamate. Buffer and extract were flash-frozen in liquid nitrogen, stored at – 80 °C and thawed on ice prior to usage. A cell-free reaction was prepared by mixing 33 % v/v cell extract with 42 % v/v buffer and 25 % v/v DNA, inducers, and other additives. Reactions were conducted at 29 °C.

#### 2.3 Experimental conditions

All experiments described in this paper were performed in cell extract. 1 nM of the TetR-YFP plasmid and 3 nM of the LacI-mScarletI plasmid were used. The concentration of IPTG was as indicated. 265 nM of mTq2 was used in droplet experiments to normalize the fluorescence to droplet volume (see Supplementary Section 2.8.2 and Supplementary Figure 12). The plasmids correspond to the topologies or to the Supplementary Figure experiments as shown in Supplementary Table 1**.**

**Supplementary Table 1 | Plasmids and experimental conditions.**

| Conditions | TetR-YFP plasmid | LacI-mScarletI plasmid |
|---|---|---|
| Topology A | pSB1C3-AD052 | pSB1A3-AD064 |
| Topology B | pSB1C3-AD052 | pSB1A3-AD084 |
| Topology C | pSB1C3-AD085 | pSB1A3-AD064 |
| Supplementary Figure 17, 18 | pSB1C3-pLacO-TetR-GFP | pSB1A3-pTetO-LacI-mCherry |

#### 2.4 Bulk experiments

Fluorescence measurements in bulk were conducted at 29 °C in a plate reader (FLUOstar Omega or CLARIOstar, BMG Labtech, Germany). 15 μL samples were pipetted in a 384-well plate (pureGrade, Brand, #781622), the plate was sealed with an optically transparent film (Microseal 'B', Bio-Rad, Germany), and centrifuged 30 s at 700 rcf.

#### 2.5 Manual droplet assembly experiments

Droplet experiments were carried out as described in Dupin & Simmel, 2019[6]. Shortly, water-in-oil emulsions were created with a pressure pump and a micromanipulator. The lipid-oil mixture consisted of 0.25 mM cholesterol, 0.25 mM DOPG, 4 mM DOPC and 0.5 mM DPhPC in 1:1 hexadecane:AR20. The lipid and cholesterol powders were dissolved in chloroform, mixed in a glass vial, evaporated under a nitrogen stream and dried under vacuum, after which the lipid film was resuspended with the oil mix and vortexed. The chambers consisted of rubber O-rings glued with an epoxy glue onto glass slides. After being washed with soap, rinsed with ethanol and double-distilled water and dried at 90 °C, the chamber was filled with 65 μL of the lipid-oil mixture. Solutions were pipetted into heat-pulled glass capillaries, which were then fixed onto a house-built micromanipulator and connected to a microinjector pump. Pressure injections were applied to create droplets of around 130 μm in diameter, between 0.5 and 1 nL volume. Droplets were incubated in the oil for 15 min on average to allow for a monolayer of lipids to assemble around them, and were moved together with the micromanipulator so that they spontaneously formed bilayers. The chamber was closed with a glass slide to limit evaporation of the droplets.

#### 2.6 Printed assembly experiments

Solutions in cell-extract, lipid-oil mixture and experiment chambers were prepared as for manual assembly experiments.

##### 2.6.1 G-code generation

The G-codes required for printing the 3D assemblies were generated manually by programming the individual coordinates required for droplet placement. Supplementary Table 2 displays an example G-code file.

**Supplementary Table 2 | Exemplary G-code for the printing of the pyramid in Supplementary Figure 1a-c.**

| Command | Comment |
|---|---|
| Set-up of the printer | |
| G90 | positioning mode set to "absolute" |
| M83 | mode of E-axis set to "relative" |
| M106 S0 | fan speed set to "0" |
| M104 S0 T0 | temperature of extruder 0 set to "0" |
| M104 S0 T1 | temperature of extruder 1 set to "0" |
| G1 Z15 F5000 | send down petri dish (speed 5000 mm/min) |
| G28 X Y | home X- and Y-axis |
| G1 X60 Y3.5 F5000 | go to autobed-leveling position |
| G28 Z | home Z-axis |
| G92 X30 Y1 Z0 | determine new zero-positions |
| M102 | extruder 0 turned on |
| G1 E-1.0000 F60000 | valve closed by negative extrusion |
| M103 | extruders turned off |
| G1 Z12.900 F300 | Z-axis moved to starting position |
| G1 Z10 F5000 | Z-axis moved to starting position |
| T1 | extruder 0 selected for further commands |
| Printing of the first layer of the assembly | |
| G1 X7.5 Y25 F2000 | printhead moved to required X,Y-position |
| G1 Z0.1 F2000 | printhead sent down into oil/lipid reservoir |
| G1 E1.0000 F200000 | extrusion |
| G1 E-2 F2000 | valve closed by negative extrusion |
| G4 S1 | pause 1 s |
| G1 Z7 F2000 | printhead sent up |
| G4 S30 | pause 30 s |
| G1 X7.5 Y24.95 F2000 | printing at a new position |
| G1 Z0.1 F2000 | |
| G1 E1.0000 F200000 | |
| G1 E-2 F2000 | |
| G4 S1 | pausing |
| G1 Z7 F2000 | |
| G4 S30 | |
| G1 X7.51 Y24.95 F2000 | printing at a new position |
| G1 Z0.1 F2000 | |
| G1 E1.0000 F200000 | |
| G1 E-2 F2000 | |
| G4 S1 | pausing |
| G1 Z7 F2000 | |
| G4 S30 | |
| G1 X7.49 Y24.95 F2000 | printing at a new position |
| G1 Z0.1 F2000 | |
| G1 E1.0000 F200000 | |
| G1 E-2 F2000 | |
| G4 S1 | pausing |
| G1 Z7 F2000 | |
| G4 S30 | |

| | |
|---|---|
| G1 X7.548 Y24.984 F2000<br>G1 Z0.1 F2000<br>G1 E1.0000 F200000<br>G1 E-2 F2000<br><br>G4 S1<br>G1 Z7 F2000<br>G4 S30<br><br>G1 X7.452 Y24.984 F2000<br>G1 Z0.1 F2000<br>G1 E1.0000 F200000<br>G1 E-2 F2000<br><br>G4 S1<br>G1 Z7 F2000<br>G4 S30<br><br>G1 X7.5 Y25.05 F2000<br>G1 Z0.1 F2000<br>G1 E1.0000 F200000<br>G1 E-2 F2000<br><br>G4 S1<br>G1 Z7 F2000<br>G4 S30 | printing at a new position<br><br><br><br>pausing<br><br><br><br><br>printing at a new position<br><br><br><br><br>pausing<br><br><br><br>printing at a new position<br><br><br><br><br>pausing |
| Printing of the second layer of the assembly | |
| G1 X7.525 Y24.957 F2000<br>G1 Z0.1 F2000<br>G1 E1.0000 F200000<br>G1 E-2 F2000<br><br>G4 S1<br>G1 Z7 F2000<br>G4 S30<br><br>G1 X7.525 Y25.043 F2000<br>G1 Z0.1 F2000<br>G1 E1.0000 F200000<br>G1 E-2 F2000<br><br>G4 S1<br>G1 Z7 F2000<br>G4 S30<br><br>G1 X7.45 Y25 F2000<br>G1 Z0.1 F2000<br>G1 E1.0000 F200000<br>G1 E-2 F2000<br><br>G4 S1<br>G1 Z7 F2000<br>G4 S30 | printing at a new position<br><br><br><br><br>pausing<br><br><br><br>printing at a new position<br><br><br><br><br><br>pausing<br><br><br><br>printing at a new position<br><br><br><br><br>pausing |
| Printing of the third layer of the assembly | |
| G1 X7.5 Y25 F2000<br>G1 Z0.1 F2000 | printing at a new position |

| | |
|---|---|
| G1 E1.0000 F200000<br>G1 E-2 F2000<br><br>G4 S1<br>G1 Z7 F2000<br>G4 S30 | <br><br><br>pausing |
| Closing sequence | |
| M102<br>G1 E-1.0000 F60000<br>M103<br>G1 Z30 F5000<br>G1 X0 Y0 F5000<br>M104 S0<br>M140 S0<br>M84 | extruder 0 turned on<br>valve closed by negative extrusion<br>extruders shut down<br>printing plate moved down<br>printhead moved to starting position<br>turn off extruders temperature regulation<br>turn off beds temperature regulation<br>disable motor |

##### 2.6.2 Automated droplet assembly

In comparison to the manually created circuits using the micromanipulator system, the 3D arrays (Supplementary Figure 1) were generated using a self-developed modular bioprinting platform (Supplementary Figure 2b). The experiment chamber was filled with 65 µl of lipid oil equilibrated to room temperature (22 °C) and the print head fitted with a Micron-S nozzle (VIEWEG, Germany) with an inner diameter of 60 µm (Supplementary Figure 2c). Pneumatic extrusion was realized using the Flow EZprecisions pressure pump (Fluigent SA, France) which was actuated by the printer software. In case of an extrusion event, the printer signals to the TTL receiver (LineUp LINK, Fluigent SA) controlling the pump, and the signal is then interpreted by the Microfluidics Automation Tool (Fluigent SA, France) using a custom code for pump actuation allowing the adjustment of pulse pressure and duration.

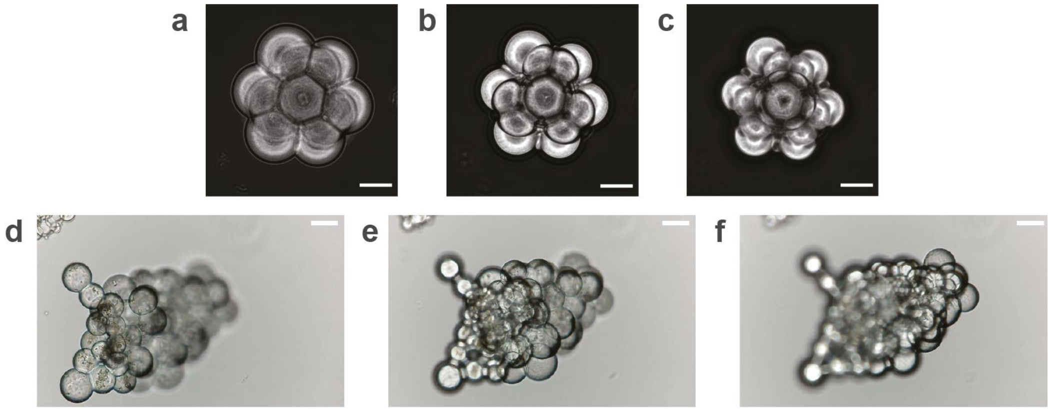

**Supplementary Figure 1 | Printed assemblies. a-c,** bright field microscopy image of a 3D-printed liposome-“pyramid” with the microscope focus on the lower (**a**), middle (**b**) and higher (**c**) layers. Micrographs were acquired with the Axio Observer.Z1 (Zeiss, Germany). Scale bar: 100 mm. **d-f,** large 3D-printed tissue-like structure at different foci: lower (**d**), middle (**e**) and higher (**f**). Photographs acquired with a modified Logitech webcam. Scale bar: 100 µm.

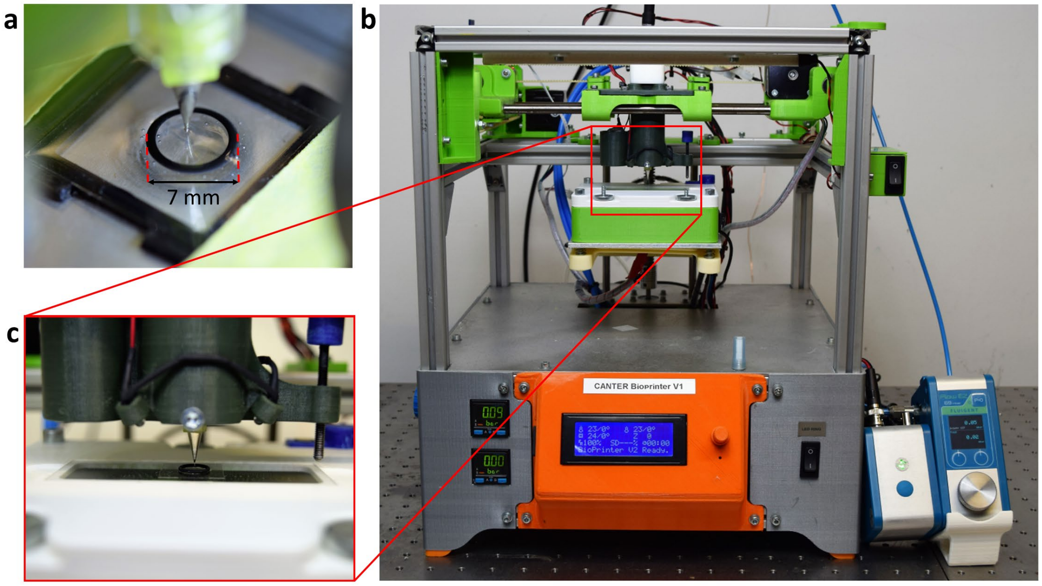

**Supplementary Figure 2 | Overview of the 3D-printer set-up for the automated assembly of droplet networks. a,** Filled experiment chamber with immersed 60 µm nozzle during droplet assembly. **b,** Overall set-up of the bioprinting platform including the print head with transmitted light microscope imaging module integrated into the z-stage and precision pump in the lower right. **c,** Close-up side view of the print head and experiment chamber.

Immediately prior to printing, the nozzle was filled manually with 94.5 µl of cell extract using a pipette. Next, the exact position of the nozzle tip was calibrated using a force sensing resistor (FSR 400, Interlink Electronics, USA). Droplet size was calibrated using the integrated microscopic imaging module of the 3D-printer. If necessary, droplet size can be adjusted by changing pulse pressure and duration of the pump setup or varying the depth of the nozzle´s intrusion into the lipid/oil bath. The nozzle is positioned along the x- and y-axis at the desired position and then immersed approximately 100 µm in the lipid reservoir and kept at this position for 1 s to allow vibrations to settle (Supplementary Figure 3a). By applying a pneumatic pulse, a drop of cell extract is extruded which remains at the tip of the nozzle due to surface tension (Supplementary Figure 3c). To overcome the surface tension, the nozzle is raised and withdrawn from the reservoir. At the liquid-air interface, the droplet is detached from the nozzle (Supplementary Figure 3d). While the droplet sinks to the bottom of the experiment chamber, the printer movement is paused (Supplementary Figure 3e, f). It is crucial that the droplet is covered by a stable lipid monolayer before it comes into contact with the existing printed droplets, to prevent it from fusing with them. The kinetics of assembly of such a lipid monolayer depends on several factors, including the droplet's surface area. In our experiments, we found that the time necessary for the droplet to sink from the oil-air interface to the existing droplet structure (approx. 1-5 s) was sufficient to prevent fusion of the droplets.

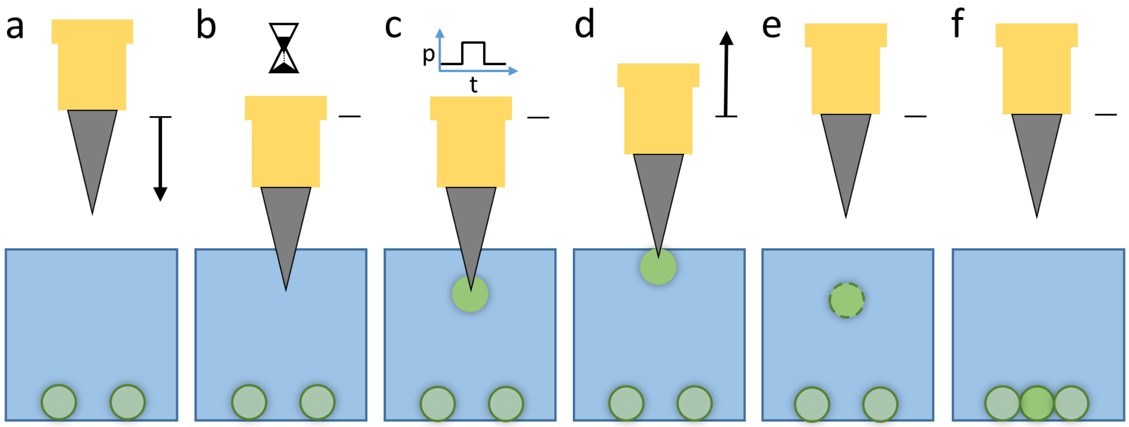

**Supplementary Figure 3 | Automated droplet generation process.** The nozzle is moved to the desired x-y-position and immersed in the lipid reservoir (**a**) and kept at this position for 1 s (**b**). By applying a pneumatic pulse, a drop of cell extract is extruded, which adheres to the tip of the nozzle, caused by surface tension (**c**). The nozzle is then withdrawn from the reservoir and the droplet is detached at the liquid-air interface (**d**). As the droplet sinks towards the substrate an intact lipid membrane is formed which stabilizes the droplet (**e, f**).

An alternative mode of printing using constant pressure (Supplementary Figure 3, skipping steps b and c) gave similar results. Here, the nozzle was dipped into the oil bath, the cell extract was extruded and the nozzle immediately lifted again to release the droplet.

##### 2.6.3 Limitations

The yield of correct prints, where accurate positioning of all droplets was achieved, was 12.5 % for a 7-droplet structure and 10 % for an 11-droplets structure (data not shown). This yield is currently limited by a number of interrelated factors. The process requires sufficient time for the lipid layer formation and nozzle movement for the droplet detachment. Vibrations and liquid currents introduced by the movement cause an erratic displacement of the droplets from their target positions, as they sink to the bottom of the experimental chamber, thus x and y coordinates of the droplets cannot be precisely controlled.

#### 2.7 Image Acquisition

The fluorescence of the networks was recorded with an inverted fluorescence microscope Eclipse Ti2 (Nikon, Japan), equipped with a SOLA light engine (Lumencor, USA) for excitation, camera Neo5.5 (Andor, Northern Ireland, UK), a 10x CFI P-Apo objective (NA 0.45, Nikon, Japan), temperature-controlled incubation chamber (Okolab, Italy), and acquisition software NIS-Elements AR from Nikon (Japan). The images were further analyzed with ImageJ and MATLAB_R2015b. The CFP channel was acquired with an exposure time of 1 s, with excitation filter 416-440 nm, dichroic mirror 458 nm, and emission filter 467-499 nm. The YFP channel was acquired with an exposure time of 500 ms, with excitation filter 479-495 nm, dichroic mirror 515 nm, and emission filter 510-540 nm. The RFP channel was acquired with an exposure time of 1 s, with excitation filter 542-576 nm, dichroic mirror 585 nm, and emission filter 617-633 nm.

For Supplementary Figure 17 and Supplementary Figure 18, the fluorescence of the assemblies was recorded with an inverted fluorescence microscope IX-71 (Olympus, Japan), with excitation LEDs, filters and dichroic mirrors from Thorlabs (USA), camera LucaEM from Andor (Northern Ireland, UK), objectives from Olympus (Japan), heating plate from Tokai Hit (Japan), and acquisition software MicroManager 1.4.16. A 10x objective was used and a bin of 2, the GFP channel was acquired with an exposure time of 1 s, with LED emission @ 470 nm, excitation filter 450-490 nm, dichroic mirror 510 nm and emission filter 518-545 nm, the RFP channel was acquired with an exposure time of 3 s, with LED 530 nm, excitation filter 530-550 nm, dichroic mirror 576 nm and for emission, a long pass filter above 590 nm.

#### 2.8 Data Analysis

We now analyzed these videos to better understand which parameters influence the quality of the developing gradients. To this end, two informative quantities that have been applied to the analysis of natural patterning systems are the positional error (PE) and the positional information (PI) in the droplet network[8–12]. The positional error $\sigma_x(x)$ at position $x$ within a droplet network can be understood as the precision with which the position $x$ of a droplet within the network can be estimated by the genetic circuit it contains, typically by 'measuring' the local concentration of potentially multiple morphogens $g_i$, where $i = 1, \ldots, I$. The positional information $\mathcal{I}\ (x; \{g_i\})$ is an information theoretical measure in units of bits and can be interpreted as the base-2 logarithm of the number of regions $\chi$ that can be globally distinguished by the genetic circuit, i.e., $\mathcal{I} = \log_2 \chi$.

In our experimental setup we use a maximum of two morphogens, TetR/YFP and LacI/RFP, hence $I = 2$. In a droplet-interface-bilayer (DIB) network we can assume that diffusive mixing within one droplet is fast (protein: $t = l^2/2D \approx 200\ \mu\mathrm{m}^2/(2 \cdot 100\ \mu\mathrm{m}^2\mathrm{s}^{-1}) = 200\ \mathrm{s}$), while the diffusion of IPTG through the network is limited by the permeation through the bilayer (Supplementary Sections 3.1.1 and 4.2.2). We therefore assume a discretized space with one unit length defined as one droplet (i.e., one bilayer to pass) and a total length of $X = 5$. This means that the theoretical maximum for the positional information in our setup is $\mathcal{I} = \log_2 5 \approx 2.32$. Similarly, a positional error of $\sigma_x(2) = 0.5$ droplets means that, for the measured morphogen concentration, the droplet is located at position 2 with a probability of about 68%. Both PE and PI quantify how well positions can be inferred by a given genetic circuit with respect to variations in morphogen concentration. Hence, we first need to estimate these variations, represented as a joint probability distribution function (pdf) $p(\{g_i\}; x)$, by measuring $g_i$ for $N$ distinct samples.

Experimentally, we can infer the morphogen concentration in a droplet by measuring the reporter fluorescence, which is proportional to the gene expression level $f \propto g$. The experimentally observed variability $\sigma_f$ , however, is a combination of the actual morphogen variability $\sigma_g$ and measurement uncertainties $\sigma_m$, i.e., $\sigma_f^2 = \sigma_g^2 + \sigma_m^2$. In our setup, the intrinsic variability $\sigma_g$ is either input noise that the genetic circuit 'sees' as a consequence of variations in droplet or bilayer size resulting in variations in the IPTG input gradient, or output noise produced by variations in gene expression strength across droplets, as discussed in the text and in Supplementary Section 3.4. In contrast, the measurement uncertainty $\sigma_m$ is related to image acquisition and analysis, i.e., it is extrinsic variability that is irrelevant to the genetic network or this analysis. In order to obtain a meaningful estimate of $\sigma_g$, it is therefore necessary to reduce the measurement noise as much as possible so that $\sigma_m/\sigma_f \ll 1$ (typically at least $< 0.2$) and hence $\sigma_g \approx \sigma_f$[10]. In the following we first use a calibration data set to estimate $\sigma_m$ and test alternative methods to reduce $\sigma_m$ from an initial 35% to about 2%, compared to a typical $\sigma_f$ of about 15-30% observed in gene expression experiments (hence at least $\sigma_m/\sigma_f < 0.15$).

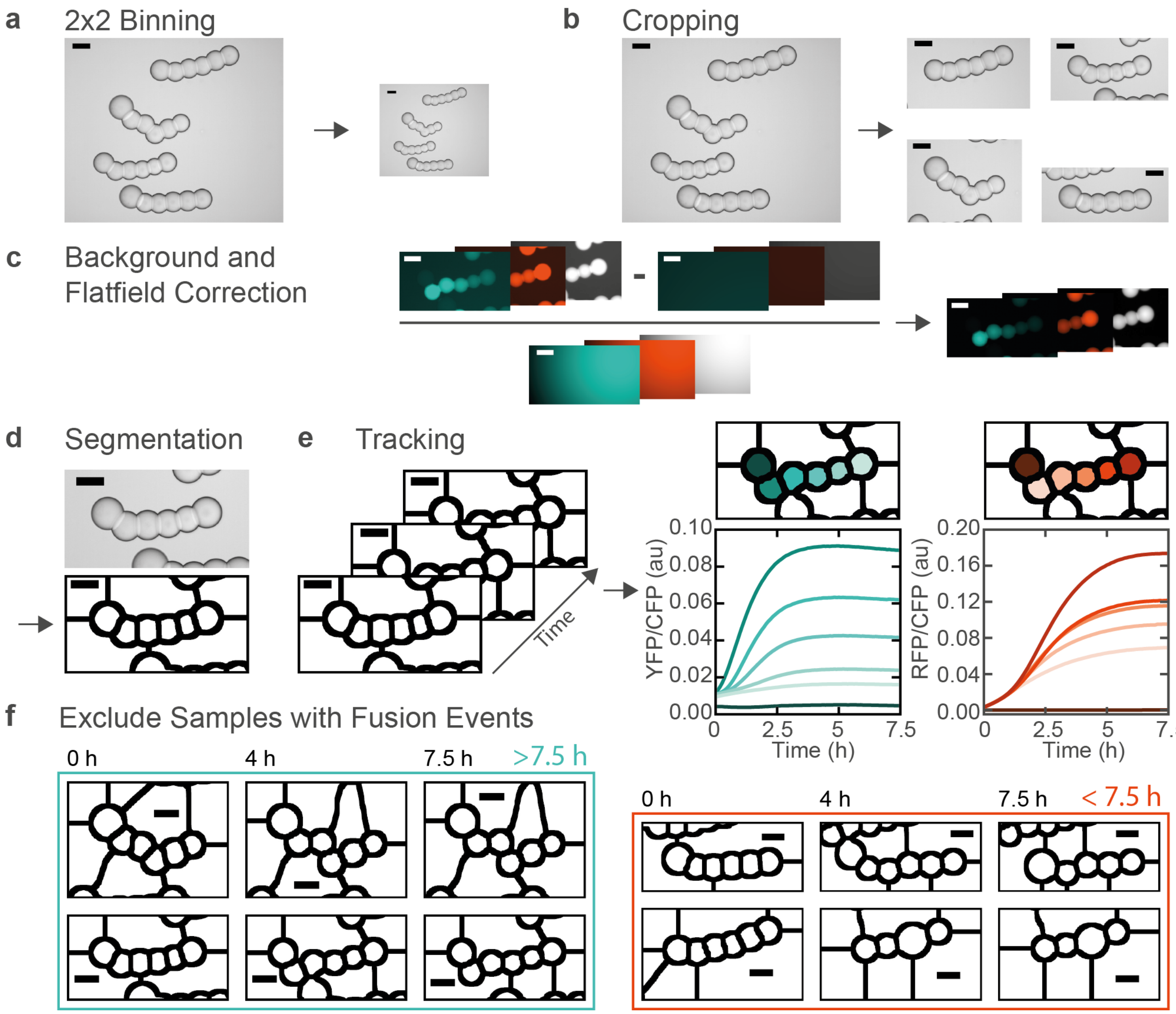

**Supplementary Figure 4 | Summary of the optimized image processing routines. a,** 2x2 binning to reduce memory and computation time. **b,** Sample cropping to reduce computation time and facilitate allocation of samples. **c,** Background and flatfield correction. For visualization the brightness of corrected, uncorrected and background images was scaled between 0 and ~5 times the average background intensity. **d,** Segmentation as described in Supplementary Section 2.7.4. **e,** Droplets were tracked over time based on the segmented masks to extract the fluorescence time traces for the droplets for the YFP, RFP and CFP channel. YFP and RFP time traces were then divided by the measured CFP intensities to yield normalized time traces. **f,** Schematic illustrating the filtering of samples with fusion events earlier than 7.5 hours after starting image acquisition.

The overall optimized image processing workflow is summarized in Supplementary Section 2.8.1, Supplementary Figure 4. The three key aspects that reduced measurement uncertainty – data normalization by reference dye, ‘trimmed’ segmentation and robust background and flatfield correction – are explained in detail in Supplementary Sections 2.8.2, 2.8.3 and 2.8.4. Finally, we describe the estimation of pdfs from our data (Supplementary Section 2.8.5) calculation of PI (Supplementary Section 2.8.6) and PE (Supplementary Section 2.8.7) and discuss the underlying assumptions. All automated image processing routines and schemes for calculation of PE and PI were implemented in a modular fashion using either FIJI or MATLAB. Code examples are available at https://github.com/lauflulu/EMB.

##### 2.8.1 Image Processing

We precede the image processing with a conversion from .nd2 to .tif, combined with a 2x2 binning (Supplementary Figure 4a) of all images to reduce the size and computation time of any downstream processes, without affecting accuracy. In the next step we use an automated cropping routine to create stacks with a single sample (Supplementary Figure 4b). This improves speed and facilitates allocation by assigning unique identifiers to the cropped stacks.

Next, we run the background and flatfield correction (Supplementary Figure 4c, Supplementary Section 2.8.3), as well as the segmentation (Supplementary Figure 4d, Supplementary Section 2.8.4). To obtain fluorescence time traces, we use a MATLAB tracking plugin developed in-house[13,14] (Supplementary Figure 4e), after which we manually selected the segmented areas that represent droplets for all samples that had 6 droplets in the first image. The sender droplet was automatically identified by having the lowest ratio of YFP to reference fluorescence in the first frame and receiver droplets were indexed according to their Euclidean distance from the sender. Finally, we exclude samples in which two droplets fuse at any time within the first 7.5 hours (after which the fluorescence signals saturate) after starting the image acquisition (Supplementary Figure 4f). The occurrence of fusion events corrupts time traces due to a change in the number of samples $N$ and is easily detected by the ending of the tracked time trace of the corresponding droplet. An overview of all samples is provided as Supplementary Data. Excluded samples that did not consist of 6 droplets in the first frame or that underwent a fusion event comprise about 45 % of all samples across all datasets.

##### 2.8.2 Normalization of fluorescence measurements

To optimize our image processing routines, we first need to quantify $\sigma_m$. We therefore acquired a calibration data set consisting of images of several droplets with varying size that were filled with a constant concentration of reporter protein (YFP, or RFP). Hence, the variation in protein concentration due to gene expression variability $\sigma_g \approx 0$ and the measured variability is an estimate of the measurement uncertainty $\sigma_f \approx \sigma_m$. The fluorescence measure $f_i$ for the segmented droplet $i$ that we extract from our video microscopy data must be normalized such that it is proportional to protein concentration. In Supplementary Figure 5 we compare two normalization procedures we called 'normalization by area' and 'normalization by reference'. For each normalization scheme, we compared different image correction (Supplementary Section 2.8.3) and segmentation procedures (Supplementary Section 2.8.4) to determine which combination yields the lowest $\sigma_m$.

For normalization by area we first segment the droplets based on brightfield images (to avoid bias due to differences in fluorescence brightness) and then normalize the sum of fluorescence intensity $F_i$ within a segmented area by its size $A_i$. The functional relation between $F_i$ and $A_i$ depends on the optical setup, as well as on the droplet geometry. We hence fit the calibration function

$$\mathcal{F}(A) = a \cdot A^b \,. \qquad (1)$$

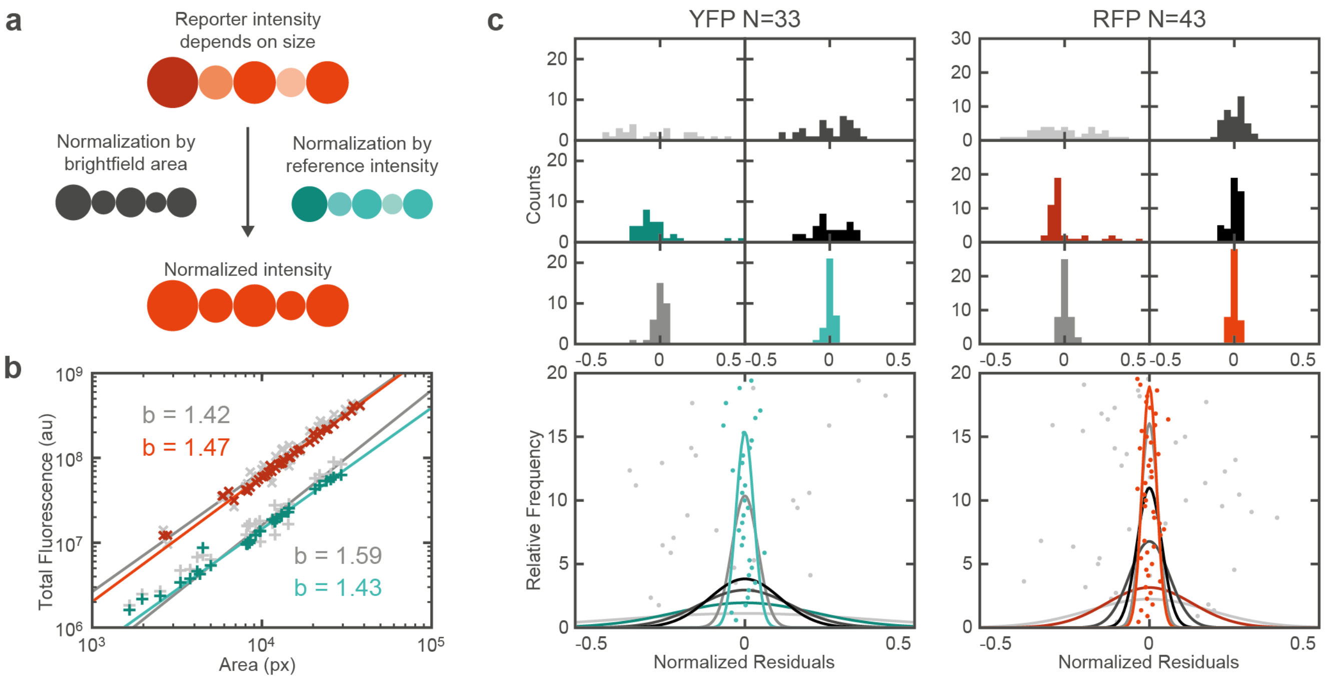

| Normalization | Segmentation | BG | FFC | CV YFP | CV RFP | Color |
|---|---|---|---|---|---|---|
| Area | Precise | - | - | 35.7% | 17.8% | |
| Area | Precise | + | + | 20.4% | 12.7% | |
| Reference | Precise | + | + | 3.8% | 2.5% | |
| Reference | Trimmed | - | - | 13.5% | 5.9% | |
| Reference | Trimmed | + | - | 10.4% | 3.6% | |
| Reference | Trimmed | + | + | 2.6% | 2.1% | |

**Supplementary Figure 5 | Measurement uncertainty of the calibration data set comparing several normalization, segmentation and image correction procedures. a**, Schematic illustrating normalization by area vs. normalization by reference dye. The intensity sum in each segmented area was either divided by a calibrated measure of the droplets size (**c**), or by the intensity sum of a reference dye. **b**, Fits to determine the exponent $b$ required for the normalization by area (Equation (1)). Fits are shown for both YFP and RFP channels, each for the original images and for background (BG) and flatfield corrected (FFC) images. The normalized residuals (Equation (4)) of the fits are shown in (**c**). **c**, Comparison of the measurement uncertainty for different image analysis procedures as indicated by the legend. Individual data points correspond to the normalized residuals. Top: Histograms with bin width 0.02. Middle: Inferred Gaussian pdfs and individual data points (jittered in y direction according to their index). Bottom: Legend listing the respective coefficients of variation (CVs) for the different procedures.

We would typically expect that $1 < b < 3/2$, where $b = 1$ corresponds to a disk-like geometry (high numerical aperture (NA) objective) and $b = 3/2$ corresponds to a sphere-like geometry (low NA). Experimentally, we find values close to $b = 1.5$ (Supplementary Figure 5b), consistent with a low NA of 0.45. We can then calculate our observable by dividing $F_i$ by the calibrated function

$$f_i = F_i/\mathcal{F}(A_i) \propto F_i \cdot A_i^{-b} \,. \quad (2)$$

For normalization by reference, we add an internal reference dye (a CFP, mTurquoise2) that is present in all droplets with a constant concentration. Because the reference fluorescence density $r_i = const.$ and hence $r_i \propto R_i \cdot A_i^{-b} = const.$, the total reference fluorescence $R_i$ is directly proportional to $A_i^b$ (assuming $b$ is similar for reporter and reference fluorescence). Hence we can simply normalize by dividing $F_i$ by $R_i$

$$f_i \propto \frac{F_i \cdot A_i^{-b}}{R_i \cdot A_i^{-b}} = \frac{F_i}{R_i}. \quad (3)$$

Importantly, the geometric factor vanishes, suggesting that his method is insensitive with respect to segmentation uncertainties.

For both normalizations, the relative measurement uncertainty is then given by the coefficient of variation (CV) of $f$

$$\sigma_m \approx \frac{\sigma_f}{\mu_f} = \frac{\sqrt{\langle (f_i - \mu_f)^2 \rangle}}{\mu_f} = \sqrt{\langle (f_i/\mu_f - 1)^2 \rangle}\ , \quad (4)$$

where $\mu_f$ and $\langle \dots \rangle$ denote the mean and we call the quantity $\varepsilon_i = f_i/\mu_f - 1$ the normalized residuals (Supplementary Figure 5c).

We note that our calibration data set includes duplicates from droplets that were imaged in different overlapping microscope positions, but in different relative positions within the field of view. The dominant source of uncertainty is related to the flatfield illumination which depends on the relative position of imaged droplets (cf. also Supplementary Figure 6b, c). We hence purposely kept these duplicates, as keeping them should not bias the $\sigma_m$ estimates, while systematically removing them might.

##### 2.8.3 Illumination related uncertainties

The two major sources of illumination uncertainty are the (varying) background intensity and the typically Gaussian illumination profile characteristic to epifluorescence microscopy[15]. It is common to account for these uncertainties with a flatfield correction (Supplementary Figure 6).

$$\hat{I}_c = \frac{\hat{I} - \hat{B}}{\hat{F}}. \quad (5)$$

The corrected image $\hat{I}_c$ is obtained from the original image $\hat{I}$ by first subtracting the background image $\hat{B}$ and then dividing pixel-wise by the normalized flatfield image $\widehat{F}$.

Besides camera dark noise and background autofluorescence (due to the oil bath), one main component of background light in our setup stems from the sample chamber boundaries. These consist of O-rings grafted onto a glass slide using rather strongly autofluorescent epoxy glue, which causes scattering light. The background is therefore uneven and has to be determined individually for each sample. Therefore, we initially employed the imageJ 'Subtract Background' routine which uses a rolling ball algorithm (typical radius of 200 pixels). However, we found that this method was not sufficiently robust and quantitative for our purpose and therefore implemented an alternative routine based on robust surface fitting of a second degree 2D polynomial in MATLAB (Supplementary Figure 6a). To improve the robustness, the areas that contained droplets were excluded from the fit. Surface fitting is computationally expensive and had to be performed on about $3\ \text{channels} \times 193\ \text{time points} \times 50\ \text{samples} \times 10\ \text{data sets} \approx 300{,}000\ \text{images}$. We therefore first scaled the images down by an empirically determined factor of 0.3 for fitting and used the fitted parameters to obtain the original-sized background image $\hat{B}$.

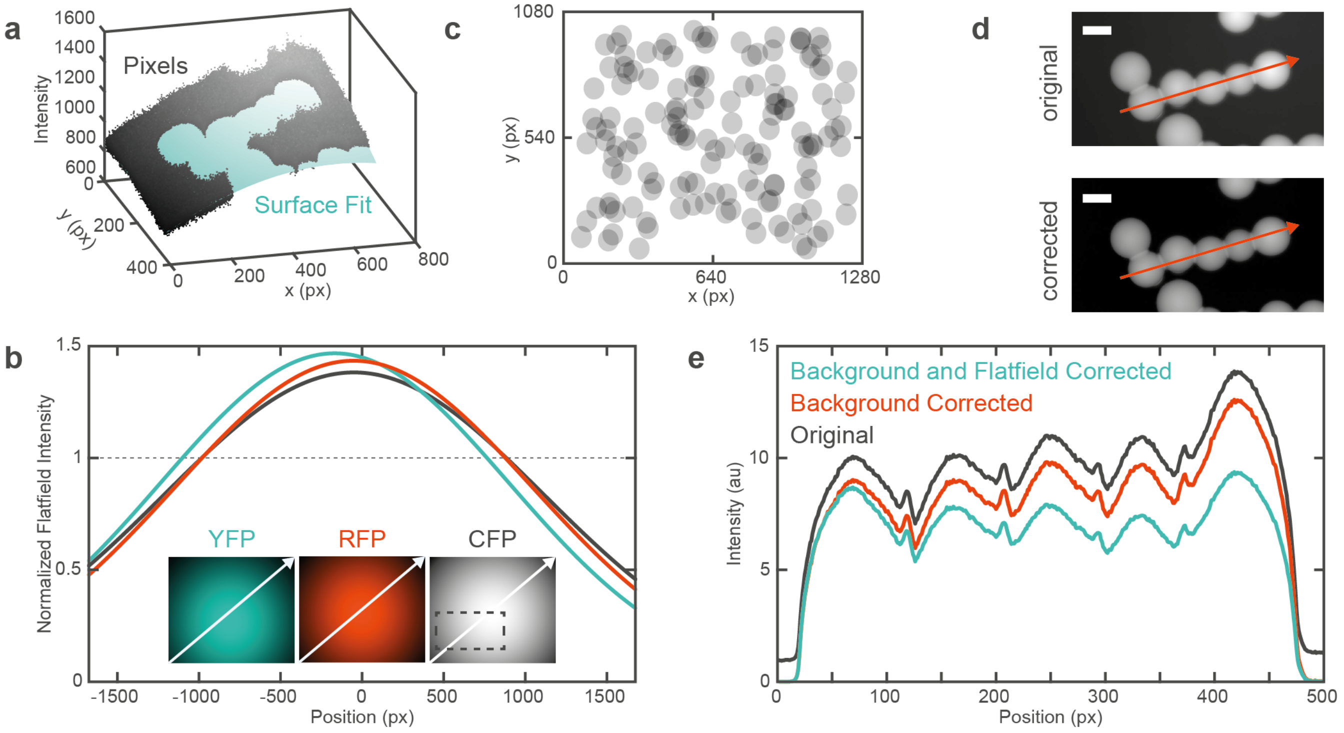

**Supplementary Figure 6 | Illumination uncertainties. a**, 3D representation illustrating the surface fit used to estimate the background for background subtraction. Data points correspond to the intensity of individual pixels and are shown as gray dots. Data points in areas belonging to the sample were excluded from the fit to improve robustness and are not displayed for clarity. Shown is the CFP channel of sample 19_3 from topology A, at 1 mM IPTG, after 4 hours of acquisition. **b**, Flatfield images for the YFP (cyan), RFP (red) and CFP (gray) channels. The arrows indicate the direction of the profile plots. The box in the CFP image indicates the position of the representative images in **a**, **d**. **c**, Distribution of droplets in the calibration data set (Supplementary Figure 5) within the field of view (FOV). The approximately uniform distribution over the FOV highlights the necessity and effectiveness of flatfield correction. **d**, Original and background corrected image from the same sample as in **a**. **e**, Profiles along the long axis of the sample in **d** illustrating the effect of background and flatfield correction.

For flatfield correction (Supplementary Figure 6b) we first recorded 10 flatfield images for each channel using a plain chamber filled with the respective fluorophore and the corresponding darkfield images for the same illumination settings. To mitigate any inhomogeneities, we next took the median intensity of each pixel, subtracted the darkfield image, fitted a 2D Gaussian surface and normalized by dividing by the mean intensity. This yields the normalized flatfield images $\hat{F}$.

Note that the normalization by reference dye already partly amounts to a flatfield correction, but does not account for differences among different channels. Additional flatfield correction therefore decreases the measurement uncertainty significantly (Supplementary Figure 5c).

##### 2.8.4 Segmentation uncertainties

Segmentation of droplets based on brightfield images is commonly achieved by contrast thresholding, or edge detection. We here developed two custom routines, tailored specifically to the two normalization procedures, to thoroughly eradicate potential segmentation related measurement uncertainties. For normalization by area, an 'accurate segmentation' procedure is of critical importance, while we found that for normalization by reference dye a technically simpler 'trimmed segmentation' routine shows a superior performance.

Investigating a typical brightfield image of a droplet assembly (Supplementary Figure 7a) we can easily discern the outer edges as a clear contrast minimum. The DIBs (droplet interface bilayers), however, are less well-defined and may consist of multiple contrast minima or maxima. It is therefore challenging to find a thresholding-based method that accurately locates the outer edges and DIBs simultaneously. We hence decided to first generate images with accurate outer edges and DIBs, separately.

The outline image in Supplementary Figure 7b (top) was generated using the auto-threshold function (imageJ) with the method 'Mean' combined with some filtering and binary operations. Note that the fringes around the assembly can easily be sorted out at a later stage. The bilayer image in Supplementary Figure 7b (bottom) was generated using Sobel filtering (ImageJ function 'find edges'), followed by a Gaussian filter (radius 20) to blur the image, auto-thresholding (Otsu method), binary operations and a watershed transformation (ImageJ plugin 'Adjustable Watershed'[16], radius 5). This procedure robustly yields images where individual droplets are segmented roughly at the bilayer positions, but is inaccurate in finding the exact locations of the outer edges.

We therefore combined these two images using binary and logic operations to get an accurate segmentation regarding both, outer edges and the bilayer positions (Supplementary Figure 7c top). As indicated in the figure, we used these accurate segmentation images to estimate the variability of droplet volumes and bilayer areas in Supplementary Figure 11. However, accurate segmentation combined with normalization by area has three fundamental weaknesses regarding measurement uncertainty. These are visualized in Supplementary Figure 7d, e that show fluorescence images divided by the reference image.

First, because the size of the segmented areas is used to calculate fluorescence, any uncertainty in determining the area (due to varying contrast etc.) propagates to our final fluorescence measure. A related problem is that due to the deformation caused by the bilayers, the exact geometry of each droplet differs, causing a deviation from the calibrated exponent $b$ (Equation (1)). Second, a ~1 px shift between fluorescence channels is common in epifluorescence microscopy[15], and can prevent the accurate location of edges. In Supplementary Figure 7d, e (right panel), this shows as a pronounced peak near the edges of the droplet. Third, as indicated by the gradually increasing fluorescence between droplets (Supplementary Figure 7d, e, left panel), the DIB positions are not well defined in the fluorescence channels either. This may be caused by the bilayer not being parallel to the observation axis[17], or by refraction at the bilayer.

We hence use normalization by reference dye in combination with the 'trimmed segmentation' scheme (Supplementary Figure 7c, bottom) to effectively avoid all these potential sources of uncertainty. A 'trimmed segmentation' image is simply generated by eroding the bilayer image (Supplementary Figure 7b, bottom) 10 times to ensure that the segmented areas do not include any pixels close to the edge or bilayer. Due to the normalization by reference dye, the normalized fluorescence within the segmented areas is approximately constant (Supplementary Figure 7d, e). This property renders this method insensitive to the exact size and location of the segmented area and additionally partly corrects for illumination uncertainties.

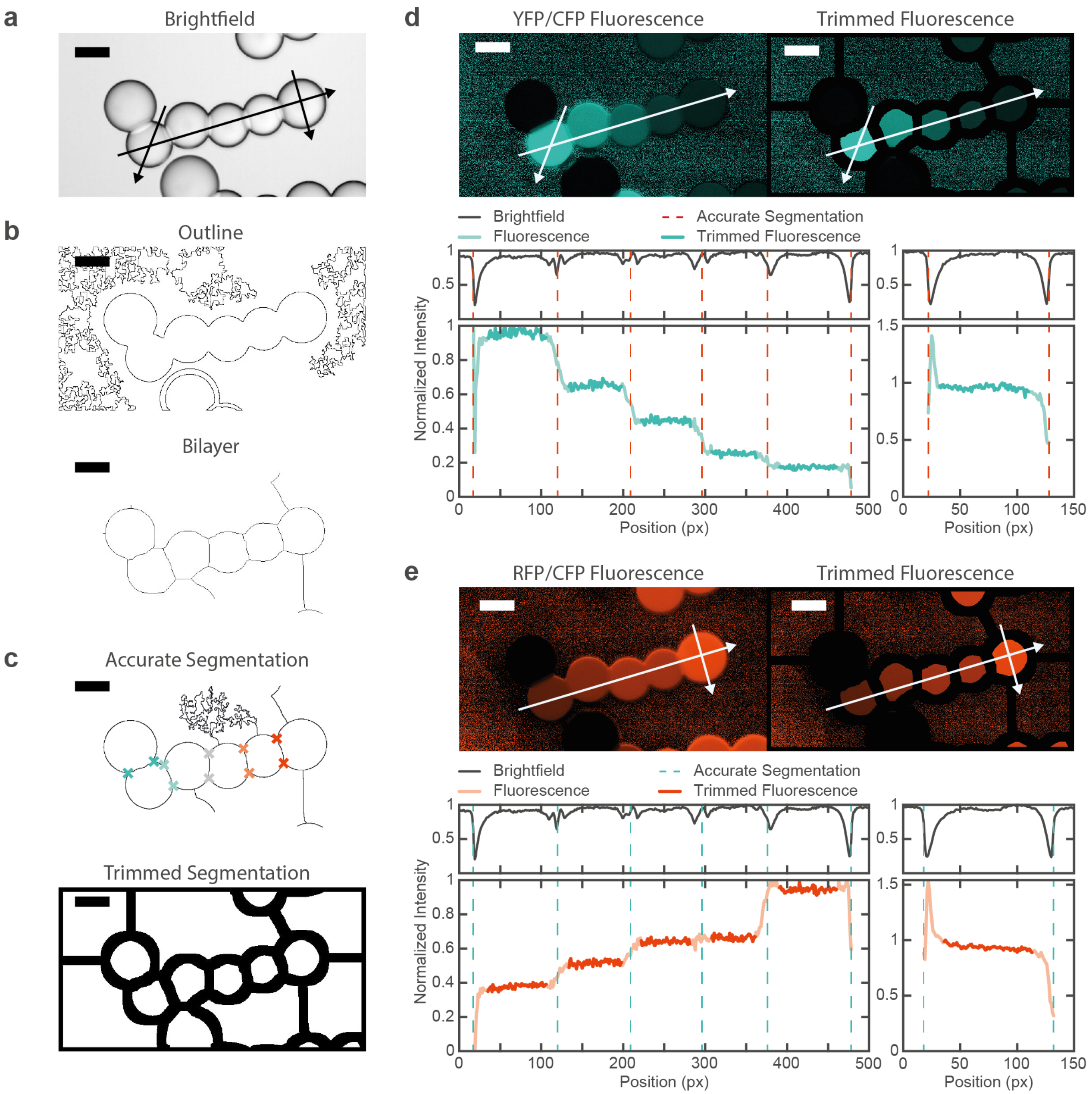

**Supplementary Figure 7 | Segmentation uncertainties. a,** Representative brightfield image. While the outer edges of the droplet assembly appear as defined contrast minima, the position of the droplet interface bilayers is less clearly defined. Arrows indicate positions of brightfield profiles in **d** and **e**. **b,** Binary images generated by thresholding the brightfield image (**a**). The outline image was generated by thresholding (Mean), and using the particle analyzer tool and additional binary operations. The bilayer image was generated by using 'Find edges', a Gaussian blur, thresholding (Otsu) and adjustable watershed. **c,** Segmentation images used for image analysis. The accurate segmentation image was generated by merging the outline and bilayer image (**b**) using binary operations and logic functions. Accurate segmentation was used to estimate the variability of droplet volumes and bilayer areas (Supplementary Figure 11) by measuring the distance between vertices, as indicated, as well as to grade the performance of normalization by area procedures in Supplementary Figure 5. The trimmed segmentation image was used to measure fluorescence with normalization by reference dye and was simply generated by eroding the bilayer image (**b**). **d-e,** Illustration of segmentation uncertainties for the d, YFP and e, RFP channel. The top left image shows the background and flatfield corrected YFP image divided by the CFP (reference) image. Note, we uniformly added +10 to the reference image to mitigate division by 0 issues for visualization, only. The top right image shows the same image but multiplied by the trimmed segmentation image **c**. Arrows indicate positions of profile plots below. As indicated by the relatively constant fluorescence within one droplet, using trimmed segmentation avoids measurement uncertainties due to the undefined bilayer positions, pixel shifts as well as size estimates. All images shown are from dataset 3 (topology A, 1 mM IPTG) sample 19_3, frame 49 (4 hours) (cf. Supplementary Data) of topology A, 1 mM IPTG dataset.

##### 2.8.5 Estimation of Probability Distribution Functions

For each of our 10 data sets, the image processing yields $N = 9 - 37$ samples drawn from an underlying joint probability distribution function (pdf) $p(\{g_i\}, x, t)$ (Supplementary Figure 8), which needs to be estimated before calculating PI (Supplementary Figure 9) and PE (Supplementary Figure 10). One set of fluorescence time trace data can be represented as a 4-dimensional array consisting of $N$ samples, $I = 2$ genes, $X = 5$ positions and $T = 91$ time points (Supplementary Figure 8a). For consistency, the data is normalized to the maximum across all $x$ and $t$ and mean over $n = 1, \dots, N$ intensity for each gene $i$. Since positions and time points are uniformly distributed with marginal distributions $p_x(x) = 1/5$ and $p_t(t) = 1/T$, respectively, the joint conditional pdf is $p(\{g_i\}|x, t) = 5 \cdot T \cdot p(\{g_i\}, x, t)$.

As illustrated in Supplementary Figure 8b, $p(\{g_i\}|x, t)$ can be estimated by binning the data or by assuming an underlying distribution. For a multivariate Gaussian distribution

$$p(\{g_i\}|x,t) = (2\pi)^{-\frac{I}{2}} |C(x,t)|^{-\frac{1}{2}} \exp\left[-\frac{1}{2}\sum_{i,j=1}^{I} \left(g_i - \bar{g}_i(x,t)\right)[C^{-1}(x,t)]_{ij} \left(g_j - \bar{g}_j(x,t)\right)\right], \quad (6)$$

the mean morphogen concentration $\bar{g}_i(x)$ and covariance matrix $C_{ij}(x)$ can be obtained from the fluorescence data as

$$\bar{g}_i(x,t) = \frac{\sum_{n=1}^{N} g_i(x,t)}{N}, \quad (7)$$

$$C_{ij}(x,t) = \frac{\sum_{n=1}^{N} g_i(x,t) g_j(x,t)}{N-1} - \bar{g}_i(x,t)\bar{g}_j(x,t). \quad (8)$$

The same procedure here defined in general for the multiple gene joint conditional pdf $p(\{g_i\}|x, t)$ also applies for the estimation of single gene conditional pdfs $p(g_i|x, t)$ (Supplementary Figure 8c, d). While estimating PI from binned pdfs requires relatively large sample sizes to correct for binning bias (Supplementary Section 2.8.6), the approximation of the underlying distribution by a Gaussian has to be properly justified. Besides the verification schemes discussed in Supplementary Section 2.8.6, additional support for assuming a Gaussian distribution can be gained by fitting a representative set of cumulative data (Supplementary Figure 8e, f). Good fits are obtained with a cumulative Gaussian pdf $c(g|x) = \frac{1}{2}\left(\mathrm{erf}\left(\frac{g-b}{a}\right) + 1\right)$, with deviations increasing in the droplets distant from the sender $x = 3{,}4{,}5$.

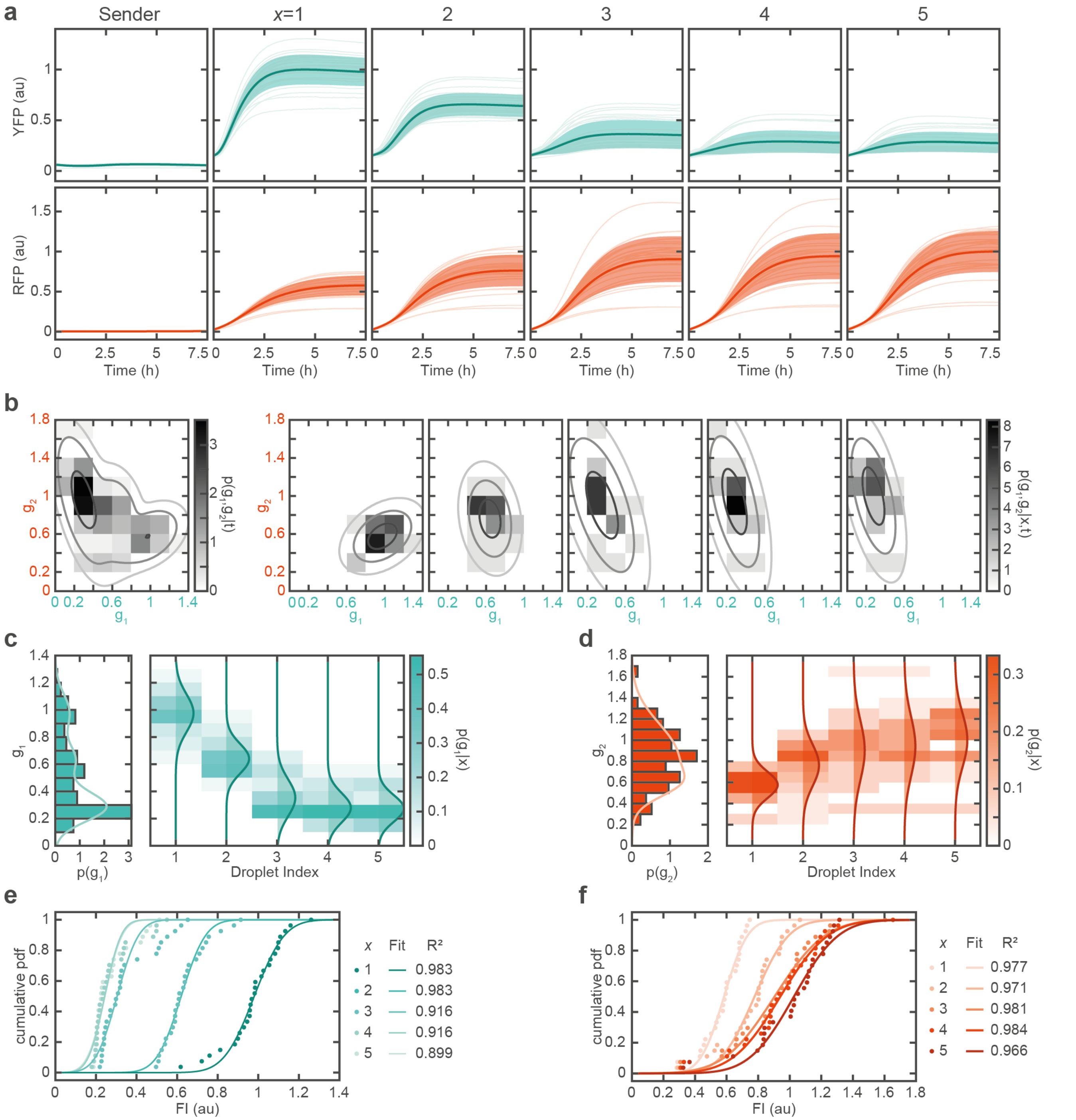

**Supplementary Figure 8 | Representing fluorescence data by probability distribution functions. a,** Individual fluorescence time traces (faint) with mean and standard deviation for the respective droplet. **b**, Joint marginal $p(g_1, g_2|t = 7.5\ h)$ and conditional pdf $p(g_1, g_2|x, t = 7.5\ h)$ represented as binned histograms ($B = 5$) as indicated by color. Overlaid are contour lines of Gaussian distributions inferred from the sample mean and SD at 1,2, and 3 SDs. **c-d,** Single gene marginal $p(g)$ and conditional pdf $p(g|x)$ for gene $g_1$ and $g_2$, represented as a histogram ($B = 10$) as indicated by color. Overlaid are Gaussian distributions inferred from the mean and standard deviation at the respective position. **e-f,** Cumulative pdf for $p(g|x)$. The cumulative Gaussian fits were performed to justify the Gaussian approximation used to calculate PE and PI.

##### 2.8.6 Calculation of Positional Information

In the following, we briefly introduce the formal definitions for PI and PE and discuss the methods presented in Tkačik, et al. (2015)[10] regarding the calculation of PI and PE estimates from real data.

Positional information is defined as the mutual information that the (joint) morphogen concentrations $\{g_i\}$ contain about position $x$ and vice versa

$$\mathcal{I}(x;\{g_i\}) = \int dx\, p_x(x) \int d^n\boldsymbol{g}\, p(\{g_i\}|x)\, \log_2 \frac{p(\{g_i\}|x)}{p_g(\{g_i\})} = \mathcal{I}(\{g_i\};x)\,. \quad (9)$$

Here $p_x(x) = 1/5$ and $p_g(\{g_i\}) = \int dx\, p_x(x)\, p(\{g_i\}|x)$ are the marginal distributions. Note that for conciseness we here have omitted to explicitly write down the time dependence of PI and the pdfs, but all calculations can simply be performed for each given time point individually to obtain the temporal evolution of PI. Equation (9) can be rewritten as the difference of the Shannon entropy $S[p(y)] = -\int dy\, p(y) \log_2 p(y)$ of the marginal and conditional pdf

$$\mathcal{I}(\{g_i\};x) = S\left[p_g(\{g_i\})\right] - \langle S[p(\{g_i\}|x)]\rangle_x \quad (10)$$

The first term in Equation (10) is called 'total entropy', the second term 'noise entropy'. Total entropy measures the range of gene expression levels that are available to the droplets, while noise entropy quantifies the 'loss' of information due to the variability in the gene expression levels at a given position. Using Equation (10) to estimate PI has a practical advantage over Equation (9) as discussed in the following paragraph.

Tkačik, et al. (2015)[10] present several alternative methods to estimate PI from real data, called the direct (DIR) method, the first and second Gaussian approximation (FGA and SGA) and a Monte Carlo integration scheme (MCI). In the following discussion we first describe the DIR and SGA estimation methods, which are well suited to the specifics of our data sets. We then briefly discuss FGA and MCI, which in our case provide only minor advantages and were therefore not considered in detail.

The DIR method is attractive as it does not build on any prior assumptions about the distribution $p(\{g_i\}|x)$. However, the sample sizes required to estimate PI with a binned pdf are relatively large and grow exponentially with the number of genes $I$. A binned pdf can be obtained by dividing the fluorescence data at each location $x$ into $B$ bins of size $\Delta_g = 1/B$ (Supplementary Figure 8b, c). Intuitively, if $B$ is chosen too low, data is grouped into large bins and information is inevitably lost. Conversely, we can find a critical value $B^*$ above which the pdfs become sparse. Naive PI estimates obtained by simply inserting a binned pdf into Equations (9) or (10) therefore suffer from an estimation bias.

As for $B < B^*$, the estimation bias scales as $1/N$ and $1/B^{I+1}$ [18,19], a DIR estimate can be obtained by extrapolating naive estimates for a series of fractions of the whole data with size $M < N$ and varying bin number $B < \mathrm{B}^*$ towards infinite sample size $M \to \infty$ and zero bin size $\Delta_g \to 0$ via linear regression (Supplementary Figure 9a, b). In practice, we first generate $K = 100$ sub-samples for each sample size $M = (6,7,\dots,11)\cdot\frac{N}{12} < N$ (rounded up to the next digit) randomly drawn from the full data set without replacement. Note, that for data sets with $N \leq 12$ we instead used $M = (7,8,\dots,N)$.

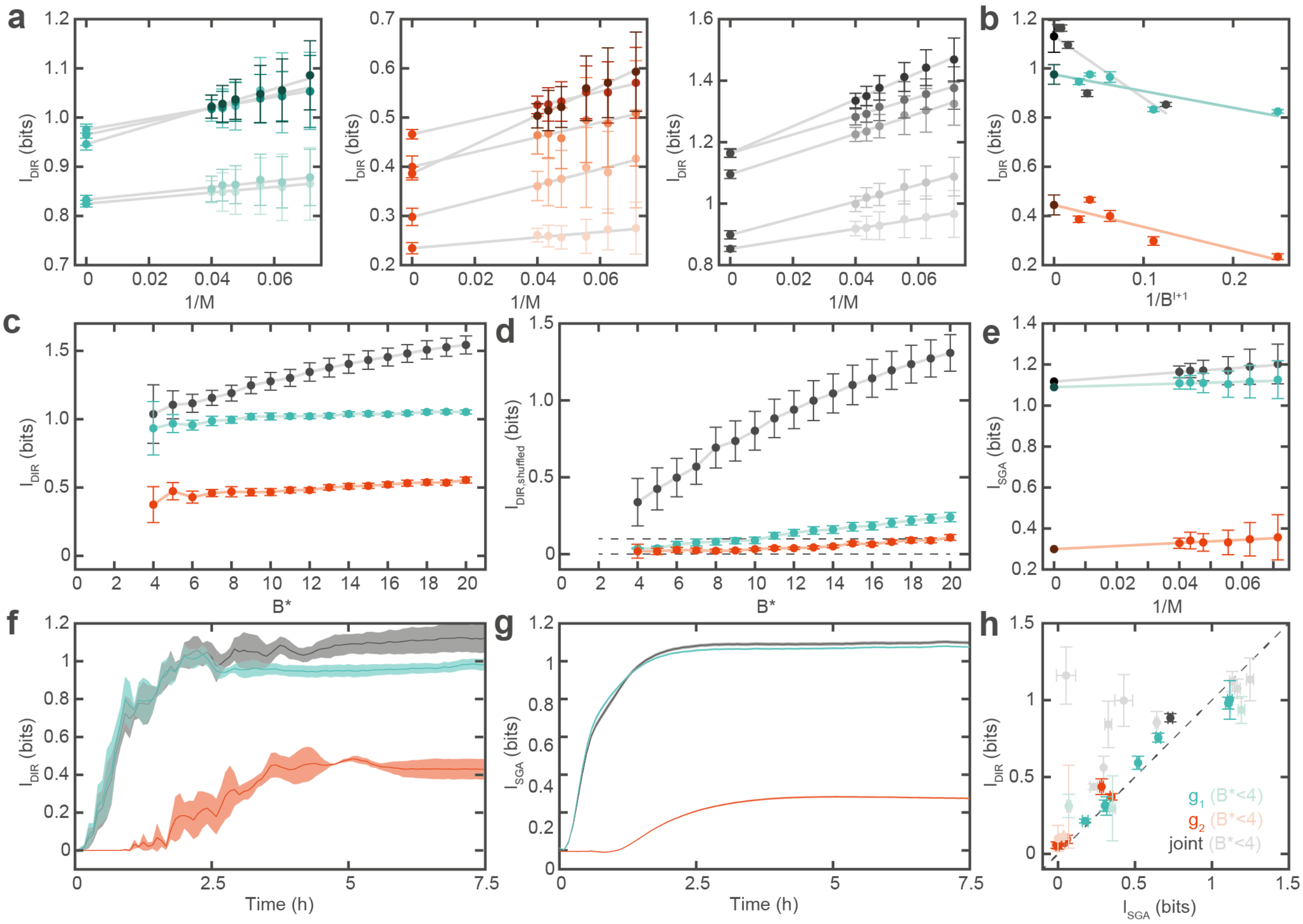

**Supplementary Figure 9 | Positional Information. a, b,** Extrapolation of positional information using the direct method (DIR), to **a**, infinite sample size and **b**, infinite bin number. **c,** Extrapolated DIR estimates for varying maximum bin numbers $B^*$. For single gene PI the estimates are approximately constant over the range tested, while for the joint PI the estimates systematically increase with increasing $B^*$. **d,** Extrapolated shuffled DIR estimates for varying $B^*$. Values above zero indicate a bias due to under-sampling. Horizontal dashed line indicates the tolerance of 0.1 bits. **e,** Extrapolation of naive estimates using the second gaussian approximation (SGA) to correct for sample size bias. **f-g,** Extrapolated DIR and SGA estimates, respectively, over time. **h,** Comparison of DIR and SGA estimates for all data sets. Estimates with insufficient data for application of the DIR method ($B^*$<4) are grayed out. **a-h,** Error bars and shaded areas represent 68% confidence intervals of the extrapolated values (t-corrected standard errors). Error bars on non-extrapolated values in a represent the SD of 100 randomly sampled datasets. In **g**, error bars are in the order of the line width. All data shown here is for data set 3 and is representative for all data sets (cf. Supplementary Data).

Then, we compute naive PI estimates for each of the sub-samples for a series of bin sizes $B = (2,3,4,5,6)$ (as shown below $B^* = 6$ was a reasonable compromise for all our data sets). Note that, per convention, we define $B$ relative to the maximum mean gene expression profile, which we have normalized to 1. The actual bin number is chosen such that it spans the entire range of gene expression profiles and is thus usually larger than $B$. Also note that because some elements of a binned pdf will be 0, we need to (informally) set $0 \cdot \log(0) = 0 \cdot \log(0/0) = 0$ which follows from continuity (Cover and Thomas (2005)[20], p.31).

For each $(B, M)$ we then compute the mean and standard deviation (SD) of the $K$ subsamples and linearly regress $\mathcal{I}_{B,M} = \mathcal{I}_B + c_M \frac{1}{M}$ (Supplementary Figure 9a). As $\frac{1}{M} \to 0$, $M \to \infty$ and hence the intersects $\mathcal{I}_B$ are the corrected estimates in the infinite data limit. In a second extrapolation (Supplementary Figure 9b) we can correct for the finite bin size by linearly regressing these $I_B$ for the different bin numbers via $\mathcal{I}_B = \mathcal{I}_0 + c_B \frac{1}{B^{I+1}}$. Again as $\frac{1}{B^{I+1}} \to 0$, we approach the limit of infinitely small bin size and $\mathcal{I}_0$ is the final PI estimate. To gauge the precision of this estimation procedure, we take the 68% confidence interval of the last extrapolation as statistical uncertainty.

To find $B^*$ for a given data set, we can perform a series of PI estimates with varying $B^*$ [19]. As shown in Supplementary Figure 9c an increase in the PI estimate, as noticeable for the joint PI estimate ($I = 2$), indicates an overestimation due to undersampling. Additionally, we can find $B^*$ by randomly shuffling positions and gene expression levels[19] (Supplementary Figure 9d), which should destroy any mutual information ($\mathcal{I}_{shuffled} = 0$), unless $B^*$ is chosen too large. We hence define $B^* = \max(B|\mathcal{I}_{shuffled} + \sigma_{\mathcal{I}_{shuffled}} < \tau)$, where we choose a tolerance $\tau = 0.1$. The minimum requirements to ensure at least 3-4 data points for each extrapolation fit are roughly $B^* \geq 4$ and $N \gtrsim 10$. However, since $B^*$ decreases with decreasing $N$, increasing $\sigma_g$, and increasing $I$, $N$ may need to be considerably higher to ensure $B^* \geq 4$. Our data sets have $N = 9 - 37$ and for single genes, the conditions for the extrapolation are mostly met (Supplementary Table 3). However, for calculating the joint information ($I = 2$), we hardly reach $B^* \geq 4$. As DIR is hence not applicable in these cases, we additionally use SGA.

To obtain an SGA estimate, we first approximate $p(\{g_i\}|x)$ by a Gaussian distribution (Equation (6)), where the mean and covariance can be calculated from the data (Equations (7) and (8)). This eliminates the binning problem, which aids PI estimation for multiple genes. We can then calculate the total and the noise entropy by inserting an inferred pdf into Equation (10). This is done numerically using a discretized multivariate normal pdf with a grid spacing of 0.01 (in the order of the smallest $\sigma_{g_i}(x,t)$). Then, we again extrapolate a series of naive estimates to infinite sample size (Supplementary Figure 9e).

**Supplementary Table 3 | Data set overview.** S: data set number (cf. Supplementary Data), T: topology, IPTG: inducer concentration in sender droplet, $N$: sample size. $B^*$ are the critical bin numbers determined by shuffling gene expression levels and positions for each data set. '<': $B^* < 4$, i.e. DIR not applicable. '>': $B^* > 20$, i.e. higher than the range tested. All DIR estimates are for $B^* = 6$. DIR/ SGA estimates are after 7.5 hours.

| S | T | IPTG (mM) | $N$ | $\mathcal{I}_{\mathrm{DIR},1}$ ($B^*$) (bits) | $\mathcal{I}_{\mathrm{SGA},1}$ (bits) | $\mathcal{I}_{\mathrm{DIR},2}$ ($B^*$) (bits) | $\mathcal{I}_{\mathrm{SGA},2}$ (bits) | $\mathcal{I}_{\mathrm{DIR},1,2}$ ($B^*$) (bits) | $\mathcal{I}_{\mathrm{SGA},1,2}$ (bits) |
|---|---|---|---|---|---|---|---|---|---|
| 1 | A | 100 | 9 | 0.10±0.19 (<) | 0.06±0.02 | 0.29±0.29 (<) | 0.06±0.01 | 0.77±0.39 (<) | 0.00±0.11 |
| 2 | A | 10 | 35 | 0.64±0.03 (>) | 0.56±0.01 | 0.38±0.02 (13) | 0.36±0.01 | 0.88±0.07 (<) | 0.67±0.01 |
| 3 | A | 1 | 27 | 1.01±0.03 (7) | 1.08±0.02 | 0.44±0.04 (17) | 0.29±0.02 | 1.14±0.07 (<) | 1.11±0.02 |
| 4 | A | 0.1 | 18 | 0.31±0.07 (7) | 0.36±0.03 | 0.07±0.03 (7) | 0.02±0.01 | 0.87±0.14 (<) | 0.37±0.03 |
| 5 | B | 100 | 25 | 0.17±0.02 (>) | 0.17±0.01 | 0.06±0.01 (12) | 0.00±0.01 | 0.39±0.01 (<) | 0.24±0.02 |
| 6 | B | 10 | 37 | 0.70±0.02 (>) | 0.67±0.02 | 0.08±0.03 (>) | 0.07±0.01 | 0.78±0.02 (4) | 0.76±0.01 |
| 7 | B | 1 | 11 | 0.40±0.13 (<) | 0.35±0.02 | 0.19±0.12 (<) | 0.00±0.00 | 1.07±0.18 (<) | 0.35±0.05 |
| 8 | B | 0.1 | 17 | 0.95±0.08 (<) | 1.15±0.02 | 0.09±0.01 (<) | 0.03±0.01 | 1.16±0.12 (<) | 1.19±0.03 |
| 9 | C | 100 | 18 | 0.29±0.05 (9) | 0.27±0.01 | 0.07±0.01 (17) | 0.00±0.01 | 0.56±0.09 (<) | 0.26±0.01 |
| 10 | C | 10 | 22 | 0.94±0.17 (10) | 1.13±0.01 | 0.05±0.02 (>) | 0.00±0.01 | 1.03±0.15 (<) | 1.18±0.01 |

Importantly, the Gaussian distribution is the maximum entropy distribution meaning that the SGA estimates can be understood as a lower bound[10]. To see how close the SGA estimates approximate the real PI value, we can compare them to DIR estimates for the data sets for which DIR is justified (Supplementary Figure 9f-h). DIR and SGA estimates agree well both over time and across all data sets.

FGA is a hybrid approach that exploits the fact that $p_g(\{g_i\})$ is sampled better than $p(\{g_i\}|x)$ because it includes the $g$ measurements from every $x$. Hence, the total entropy in Equation (10) can be estimated directly, while the noise entropy can be calculated using the Gaussian approximation. However, our system only has 5 naturally discrete positions, whereas in Tkačik, et al. (2015)[10] for the case of Drosophila embryos the continuous expression profiles were binned into 1000 spatial bins. Hence, in our case the sampling of $p_g(\{g_i\})$ is not significantly better than $p(\{g_i\}|x)$ and consequently, FGA provided only minor advantages over DIR.

MC integration becomes relevant when estimating high dimensional joint PI carried by $I \geq 3$ genes. Then the fraction of the 'interesting' phase space becomes small and computations inefficient due to the curse of dimensionality. As we are interested in $I \leq 2$, we did not consider MC integration.

##### 2.8.7 Calculation of Positional Error

Positional error $\sigma_x(x)$, here defined in units of droplets, can be calculated from the Fisher information

$$\mathcal{J}(x) == \int d^I g \; p(\{g_i\}|x) \cdot (\partial_x \log p(\{g_i\}|x)\,)^2 \,. \tag{11}$$

In combination with the Cramér-Rao bound we get a lower limit for the positional error

$$\sigma_x^2(x) \geq \frac{1}{\mathcal{J}(x)} \,. \tag{12}$$

We can again approximate $p(\{g_i\}|x)$ with a Gaussian distribution Equation (6) which inserted into Equation (11) gives[10]

$$\mathcal{J}(x) = \; (\partial_x \, \bar{\boldsymbol{g}}(x))^T \boldsymbol{C}(x)^{-1} (\partial_x \, \bar{\boldsymbol{g}}(x)) \; + \frac{1}{2} \mathrm{Tr}[\boldsymbol{C}(x)^{-1} (\partial_x \boldsymbol{C}(x)) \boldsymbol{C}(x)^{-1} (\partial_x \boldsymbol{C}(x))] \,, \tag{13}$$

from which a lower bound for PE can be calculated from the mean and covariance (Equations (7) and (8)). Equation (13) is more intuitive to understand in the case of a single gene with the mean morphogen profile $\bar{g}(x) = \bar{g}_i(x)$ and variance $\sigma_g^2(x) = C_{ii}(x)$. Then Equation (13) reduces to

$$\mathcal{J}(x) = \frac{\left(\partial_x \, \bar{g}(x)\right)^2}{\sigma_g^2(x)} + 2 \, \frac{\left(\partial_x \, \sigma_g(x)\right)^2}{\sigma_g^2(x)} \;, \tag{14}$$

where the first term represents the information that can be retrieved from the gradient of the mean profile and the second term represents information that can be retrieved from spatial variations in the noise (Supplementary Figure 10a).

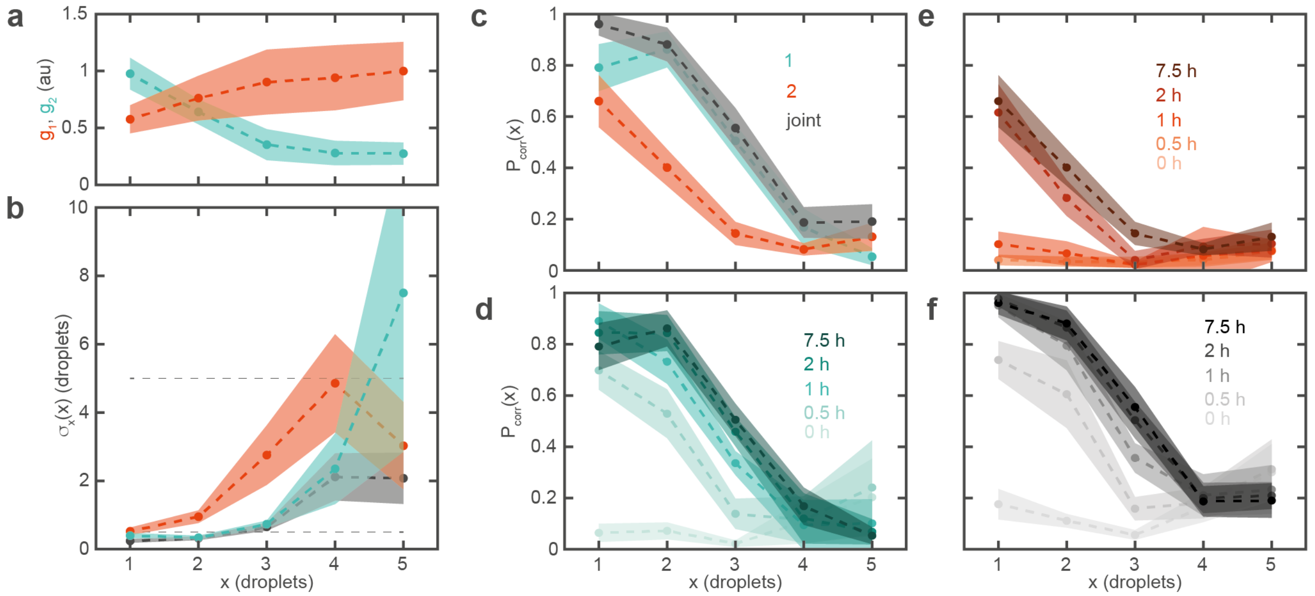

**Supplementary Figure 10 | Positional Error. a,** Mean fluorescence profiles with standard deviation. **b,** PE at the respective droplet positions. Dashed lines indicate a positional error of 1 and 5 droplets (i.e. the length of the entire sample). Lines are a guide to the eye. **c,** Probability of correctly identifying droplet position after 7.5 hours. **d-f,** Temporal evolution of $P_{corr}$. Error bars in b-f are standard deviations estimated by bootstrapping and Gaussian error propagation.

In practice, we first calculate PE by extrapolating a series of naive PE estimates (Equation (13) with Equation (12)) to the infinite sample size limit ($K = 100$, $M = (6,7,\ldots,11) \cdot \frac{N}{12} < N$ as for PI). The mean and SD of PE are then estimated by naively bootstrapping 500 times.

$\sigma_x(x) \in \mathbb{R}^+$ diverges for small $\mathcal{J}(x)$, which is inconvenient for visualization (Supplementary Figure 10b). We therefore consider the probability that the position of a given droplet can be correctly identified $P_{corr}(\sigma_x(x)): \mathbb{R}^+ \to [0,1]$ (Supplementary Figure 10c-f)

$$P_{corr}(x) = \frac{1}{\sqrt{2\pi\sigma_x^2(x)}} \int_{\tilde{x}-\frac{\Delta x}{2}}^{\tilde{x}+\frac{\Delta x}{2}} \exp\left(-\frac{(x'-\tilde{x})^2}{\sigma_x(x)}\right) dx' \,, \tag{15}$$

where $\Delta x = 1$ is the spacing between two droplets and $P_{corr}(x) \to 0$ as $\sigma_x(x) \to \infty$ and $P_{corr}(x) \to 1$ as $\sigma_x(x) \to 0$. After a coordinate transformation $x'' - 0.5 = x' - \tilde{x}$, the analytical solution to the integral is

$$P_{corr}(x) = \frac{1}{\sqrt{2\pi\sigma_x^2(x)}} \int_0^1 \exp\left(-\frac{(x''-0.5)^2}{\sigma_x(x)}\right) dx'' \approx \text{erf}\left(\frac{0.353553}{\sigma_x(x)}\right). \tag{16}$$

The uncertainty $\Delta P_{corr}(x)$ can be estimated via Gaussian error propagation

$$\Delta P_{corr}(x) = \left|\frac{\partial P_{corr}(x)}{\partial \sigma_x(x)} \Delta\sigma_x(x)\right| \approx \frac{0.398942}{\sigma_x^2(x)} \exp\left(-\frac{1}{8\,\sigma_x^2(x)}\right) \Delta\sigma_x(x)\,. \tag{17}$$

We note that one important consideration specific to our system is the question of how to calculate the discrete derivative $\partial_x f(x)$ of function $f(x)$ (mean or covariance matrix). As our system represents a discrete space of length $L = 5$, we cannot choose any arbitrary bin width smaller than $\Delta x = 1$ to improve accuracy. Hence, to get a suitable estimate of $\partial_x f(x)$ for all 5 positions, we take the discrete forward derivative at the first droplet, the centered derivative at the middle droplets ($x = 2{,}3{,}4$), and the backward derivative at the last droplet, i.e.

$$\partial_x f(1) = \frac{f(2) - f(1)}{1}, \tag{18}$$

$$\partial_x f(x = 2{,}3{,}4) = \frac{f(x+1) - f(x-1)}{2}, \tag{19}$$

$$\partial_x f(5) = \frac{f(5) - f(4)}{1}. \tag{20}$$

### 3 Models

#### 3.1 Diffusion in droplet assemblies

##### 3.1.1 General diffusion equation

Diffusion in droplet assemblies consists of diffusion in three dimensions in the aqueous droplets and of permeation of the lipid bilayer in quasi-one dimension. In linear assemblies of droplets, we only considered diffusion in 1D along the axis of the assembly. This was modeled as described in Dupin & Simmel (2019)[6]. Briefly, the thickness of the interfaces ($L$ = 5 nm) is much smaller than the diameter of the droplets ($l$ ≈ 100 μm), which leads to an effective diffusion coefficient $D_{eff}$[21]:

$$\frac{1}{D_{eff}} = \frac{1}{D} + \frac{1}{P \cdot l} \tag{21}$$

$D$ is the free diffusion coefficient of the considered signal, and is typically on the order of 1000 μm$^2$/s for small chemicals, based on the Stokes-Einstein equation. $P$ is the permeability of the bilayer to the signal (units: m/s). The change in concentration $n_i$ of the signal in droplet $i$ can be described as:

$$\frac{dn_i}{dt} = \sum_{j \in \{neighbors\ of\ i\}} -\frac{1}{V_i} \frac{D \cdot P \cdot A_{ij}}{D + P \cdot l_i} (n_i - n_j) \tag{22}$$

$V_i$ is the volume of droplet $i$ and $A_{ij}$ is the effective area of transport of the interface between droplets $i$ and $j$. For unspecific, non-pore-mediated diffusion of the signal across the bilayer (as is the case with the signals used here), $A_{ij}$ is the area of the whole bilayer, which is approximately $8 \cdot 10^3$ □m$^2$ on average. This model is used to fit the permeability of the bilayer to IPTG signal.

##### 3.1.2 Geometrical parameters

The geometry of the droplet assemblies is a key parameter of the signal diffusion process. Particularly, the effective area of transport of the interface $A_{ij}$ is a crucial component in the kinetics of the permeation step, and the length of the compartment $l_i$ is crucial to the bulk diffusion step. They are calculated as follows[17].

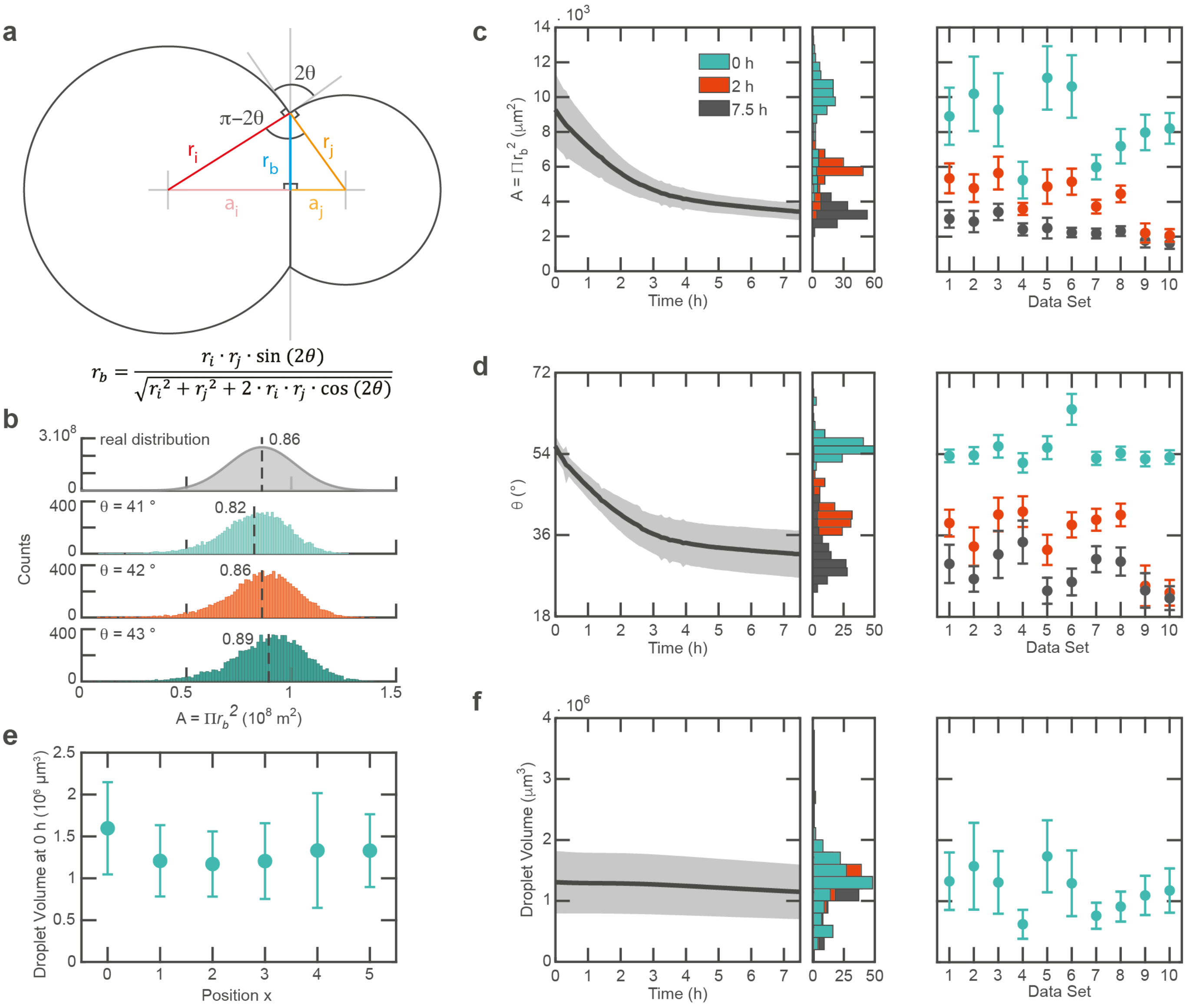

**Supplementary Figure 11 | Geometrical parameters of the droplet assemblies. a,** Scheme of a droplet bilayer of radius $r_b$ (blue), involving two droplets of respective radii $r_i$ (dark red) and $r_j$ (dark orange), truncated droplet radii $a_i$ (light red) and $a_j$ (light orange), with contact angle $\theta$ (black). **b,** Comparison of the real distribution (top panel, assumed Gaussian, with mean $0.86 \cdot 10^8$ μm$^2$ and standard deviation $0.16 \cdot 10^8$ □m$^2$) of droplet bilayer areas and simulations with different contact angles (from the top, second panel $\theta$ = 41°, third panel $\theta$ = 42°, fourth panel $\theta$ = 43°). The simulated bilayer area were calculated from 1000 2-droplet assemblies with each volume randomly picked from a Gaussian distribution ($V$ = 1.2 $\pm$ 0.42 nL, as measured in experiments) for each contact angle. $\theta$ = 42° showed the best fit to the data and was taken as the contact angle value in all other simulations. **c-f**, Bilayer area $A$ (**c**), contact angle $\theta$ (**d**) and droplet volume $V$ (**e,f**) of a set of experiments over time. Mean is indicated by a thick line and standard deviation by a shaded area. Histogram: distribution at 0 h, 2 h and 7.5 h.

The length of the compartments corresponds to the diameter of a sphere truncated by a spherical cap of base $A_{ij}$ on both ends for the central droplets and the diameter of a sphere truncated by a spherical cap of base $A_{ij}$ on one end for the edge droplets. The length of the middle compartments is defined by the droplet volume and the bilayer area:

$$l_i = \sqrt{\left(\frac{3 \cdot V_i}{4\pi}\right)^{\frac{2}{3}} - \frac{A_{i-1,i}}{\pi}} + \sqrt{\left(\frac{3 \cdot V_i}{4\pi}\right)^{\frac{2}{3}} - \frac{A_{i,i+1}}{\pi}} \tag{23}$$

The length of edge droplets is, for example for droplet 1:

$$l_1 = \sqrt[3]{\frac{3}{4\pi} V_1} + \sqrt{\left(\frac{3 \cdot V_1}{4\pi}\right)^{\frac{2}{3}} - \frac{A_{1,2}}{\pi}} \tag{24}$$

The area $A_{ij}$ depends on the radii $r_i, r_j$ of the two droplets interfacing and on the contact angle $\theta$. The contact angle is determined by the monolayer surface tension and the bilayer interface tension, and we assume it to depend on the lipids and oil composition, therefore to be constant. The volume of the droplets is known (from our experiments, the distribution is $V = 1.2 \pm 0.42$ nL, Supplementary Figure 11e, f). The radius of the droplets is $r_i = \sqrt[3]{\frac{4}{3\pi} V_i}$. The geometry of the bilayer is represented in Supplementary Figure 11a. The law of cosines gives on the one hand $(a_i + a_j)^2 = r_i^2 + r_j^2 + 2 \cdot r_i \cdot r_j \cdot \cos(2\theta)$, and the Pythagorean theorem gives on the other hand $(a_i + a_j)^2 = (\sqrt{r_i^2 - r_b^2} + \sqrt{r_j^2 - r_b^2})^2$, which leads to $r_b^2 - \sqrt{r_i^2 - r_b^2} \cdot \sqrt{r_j^2 - r_b^2} + r_i \cdot r_j \cdot \cos(2\theta) = 0$. This equation has a unique positive solution, from which the area $A_{ij}$ is calculated:

$$A_{ij} = \pi \frac{r_i^2 \cdot r_j^2 \cdot sin^2(2\theta)}{r_i^2 + r_j^2 + 2 \cdot r_i \cdot r_j \cdot \cos(2\theta)} \tag{25}$$

We measured the distribution of the bilayer area in our experiments, $A = 8.6 \pm 1.6 \cdot 10^3$ μm$^2$ (Supplementary Figure 11c), compared it to the distribution calculated from the droplet volume's distribution and Equation (25), as shown in Supplementary Figure 11b, and estimated $\theta \approx 42°$. This fitting method was preferred to directly taking the measured contact angle, as the measurement of the bilayer area was considered more accurate than the measurement of the contact angle.

It should be noted that the geometry of the bilayer changes during the run of a typical experiment: the contact angle $\theta$ decreases and consequently so does the area $A_{ij}$ (Supplementary Figure 11c, d). We assume that this is due to denaturation of proteins from the cell-extract at the water-oil interface, which changes the interfacial tension and in turn the contact angle. Additionally, evaporation causes the droplet volumes to decrease slightly (Supplementary Figure 11f).

#### 3.2 Protein expression, repression and induction

##### 3.2.1 Protein expression

We model transcription-translation of a protein $P$ from an active DNA template $D_{active}$ as a linear amplification with factor $\alpha_{max}$, with an initial delay due to the expression of the first RNA and its translation:

$$\frac{d[P]}{dt} = \alpha_{max} \cdot a_P \cdot (1 - e^{-t/t_{RNA}}) \cdot [D_{active}] \tag{26}$$

The lifetime of the RNA $t_{RNA}$ is typically 15 min in this cell-extract[22]. We assume $\alpha_{max}$ to be around 6 nM.min$^{-1}$ per nM of 1,000-nucleotides-length DNA template. We multiply this expression by a factor $a_P$ taking into consideration the DNA template concentration, the length of the protein and the strength of the ribosome-binding site (RBS) normalized to 40,000 a.u. (see Supplementary Section 4.3). For example, for 3 nM of a LacI DNA template of gene length 1,077 nucleotides and RBS strength 40,380, the factor will be $a_{LacI} = 3 \cdot \frac{1000}{1077} \cdot \frac{40380}{40000} \approx 2.81$.

To account for the degradation of the cell-extract and the depletion of resources, we model an exponential decay of expression over a time $t_2$ occurring after a delay $t_1$, so that for $t < t_1$, the expression is determined by Equation (26), and for $t > t_1$:

$$\frac{d[P]}{dt} = \alpha_{max} \cdot a_P \cdot (1 - e^{-t/t_{RNA}}) \cdot e^{-(t-t_1)/t_2} \cdot [D_{active}] \tag{27}$$

For fluorescent proteins, the maturation of the chromophore (characteristic time $t_{mat}$) is also taken into consideration. The mature, fluorescent protein is denoted $P_{mat}$:

$$\frac{d[P]}{dt} = \alpha_{max} \cdot a_P \cdot \left(1 - e^{-t/t_{RNA}}\right) \cdot [D_{active}] - \frac{[P]}{t_{mat}} \tag{28}$$

$$\frac{d[P_{mat}]}{dt} = \frac{[P]}{t_{mat}} \tag{29}$$

##### 3.2.2 Repression and induction

The repression of transcription by a protein $R$ of active concentration $[R_{active}]$ affects the concentration of active DNA following a Hill function with dissociation constant $K_{d,R}$ and cooperativity $n_R$:

$$[D_{active}] = \frac{{K_{d,R}}^{n_R}}{[R_{active}]^{n_R} + {K_{d,R}}^{n_R}} \tag{30}$$

Despite the presence of repressor proteins, transcription can still be leaky. Proteins in our system were found to have a basal expression even in presence of a large excess of repressor proteins. We tested several approaches to model this leaky expression and found that our simulations approached most closely experimental behavior with a constant leaky expression $\alpha_0$ of about 5 % maximum expression:

$$\frac{d[P]}{dt} = \alpha_{max} \cdot a_P \cdot (1 - e^{-t/t_{RNA}}) \cdot (\alpha_0 + [D_{active}]) \tag{31}$$

In our system, the repression of transcription by proteins $(R)$ such as LacI and TetR is lifted by small inducer molecules $(I)$, respectively IPTG and anhydrotetracycline. The binding of such small molecules to the proteins renders them inactive. We model the kinetic steps of this binding as follows:

$$[R_{active}] + I \underset{k_{off}}{\overset{k_{on}}{\rightleftarrows}} [R_{inactive}] \tag{32}$$

$$\frac{d[R_{active}]}{dt} = \frac{d[I]}{dt} = -\frac{d[R_{inactive}]}{dt} = -k_{on} \cdot [R_{active}] \cdot [I] + k_{off} \cdot [R_{inactive}] \tag{33}$$

##### 3.2.3 Protein degradation and life-time of cell-extract

Finally, active degradation of tagged proteins by ClpX was modeled as zeroth-order kinetics[22] with a rate $k_{deg}$ estimated at 1 nM.min$^{-1}$. Other studies have found zeroth-order degradation rates of around 10 to 15 nM.min$^{-1}$ for protein concentrations of 0.1 µM and above[22,23], while some have shown a protein-concentration-dependent degradation rate which was at maximum around 10 nM.min$^{-1}$ for 10 µM protein[24]. To account for protein-concentration-dependent degradation at low protein concentrations, and to avoid that degradation would be stronger than expression and cause negative concentration values, we multiplied the degradation term with a Hill function assuming an activation of the degradation with a $K_{d,deg}$ estimated as the concentration of the ClpX protein, around 1 nM[22]. We also assumed that protein degradation in the cell-extract is affected by resource depletion[24]. The degradation of a protein is therefore modeled as follows for $t > t_1$:

$$\frac{d[P]}{dt} = -k_{deg} \frac{[P]}{K_{d,deg} + [P]} e^{-(t-t_1)/t_2} \tag{34}$$

For $t < t_1$, the exponential term is absent.

##### 3.2.4 Protein and fluorescence intensity

For fitting purposes in bulk measurements, we relate the measured fluorescence measurements to the concentration of the proteins with a linear dependency:

$$FI = \alpha \cdot [P_{mat}] + \beta \tag{35}$$

The dependency of fluorescence to protein concentration in droplet experiments is discussed in detail in Supplementary Section 2.8.2.

#### 3.3 Reaction-diffusion model

The full reaction-diffusion model of the circuit consists of the following set of ordinary differential equations (for topology A, other topologies are modelled accordingly), in any compartment $i$ with two neighbors (the diffusion term adapts accordingly for the first and last compartments), for $t > t_1$. $L$ is LacI ($K_{d,L}$ and $n_L$ to pLacO), $I$ is IPTG, $LI$ is the LacI-IPTG complex, $R$ is RFP, $R_{mat}$ is mature RFP, $T$ is TetR ($K_{d,T}$ and $n_T$ to pTetO), $Y$ is YFP, $Y_{mat}$ is mature YFP.

$$\begin{aligned}\frac{d[I]_i}{dt} = &-\frac{D \cdot P}{V_i(D + P \cdot l_i)}\left(A_{i-1,i} \cdot (n_i - n_{i-1}) + A_{i,i+1} \cdot (n_i - n_{i+1})\right) \\ &-k_{on} \cdot [L]_i \cdot [I]_i + k_{off} \cdot [LI]_i\end{aligned} \tag{36}$$

$$\begin{aligned}\frac{d[L]_i}{dt} = &\alpha_{max} \cdot a_L \cdot \left(1 - e^{-t/t_{RNA}}\right) \cdot e^{-(t-t_1)/t_2} \cdot \left(\alpha_0 + \frac{{K_{d,T}}^{n_T}}{[T]^{n_T} + {K_{d,T}}^{n_T}}\right) \\ &-k_{deg} \frac{[L]}{K_{d,deg} + [L]} e^{-(t-t_1)/t_2} - k_{on} \cdot [L]_i \cdot [I]_i + k_{off} \cdot [LI]_i\end{aligned} \tag{37}$$

$$\frac{d[LI]_i}{dt} = -k_{deg} \frac{[LI]}{K_{d,deg} + [LI]} e^{-(t-t_1)/t_2} + k_{on} \cdot [L]_i \cdot [I]_i - k_{off} \cdot [LI]_i \tag{38}$$

$$\begin{aligned}\frac{d[R]_i}{dt} = &\alpha_{max} \cdot a_R \cdot \left(1 - e^{-t/t_{RNA}}\right) \cdot e^{-(t-t_1)/t_2} \cdot \left(\alpha_0 + \frac{{K_{d,T}}^{n_T}}{[T]^{n_T} + {K_{d,T}}^{n_T}}\right) \\ &-k_{deg} \frac{[R]}{K_{d,deg} + [R]} e^{-(t-t_1)/t_2} - \frac{[R]}{t_{mat,R}}\end{aligned} \tag{39}$$

$$\frac{d[R_{mat}]_i}{dt} = \frac{[R]}{t_{mat,R}} - k_{deg} \frac{[R_{mat}]}{K_{d,deg} + [R_{mat}]} e^{-(t-t_1)/t_2} \tag{40}$$

$$\frac{d[T]_i}{dt} = \alpha_{max} \cdot a_T \cdot \left(1 - e^{-t/t_{RNA}}\right) \cdot e^{-(t-t_1)/t_2} \cdot \left(\alpha_0 + \frac{{K_{d,L}}^{n_L}}{[L]^{n_L} + {K_{d,L}}^{n_L}}\right) \tag{41}$$

$$\frac{d[Y]_i}{dt} = \alpha_{max} \cdot a_Y \cdot \left(1 - e^{-t/t_{RNA}}\right) \cdot e^{-(t-t_1)/t_2} \cdot \left(\alpha_0 + \frac{{K_{d,L}}^{n_L}}{[L]^{n_L} + {K_{d,L}}^{n_L}}\right) - \frac{[Y]}{t_{mat,Y}} \tag{42}$$

$$\frac{d[Y_{mat}]_i}{dt} = \frac{[Y]}{t_{mat,Y}} \tag{43}$$

##### 3.4 Noise

Noise was modelled using a Monte Carlo method. For each potential noise source, a Gaussian distribution of the corresponding model parameter was determined. For example, droplet volume was estimated for experimental data to follow a Gaussian distribution with mean 1.2 nL and standard deviation 0.42 nL. The distribution of the biochemical species was determined by running a given number of simulations (typically 500) where the noise parameter was randomly sampled from the Gaussian distribution. Note that we do not consider stochastic effects due to the large number of molecules in one droplet ($1\ \mathrm{nM} \cdot 1\ \mathrm{nL} \cdot N_A \approx 10^6$).

#### 4 Supplementary experiments

##### 4.1 Toxicity of reference dye

The purified blue-fluorescent protein mTurquoise2 was used as a reference dye to normalize the fluorescence in droplets (see Supplementary Section 2.8.2). The toxicity of mTq2, more specifically of its storage buffer, was tested in order to determine the optimal concentration for experiments. 0 nM, 265 nM, 530 nM, 2.65 or 5.3 μM purified mTq2 was added to a bulk cell-extract experiment of topology A with either 0 mM IPTG (strong LacI-RFP expression) or 10 mM IPTG (strong TetR-YFP expression). As shown in Supplementary Figure 12, 265 nM and 530 nM of mTq2 did not significantly affect protein expression, whereas higher concentrations did. To minimize any effect on the function of the circuit, 265 nM of mTq2 was chosen as the reference dye in all droplets experiments.

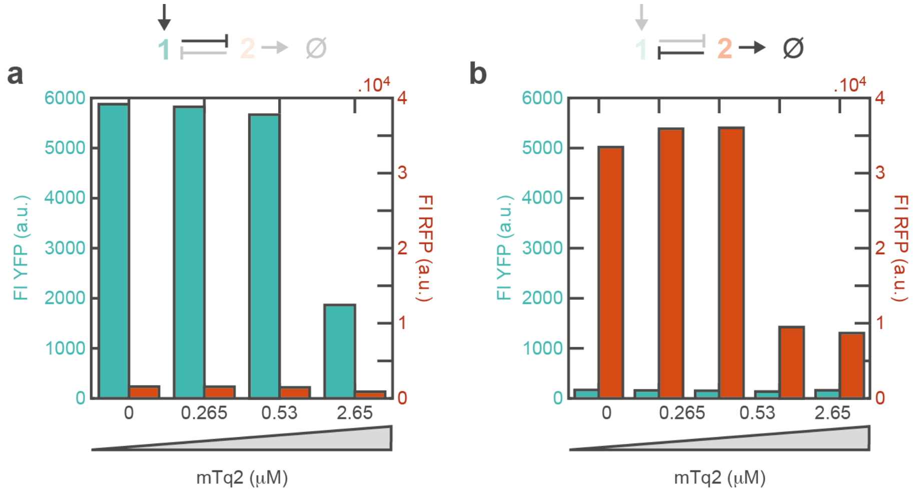

**Supplementary Figure 12 | Effect of reference dye on circuit expression.** Topology A is expressed in bulk, with (**a**) or without (**b**) 10 mM IPTG, with different concentrations of mTq2, the reference dye.

#### 4.2 Fitting of model parameters

##### 4.2.1 Repression constants

Bulk screening experiments were conducted to determine the repression constants, namely $K_{d,R}$ and $n$, for the repression of the pLacO promoter by LacI and the repression of the pTetO promoter by TetR. To this end, titrations of LacI (respectively TetR) in ranges from 1 nM to 6 µM (respectively 10 nM to 3 µM) were conducted in cell-extract against 5 nM of reporter plasmids (pSB1A3-pLacO-GFP, respectively pSB1A3-I13521 that consists of pTetO-mRFP). The final protein levels at 16 h (LacI) or 10 h (TetR) were used to fit the repression model, with the following parameters:

- $\alpha$ = 13360 (Supplementary Figure 13a) and 510 (Supplementary Figure 13b)
- $\beta$ = 5279 (Supplementary Figure 13a) and 110 (Supplementary Figure 13b)

The titration of LacI against pSB1A3-pLacO-GFP (Supplementary Figure 13a) gave the following fitted values: $K_{d,L} = 2.9 \cdot 10^{-8}$ M, $n_L$ = 1.5, squared correlation coefficient $R^2$ = 0.99.

The titration of TetR against pSB1A3-I13521 (Supplementary Figure 13b) gave the following fitted values: $K_{d,T} = 1.3 \cdot 10^{-7}$ M and $n_T$ = 4.3, squared correlation coefficient $R^2$ = 0.9936.

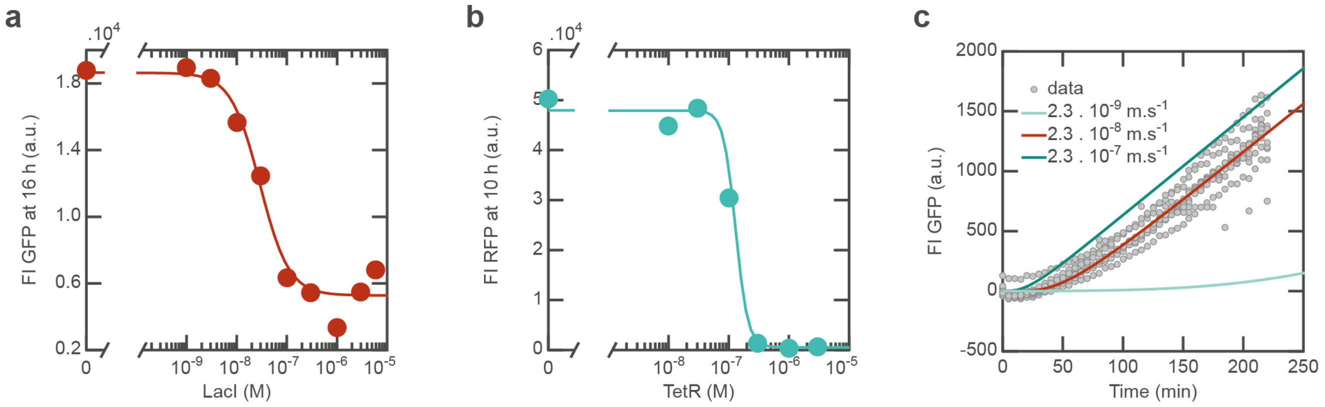

**Supplementary Figure 13 | Fit of titration of repressor proteins and bilayer permeability to IPTG. a-b,** Titration of repressor proteins and fit of repression constants. Final fluorescence intensities are taken and fitted with a Hill function. **a**, LacI is titrated against pLacO-GFP (5 nM plasmid). **b**, TetR is titrated against pTetO-RFP (I13521, 5 nM plasmid). **c,** Induction of GFP expression by diffusing IPTG (3 mM) in a droplet assembly and fit of bilayer permeability to IPTG. Mean intensities of the pLacO-GFP receiver droplets in 10 assemblies are plotted as grey dots. The permeability fit ($P = 2.3 \cdot 10^{-8}$ m s$^{-1}$) to the data is plotted in red line, as well as 2 simulations with different permeability values ($P = 2.3 \cdot 10^{-7}$ m s$^{-1}$, dark blue line and $P = 2.3 \cdot 10^{-9}$ m s$^{-1}$, light blue line).

##### 4.2.2 Permeability of bilayers to IPTG

The permeability of lipid bilayers to IPTG was characterized through the following experiment. A receiver droplet contained 5 nM pSB1A3-pLacO-GFP, 100 nM LacI. It was connected to a buffer droplet, which contained 5 µM Atto655-DNA as a marker and 1.5 µM α-hemolysin (α-HL) pore (this was used in a control experiment to confirm that IPTG diffusion was not pore-mediated and has no influence on diffusion kinetics). A sender droplet containing 3 mM IPTG, 1.5 µM α-HL was then connected to the buffer droplet. All droplets contained 75 % v/v extract and buffer. The model of LacI repression of the pLacO promoter lifted by IPTG binding to LacI was used to fit the time-fluorescence data in the receiver. Cell-extract degradation was not taken into account, as only the first 200 minutes of expression were used for fitting, in which cell-extract degradation is neglected. The parameters used are summarized in Supplementary Table 4**.**

**Supplementary Table 4 | Parameters used in bulk fitting.**

| Parameter | Value | Description |
|---|---|---|
| $A_{ij}$ | $8.21 \cdot 10^{-9}$ m$^2$ | effective cross-section for transport (calculated from measured droplet radii as described in the Supplementary Section 3.1.2) |
| $\alpha_{max}$ | $0.500 \cdot 10^{-9}$ M s$^{-1}$ | expression factor for 5 nM DNA template |
| $D$ | $7.44 \cdot 10^{-10}$ m$^2$ s$^{-1}$ | diffusion coefficient of IPTG in water (calculated with the Stokes-Einstein equation from the molecular modelling of the radius of IPTG, see Dupin (2019)[6]) |
| $l$ | $93.7 \cdot 10^{-6}$ m | average diameter of the droplets |
| $V$ | $1.90 \cdot 10^{-9}$ L | average volume of the droplets |
| $K_{d,LacI}$ | $2.9 \cdot 10^{-8}$ M | repression constant of LacI on pLacO (determined with bulk titration of LacI, see Supplementary Figure 12a) |
| $n$ | 1.5 | Hill factor of LacI repression on pLacO (determined with bulk titration of LacI, see Supplementary Figure 12b) |
| $k_{on}$ | $7.2 \cdot 10^6$ M$^{-1}$ min$^{-1}$ | on-rate of LacI binding to IPTG[25] |
| $k_{off}$ | 12.6 min$^{-1}$ | off-rate of LacI binding to IPTG[25] |
| $[I]_0$ | $3 \cdot 10^{-3}$ M | initial concentration of IPTG |
| $[R]_0$ | $100 \cdot 10^{-9}$ M | initial concentration of LacI |
| $t_{RNA}$ | 15 min | lifetime of the RNA[22] |
| $t_{mat}$ | 7 min | maturation time of GFP[26] |

The fitted parameters were $\alpha$ (fitted to 276) and $\beta$ (fitted to 0), the fluorescence intensity to protein concentration factors, and the permeability $P$. The permeability of lipid bilayers to IPTG was fitted to $2.26 \cdot 10^{-8}$ m s$^{-1}$ (Supplementary Figure 13c). In a previous characterization of the diffusion regimes in the droplet assemblies[6], we determined that if the permeability of the bilayers to the signal is lower than $10^{-7}$ m s$^{-1}$, the diffusion of the signal is limited by the permeation of the bilayer. We can therefore approximate that the IPTG concentration is instantaneously homogenized inside the droplets, and that what limits IPTG diffusion is the number of bilayers it encounters. This is confirmed by the experiment shown in Supplementary Figure 18, where the protein expression gradient is similar in assemblies of 5, 7 or 9 receivers: the expression is differentiated over the 5 first droplets, and in a low GFP-high RFP state for all receivers further from the sender droplet.

#### 4.3 Simulations of bulk response

Simulations of the 3 topologies in bulk was used to fit the experimental behavior and to optimize model parameters. The circuit was simulated for 10 h of expression (10 min intervals) at different IPTG concentrations (0 µM and 100 values distributed logarithmically between 0.1 µM and 10 mM).

First, the effect of model parameters on circuit response was systematically screened (Supplementary Figure 14). Based on this, protein expression was readjusted for each protein by a factor ($a_{protein}$), taking into account the template DNA concentration (normalized to 1 nM), the strength of its ribosome binding site (calculated with the RBS Calculator v2.1 (De Novo DNA)[27], available at https://www.denovodna.com/software, normalized to 40,000 a.u.), and the length of the protein (normalized to 1,000 bp, 333 amino acids). Decay parameters of the cell-extract activity ($t_1, t_2$) were adjusted to fit experimental time curves.

The response of the circuit is particularly strongly affected by the ratio of protein expression to protein degradation rates. Therefore, the effects of parameters $\alpha_{max}$ and $k_{deg}$ were investigated further: both parameters were simultaneously varied over 1 order of magnitude and the resulting fitted circuit thresholds were compared to the experimental values obtained in bulk (Supplementary Figure 15). From this, the values for parameters $\alpha_{max}$ and $k_{deg}$ were optimized to fit experimental response (Supplementary Figure 16).

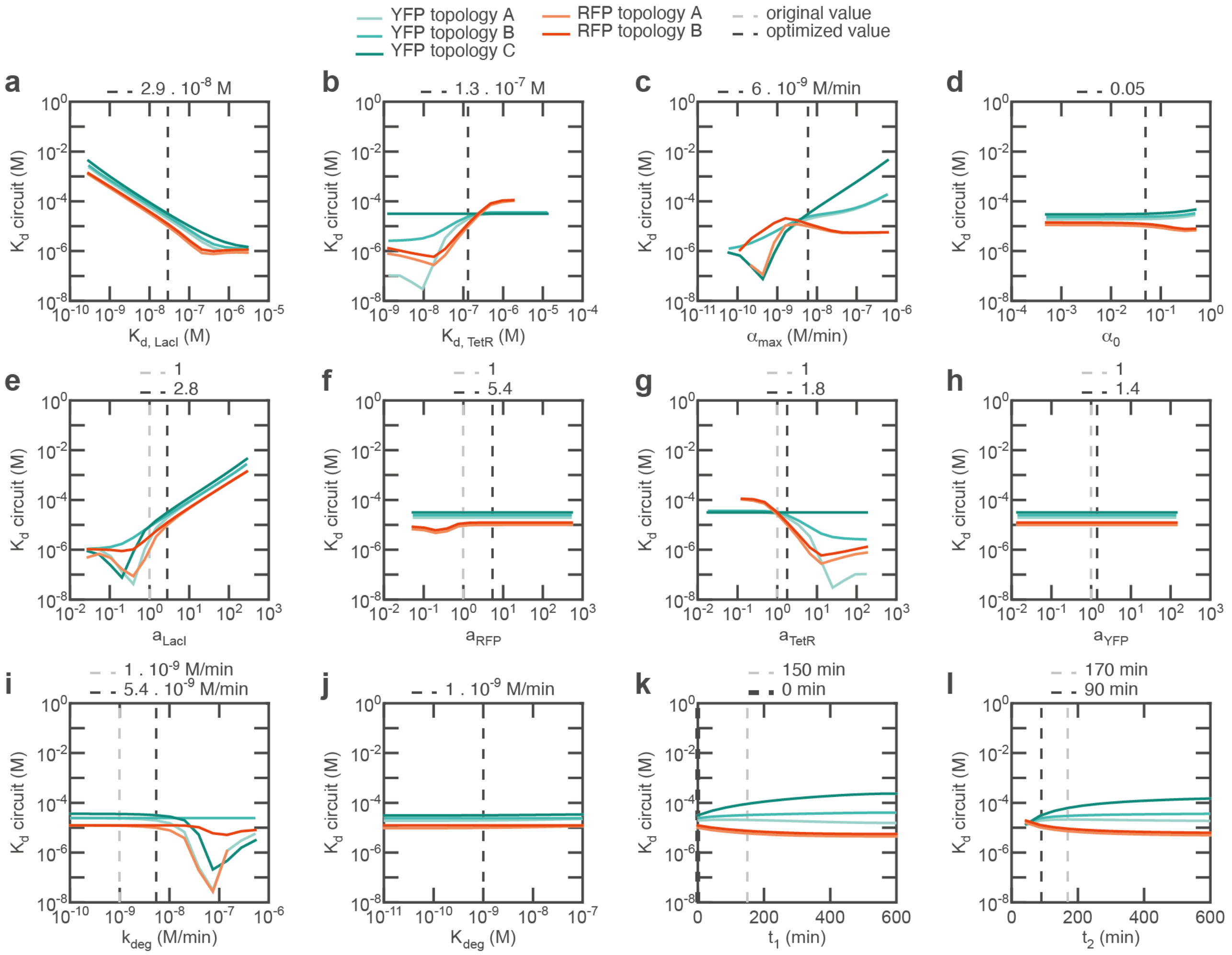

**Supplementary Figure 14 | Parameter effect on bulk response.** All model parameters were kept constant at their original value except the one which was varied, and the circuit was simulated for different IPTG concentrations. The resulting end-point protein concentrations for each IPTG concentration were used to fit the $K_d$ of YFP induction (blue lines) and RFP repression (red lines) with a Hill function. This was done for all three topologies (light colour: topology A, middle colour: topology B, dark colour: topology C). The parameter values were screened logarithmically in four orders of magnitude centered around their original value in the simulations, except for $t_1$ and $t_2$, which were screened linearly between 0 min and 600 min. The fitted $K_d$ values for each topology are plotted against the parameter values. The original value estimated or calculated for each parameter is indicated by a vertical light grey dashed line. The optimized value used in final simulations is indicated by a vertical dark grey dashed line. **a**, $K_{d,L}$. **b**, $K_{d,L}$. **c**, $\alpha_{max}$. **d**, $\alpha_0$. **e**, $a_{LacI}$. **f**, $a_{RFP}$. **g**, $a_{TetR}$. **h**, $a_{YFP}$. **i**, $k_{deg}$. **j**, $K_{deg}$. **k**, $t_1$. **l**, $t_2$.

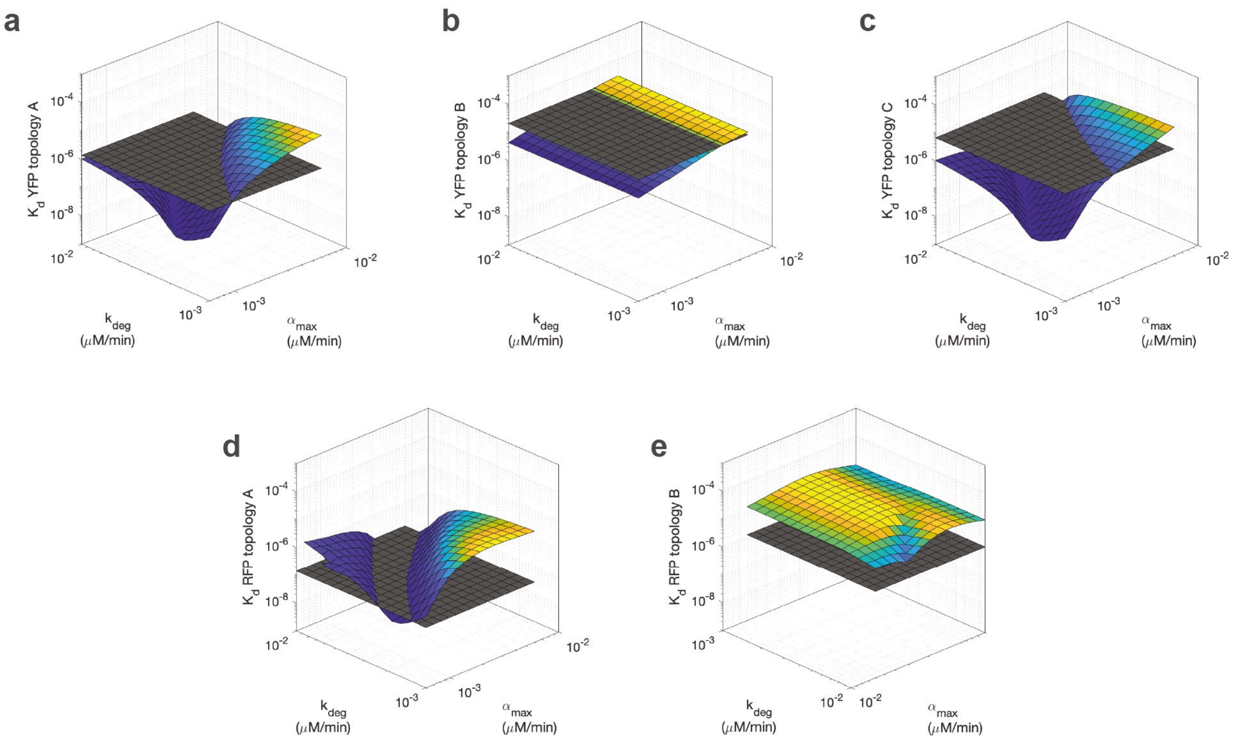

**Supplementary Figure 15 | Effect of expression and degradation kinetics on bulk response.** The method was identical to Supplementary Figure 14, except that both $\alpha_{max}$ and $k_{deg}$ were simultaneously screened (logarithmically, one order of magnitude). This was done for all three topologies. The $K_d$ of YFP induction or RFP repression was fitted. The graphs represent the fitted $K_d$ values (colour) and the experimental value (grey). **a,** topology A YFP. **b,** topology B YFP. **c,** topology C YFP. **d,** topology A RFP. **e,** topology B RFP.

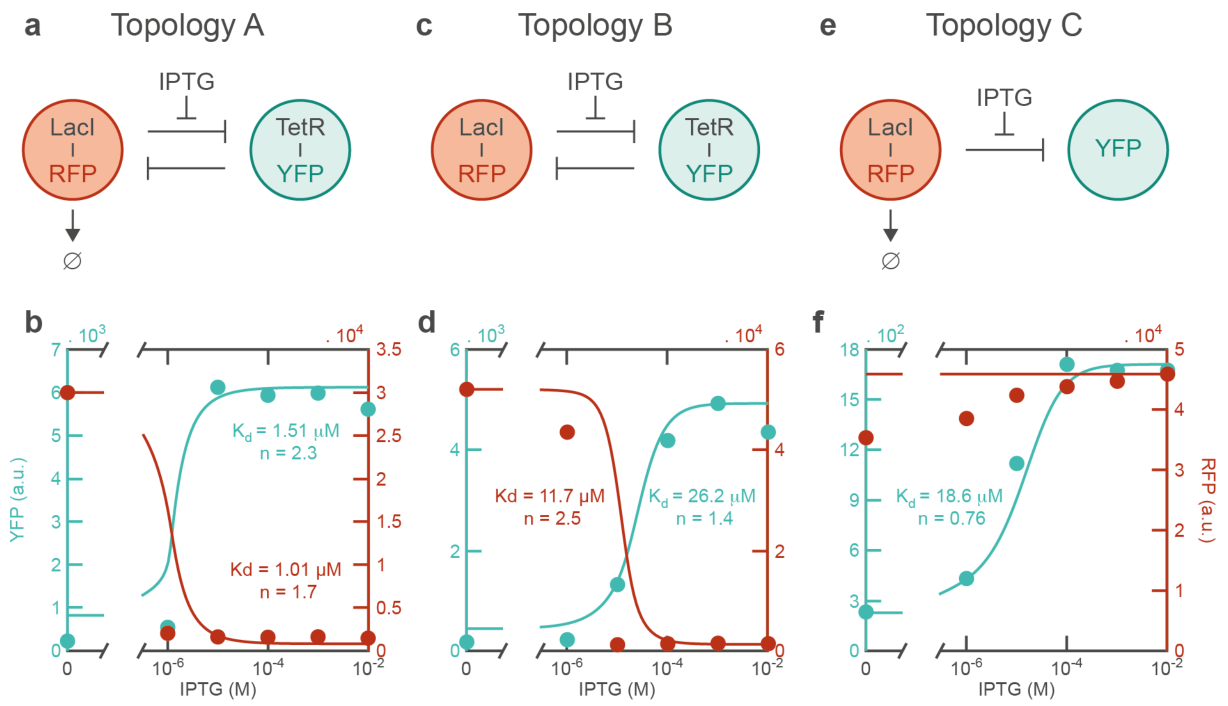

**Supplementary Figure 16 | Topologies and their transfer function in bulk. a, c, e**, topologies of the circuit. **b, d, f,** corresponding transfer functions. The simulated final protein levels are plotted (lines) against the IPTG concentration, normalized to the experiment values (dots, blue: YFP, red: RFP). These slopes are fitted with a Hill function, and the fitted $K_d$ and $n$ are noted in corresponding colours.

#### 4.4 Characterization in droplets

The circuit was characterized in droplets, first by monitoring the effect of IPTG and aTc diffusing gradients on circuit response and assembly differentiation (Supplementary Figure 17). As an aTc source droplet led to a relatively homogeneous protein expression response across the assembly, an IPTG source droplet, with a diffusing gradient established over a shorter distance and therefore a differentiated induced protein expression, was preferred. Second, the effect of the length of the assemblies was characterized (Supplementary Figure 18). The model predicts that, since bilayer permeation is the limiting step in IPTG diffusion (Supplementary Section 4.2.2, Supplementary Figure 13c), IPTG will only diffuse into the first few droplets, and further droplets in the assembly will have an undifferentiated, "off" expression profile. This is confirmed by experiments, where differentiation occurs over the first 5 droplets independently of the length of the assembly.

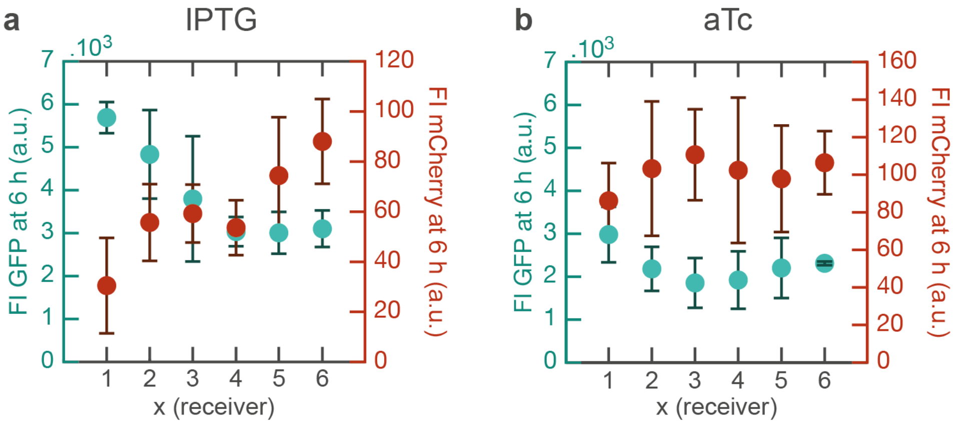

**Supplementary Figure 17 | Diffusion of 2 inducer signals and response of the circuit.** Signals diffuse from a sender droplet into an area of 6 identical receivers containing 5 nM pSB1C3-pLacO-TetR-GFP and 5 nM pSB1A3-pTetO-LacI-mCherry (equivalent to topology B) in cell-extract. Circles and error bars represent the mean and standard deviation over 7 assemblies of the GFP and RFP droplet mean intensities at 6 h. a, Signal is 10 mM IPTG. The differentiated levels of protein expression between the receivers indicate that the characteristic length of diffusion of IPTG during the course of the experiment matches the length of the assembly. b, Signal is 10 µM aTc. The homogeneous levels of protein expression with high RFP and low GFP (corresponding to induction by aTc) indicate that aTc diffuses quickly compared to the time of the experiment and that its concentration rapidly equilibrates over the length of the assembly.

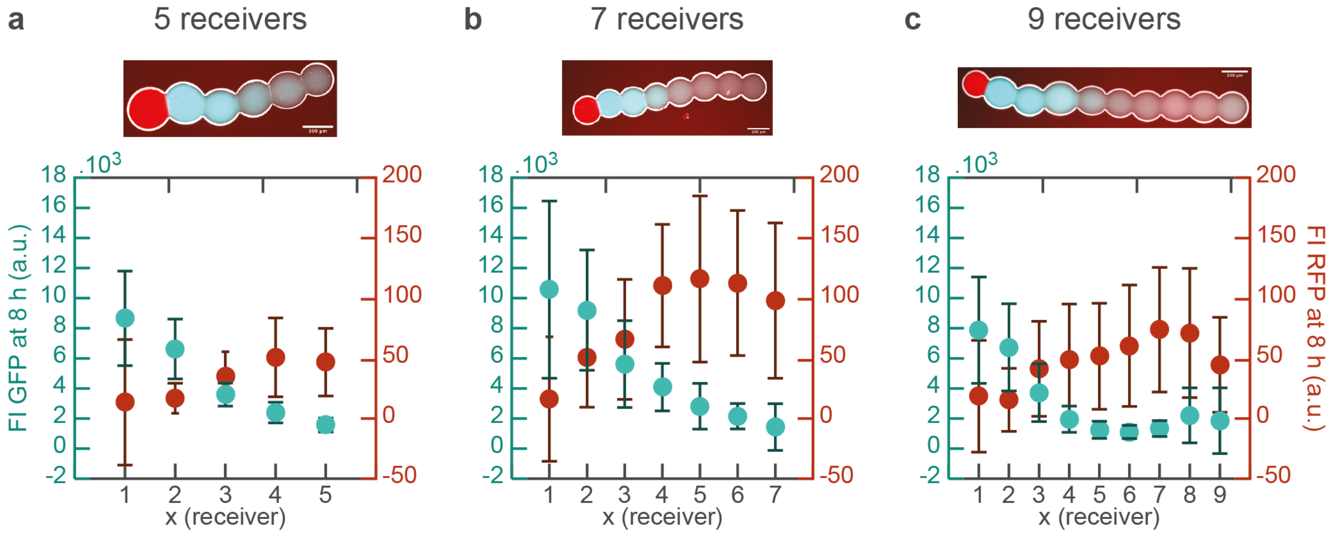

**Supplementary Figure 18 | Dependence of the circuit response on the number of receivers.** A sender droplet containing 10 mM IPTG (marker: 1 µM Atto655-DNA) is connected to an array of 5 (**a**), 7 (**b**) or 9 (**c**) receivers, containing 5 nM pSB1C3-pLacO-TetR-GFP and 5 nM pSB1A3-pTetO-LacI-mCherry (equivalent to topology B) in cell-extract. Circles and error bars represent the mean and standard deviation over 10 assemblies of the GFP and RFP droplet mean intensities at 8 h. Microscopy images show an overlay of brightfield (white), GFP (blue) and RFP (red) images of a representative assembly at 8 h. Scale bar: 100 µm.

#### 4.5 Simulations of reaction-diffusion circuit

Simulations of the reaction-diffusion circuit in the droplet assemblies allowed insights into the spatial differentiation of protein expression and the timing of cell-extract activity versus signal gradient. Additionally, the modelling of noise brought evidence that the assembly-to-assembly variability is principally due to noise in protein expression, and to a lesser extent to noise in droplet volume (Supplementary Figure 19). By modelling accurately the noise in our system, we were able to simulate the positional information (PI) given by each protein profile as well as the joint positional information, for each of the 3 circuit topologies and each inducer initial concentration (Supplementary Figure 22). The effect of model parameters on the PI was systematically screened (Supplementary Figure 20). The dependance of the PI on both the dynamics of the circuit and the diffusion profile of the signal was characterized for all 3 topologies (Supplementary Figure 21).

The circuit was simulated for 8 h of expression (10 min intervals), with 1 sender and 5 receivers. For each condition, $n$ = 500 assemblies were simulated unless indicated otherwise. The sources of noise were either droplet volume, protein expression or both. The volume $V$ of each of the droplets in each assembly was randomly sampled from a normal distribution with mean 1.2 nL and standard deviation 0.42 nL (when not varied, the volume of all droplets was fixed at 1.2 nL). The maximum protein expression $\alpha_{max}$ in each droplet and for each pair of proteins (TetR and YFP or LacI and RFP) was modulated by a factor randomly picked from a normal distribution with mean 1 and standard deviation 0.2 (when not varied, this factor was fixed at 1). The positional information (PI) was calculated with the second gaussian approximation (SGA) approach, and the values for the last 3 hours were averaged (PI is roughly fixed after 3-4 hours of expression), unless PI is plotted against time.

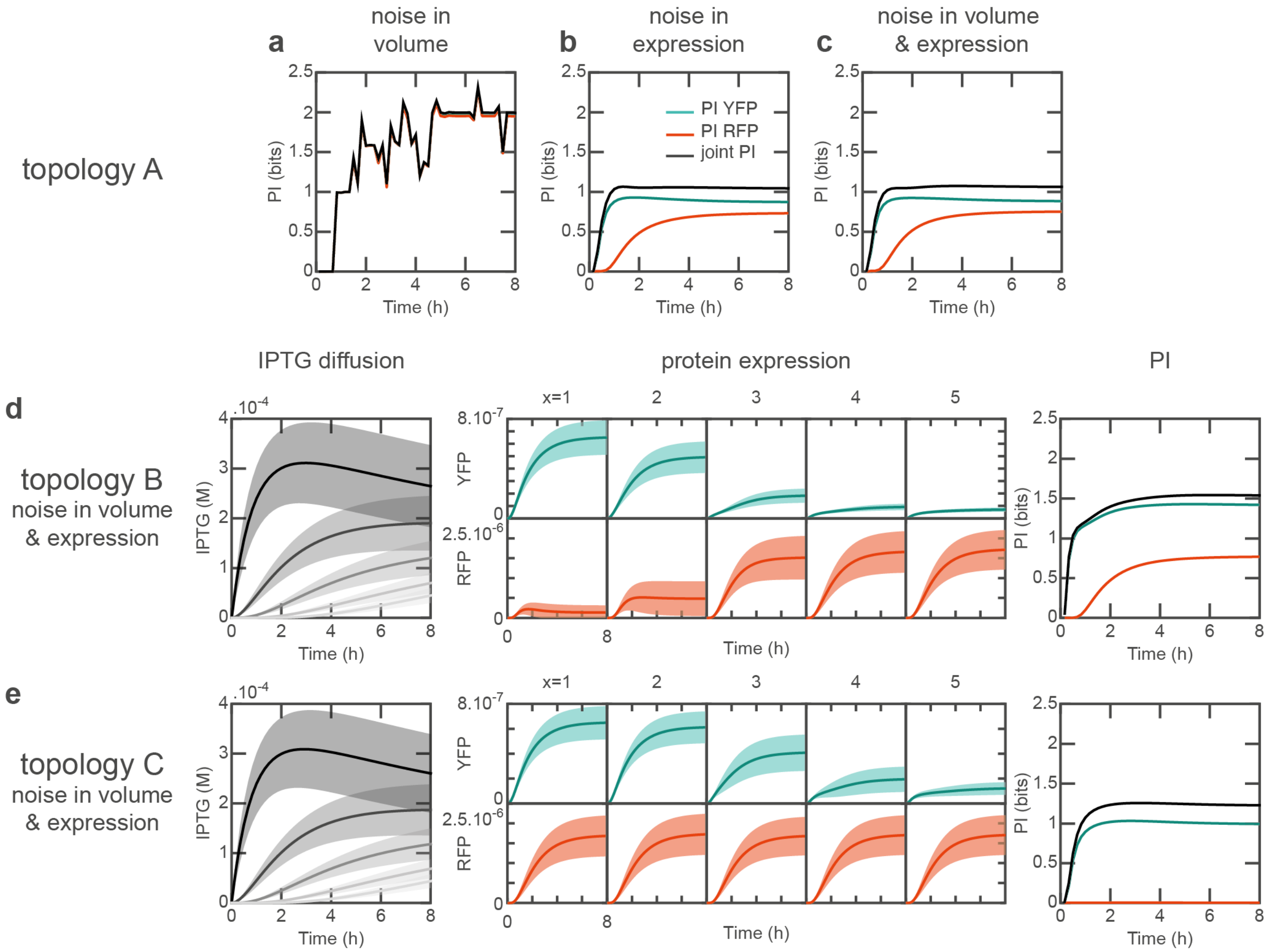

**Supplementary Figure 19 | Simulation of signal diffusion, protein expression and PI for all 3 topologies.** Initial IPTG concentration in the sender droplet: 1 mM. **a**, **b, c**, PI of topology A, noise in volume (**a**), protein expression (**b**), or both (**c**). **d, e**, IPTG (grey, left panel), YFP (blue, central panels top) and RFP (red, central panels bottom)

concentrations (mean: thick line, standard deviation: shaded area) and PI (right panel, blue: YFP, red: RFP, grey: joint) of topologies B (**d**, $n$ = 400 assemblies) and C (**e**) with noise sources: both volume and protein expression.

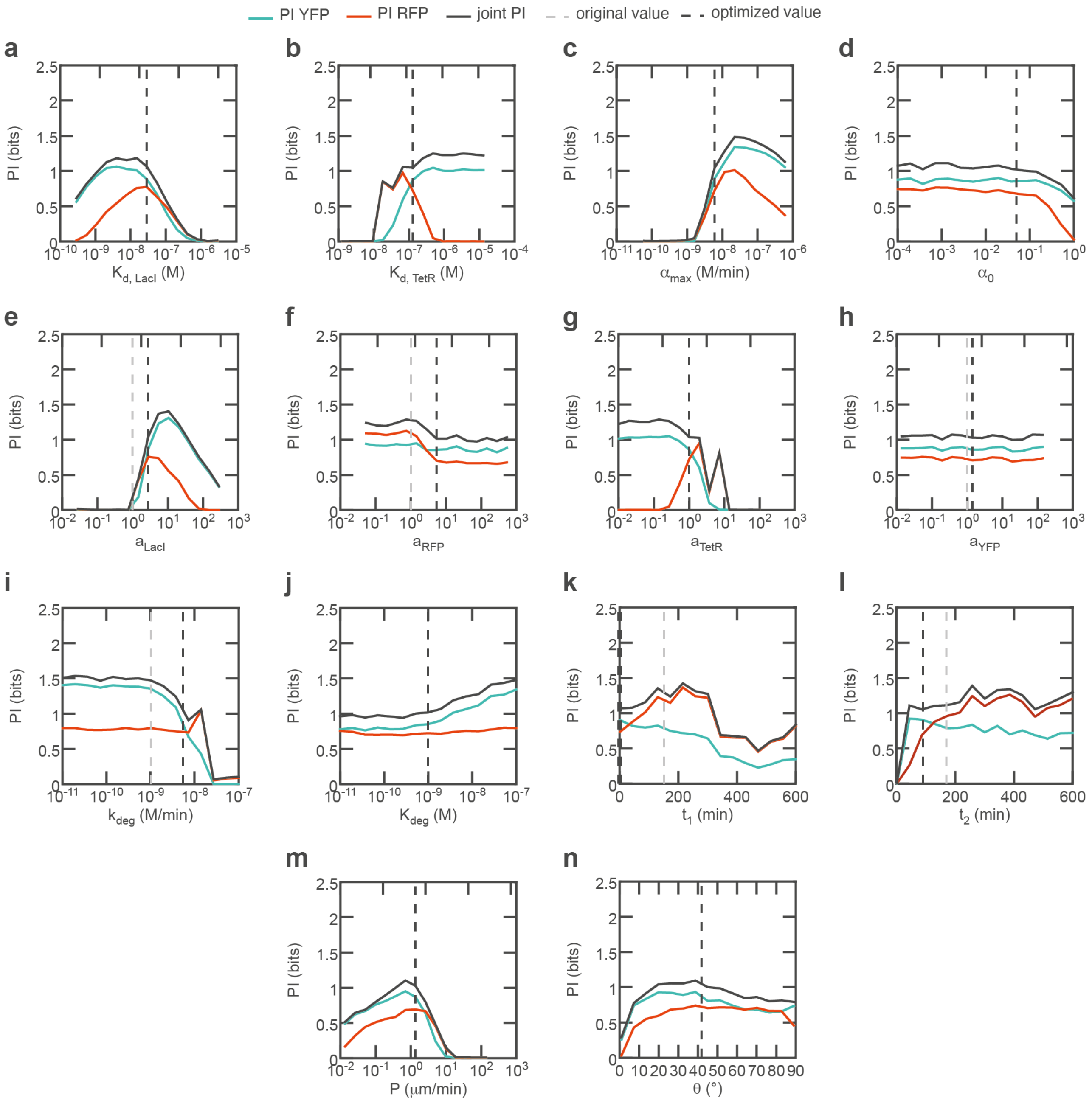

**Supplementary Figure 20 | Effect of model parameters on PI.** Topology A, initial IPTG concentration in the sender droplet: 1 mM. The parameter values were screened logarithmically in four orders of magnitude centered around their original value in the simulations, except for $t_1$, $t_2$ and $\theta$, which were screened linearly ($t_1$ and $t_2$: between 0 min and 600 min, $\theta$: between 1 and 89°). For each condition, $n$ = 500 assemblies were simulated, and the sources of noise were both volume and protein expression. Positional information calculated from the YFP profile (blue), the RFP profile (red), or the joint positional information (dark grey) are plotted against parameter values. The original value estimated or calculated for each parameter is indicated by a vertical light grey dashed line. The optimized value used in final simulations is indicated by a vertical dark grey dashed line. **a,** $K_{d,L}$. **b,** $K_{d,L}$. **c,** $\alpha_{max}$. **d,** $\alpha_0$. **e,** $a_{LacI}$. **f,** $a_{RFP}$. **g,** $a_{TetR}$. **h,** $a_{YFP}$. **i,** $k_{deg}$. **j,** $K_{deg}$. **k,** $t_1$. **l,** $t_2$. **m,** $P$. **n,** $\theta$.

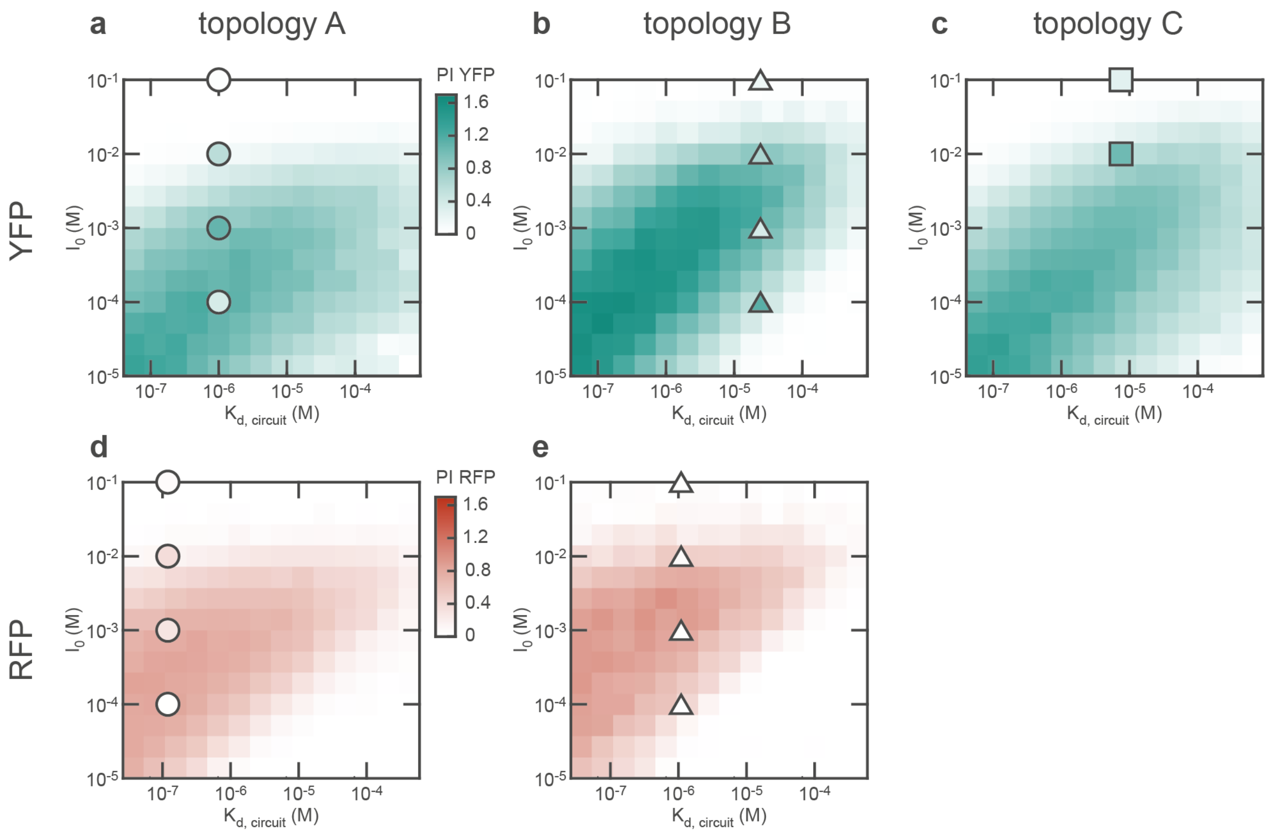

**Supplementary Figure 21 | PI maximization depends on the circuit's threshold and the initial inducer concentration.** The simulations were conducted for different pairs of IPTG concentrations (15 values distributed logarithmically between 10 μM and 100 mM) and circuit's threshold (varied by varying $K_{d,LacI}$ across 15 values distributed logarithmically across 4 orders of magnitude centered on its original value). The circuit's threshold depends on $K_{d,LacI}$ by a power law, determined and fitted from Supplementary Figure 14a. The sources of noise were both volume and protein expression. The circles, squares and triangles indicate the experimental data points for each topology and protein. **a-c,** YFP. **d-e,** RFP. **a, d,** topology A. **b, d,** topology B. **c,** topology C.

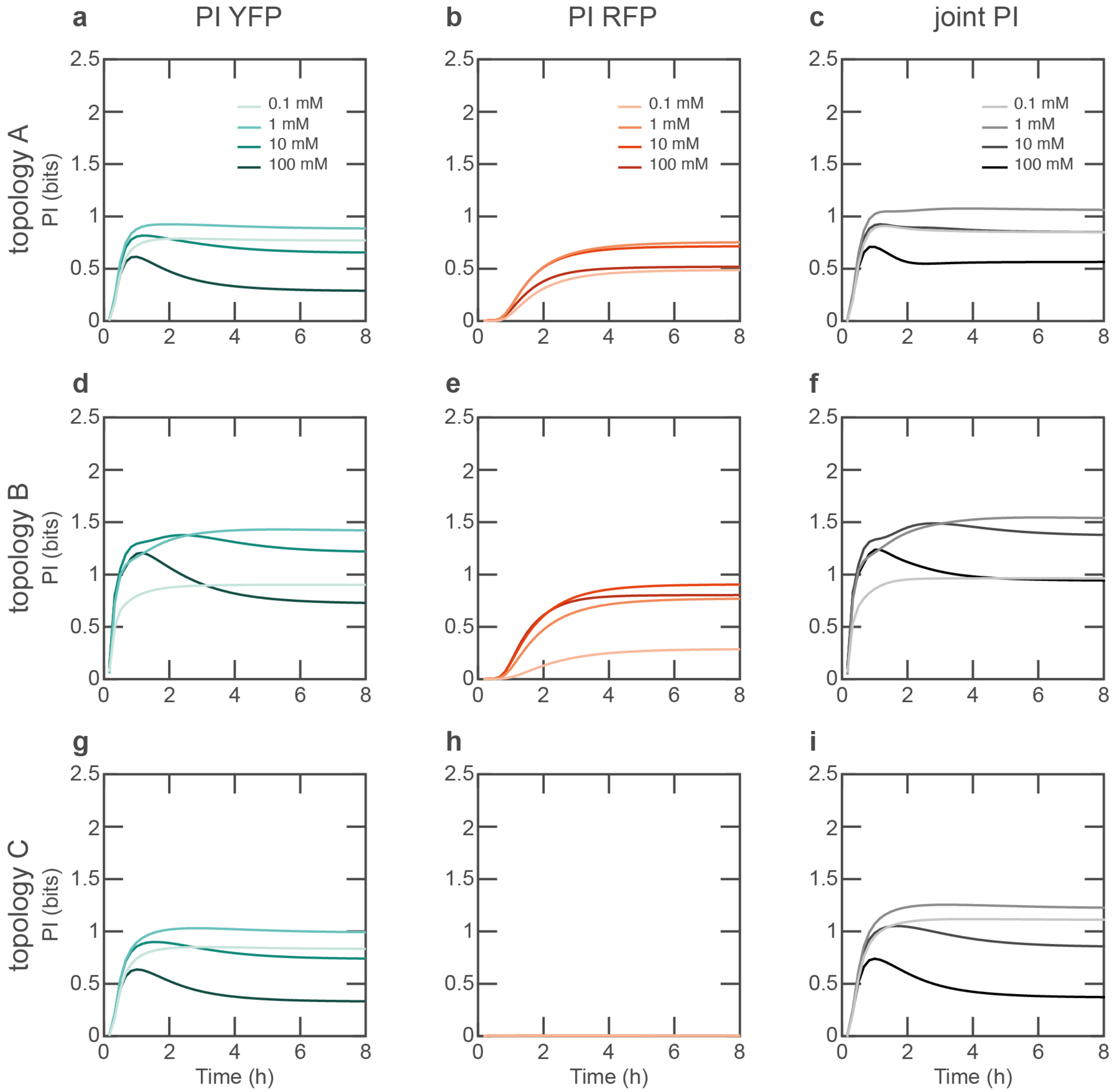

**Supplementary Figure 22 | Simulation of signal diffusion, protein expression and PI for all 3 topologies.** Initial IPTG concentration in the sender: 0.1 mM (light color), 1 mM (medium light color), 10 mM (medium dark color) or 100 mM (dark color). The sources of noise were both droplet volume and protein expression. **a-c,** PI of topology A. **d-f,** PI of topology B ($n$ = 400 assemblies). **g-i,** PI of topology C. **a**, **d**, **g**, PI of YFP. **b, e, h,** PI of RFP. **c, f, i,** joint PI.

#### 5 Sequences

For all double-stranded sequences, the non-template, (+), coding strand is given. The sequence of the insert is indicated, from the upstream EcoRI to the downstream PstI cloning sites.

**Atto655-DNA**

(Atto655- TTGATGTAT)

**pSB1A3-pLacO-GFP**

pLacO-1, RBS, GFPmut3 (BBa_E0040), terminator

plasmid map: https://benchling.com/s/seq-6XyH5h3vss0N7I7OIycL

GAATTCATTCTGATAAATGTGAGCGGATAACATTGACATTGTGAGCGGATAACAAGATACTGAGCACCTATGGACTATGTTTTCACATACGAGG
GGGATTAGATGCGTAAAGGAGAAGAACTTTTCACTGGAGTTGTCCCAATTCTTGTTGAATTAGATGGTGATGTTAATGGGCACAAATTTTCTGT
CAGTGGAGAGGGTGAAGGTGATGCAACATACGGAAAACTTACCCTTAAATTTATTTGCACTACTGGAAAACTACCTGTTCCATGGCCAACACTT
GTCACTACTTTCGGTTATGGTGTTCAATGCTTTGCGAGATACCCAGATCATATGAAACAGCATGACTTTTTCAAGAGTGCCATGCCCGAAGGTT
ATGTACAGGAAAGAACTATATTTTTCAAAGATGACGGGAACTACAAGACACGTGCTGAAGTCAAGTTTGAAGGTGATACCCTTGTTAATAGAAT
CGAGTTAAAAGGTATTGATTTTAAAGAAGATGGAAACATTCTTGGACACAAATTGGAATACAACTATAACTCACACAATGTATACATCATGGCAG
ACAAACAAAAGAATGGAATCAAAGTTAACTTCAAAATTAGACACAACATTGAAGATGGAAGCGTTCAACTAGCAGACCATTATCAACAAAATACT
CCAATTGGCGATGGCCCTGTCCTTTTACCAGACAACCATTACCTGTCCACACAATCTGCCCTTTCGAAAGATCCCAACGAAAAGAGAGACCAC
ATGGTCCTTCTTGAGTTTGTAACAGCTGCTGGGATTACACATGGCATGGATGAACTATACAAATAAGGAAACACAGAAAAAAGCCCGCACCTG
ACAGTGCGGGCTTTTTTTTTCGACCAAAGGGGCTTTTTTGTTATTTCTGCAG

**pSB1A3-I13521**

pLTetO-1 (BBa_R0040) RBS mRFP (BBa_E1010) terminator

plasmid map: https://benchling.com/s/33pnk7CI

GAATTCGCGGCCGCTTCTAGAGTCCCTATCAGTGATAGAGATTGACATCCCTATCAGTGATAGAGATACTGAGCACTACTAGAGAAAGAGGAG
AAATACTAGATGGCTTCCTCCGAAGACGTTATCAAAGAGTTCATGCGTTTCAAAGTTCGTATGGAAGGTTCCGTTAACGGTCACGAGTTCGAAA
TCGAAGGTGAAGGTGAAGGTCGTCCGTACGAAGGTACCCAGACCGCTAAACTGAAAGTTACCAAAGGTGGTCCGCTGCCGTTCGCTTGGGAC
ATCCTGTCCCCGCAGTTCCAGTACGGTTCCAAAGCTTACGTTAAACACCCGGCTGACATCCCGGACTACCTGAAACTGTCCTTCCCGGAAGGT
TTCAAATGGGAACGTGTTATGAACTTCGAAGACGGTGGTGTTGTTACCGTTACCCAGGACTCCTCCCTGCAAGACGGTGAGTTCATCTACAAA
GTTAAACTGCGTGGTACCAACTTCCCGTCCGACGGTCCGGTTATGCAGAAAAAAACCATGGGTTGGGAAGCTTCCACCGAACGTATGTACCC
GGAAGACGGTGCTCTGAAAGGTGAAATCAAAATGCGTCTGAAACTGAAAGACGGTGGTCACTACGACGCTGAAGTTAAAACCACCTACATGG
CTAAAAAACCGGTTCAGCTGCCGGGTGCTTACAAAACCGACATCAAACTGGACATCACCTCCCACAACGAAGACTACACCATCGTTGAACAGT
ACGAACGTGCTGAAGGTCGTCACTCCACCGGTGCTTAATAACGCTGATAGTGCTAGTGTAGATCGCTACTAGAGCCAGGCATCAAATAAAACG
AAAGGCTCAGTCGAAAGACTGGGCCTTTCGTTTTATCTGTTGTTTGTCGGTGAACGCTCTCTACTAGAGTCACACTGGCTCACCTTCGGGTGG
GCCTTTCTGCGTTTATATACTAGTAGCGGCCGCTGCAG

**pSB1C3-AD052**

pLacO-1, ribozyme, RBS, TetR, YFP, terminator

plasmid map: https://benchling.com/s/seq-rbG6fGZqB6NoKi8FToH6

GAATTCATTCTGATAAATGTGAGCGGATAACATTGACATTGTGAGCGGATAACAAGATACTGAGCACCTGAGCGCTCAACGGGTGTGCTTCCC
GTTCTGATGAGTCCGTGAGGACGAAAGCGCCTCTACAAATAATTTTGTTTAACCCCCCGAGGAGTAGCACATGTCCAGATTAGATAAAAGTAAA
GTGATTAACAGCGCATTAGAGCTGCTTAATGAGGTCGGAATCGAAGGTTTAACAACCCGTAAACTCGCCCAGAAGCTAGGTGTAGAGCAGCCT
ACATTGTATTGGCATGTAAAAAATAAGCGGGCTTTGCTCGACGCCTTAGCCATTGAGATGTTAGATAGGCACCATACTCACTTTTGCCCTTTAG
AAGGGGAAAGCTGGCAAGATTTTTTACGTAATAACGCTAAAAGTTTTAGATGTGCTTTACTAAGTCATCGCGATGGAGCAAAAGTACATTTAGG
TACACGGCCTACAGAAAAACAGTATGAAACTCTCGAAAATCAATTAGCCTTTTTATGCCAACAAGGTTTTTCACTAGAGAATGCATTATATGCAC
TCAGCGCTGTGGGGCATTTTACTTTAGGTTGCGTATTGGAAGATCAAGAGCATCAAGTCGCTAAAGAAGAAAGGGAAACACCTACTACTGATA
GTATGCCGCCATTATTACGACAAGCTATCGAATTATTTGATCACCAAGGTGCAGAGCCAGCCTTCTTATTCGGCCTTGAATTGATCATATGCGG
ATTAGAAAAACAACTTAAATGTGAAAGTGGGTCCTAATAAGCTCCGGCAAGCAATATCTGAGTAGTCACCGGCTGTGCTTGCCGGTCTGATGA
GCCTGTGAAGGCGAAACTACCTCTACAAATAATTTTGTTTAACTATGGACTATGTTTTCACATACGAGGGGGATTAGATGCGTAAAGGAGAAGA
ACTTTTCACTGGAGTTGTCCCAATCTTAGTAGAATTGGACGGAGACGTCAATGGACACAAGTTCTCCGTTTCGGGAGAGGGGGAGGGGGATG
CGACGTATGGTAAATTGACGCTGAAGCTGTTATGTACCACCGGTAAGTTGCCTGTGCCGTGGCCCACATTAGTAACAACCCTGGGGTACGGC
GTGCAATGCTTCGCGCGTTATCCTGACCATATGAAGCAACATGACTTCTTTAAGTCTGCCATGCCTGAAGGATATGTTCAAGAGCGCACGATTT
TTTTTAAAGATGATGGGAATTATAAAACTCGTGCAGAAGTGAAATTTGAGGGCGACACATTAGTGAACCGCATCGAATTAAAAGGAATCGACTT
TAAAGAAGATGGAAACATTTTGGGGCACAAATTGGAGTACAATTATAACAGCCATAACGTCTATATCACAGCGGATAAGCAGAAGAATGGCATC
AAAGCGAATTTTAAGATTCGTCACAATATTGAAGATGGGGGAGTTCAACTGGCGGACCATTATCAACAAAATACACCGATCGGTGACGGACCG
GTGTTGTTGCCGGACAACCACTATTTATCATATCAGAGCGCGTTATTTAAGGACCCGAACGAGAAGCGTGACCACATGGTCCTGTTGGAATTT
CTGACAGCCGCTGGTATTACTGAGGGTATGAATGAGCTTTACAAATAAGGAAACACAGAAAAAAGCCCGCACCTGACAGTGCGGGCTTTTTTT
TTCGACCAAAGGGGCTTTTTTGTTATTTCTGCAG

**pSB1C3-AD085**

pLacO-1, ribozyme, RBS, YFP, terminator

plasmid map: https://benchling.com/s/seq-XZwOKqviHo11LnkCaf7M

GAATTCATTCTGATAAATGTGAGCGGATAACATTGACATTGTGAGCGGATAACAAGATACTGAGCACCTGAGTAGTCACCGGCTGTGCTTGCC
GGTCTGATGAGCCTGTGAAGGCGAAACTACCTCTACAAATAATTTTGTTTAACTATGGACTATGTTTTCACATACGAGGGGGATTAGATGCGTA
AAGGAGAAGAACTTTTCACTGGAGTTGTCCCAATCTTAGTAGAATTGGACGGAGACGTCAATGGACACAAGTTCTCCGTTTCGGGAGAGGGG
GAGGGGGATGCGACGTATGGTAAATTGACGCTGAAGCTGTTATGTACCACCGGTAAGTTGCCTGTGCCGTGGCCCACATTAGTAACAACCCT
GGGGTACGGCGTGCAATGCTTCGCGCGTTATCCTGACCATATGAAGCAACATGACTTCTTTAAGTCTGCCATGCCTGAAGGATATGTTCAAGA

GCGCACGATTTTTTTTAAAGATGATGGGAATTATAAAACTCGTGCAGAAGTGAAATTTGAGGGCGACACATTAGTGAACCGCATCGAATTAAAA
GGAATCGACTTTAAAGAAGATGGAAACATTTTGGGGCACAAATTGGAGTACAATTATAACAGCCATAACGTCTATATCACAGCGGATAAGCAGA
AGAATGGCATCAAAGCGAATTTTAAGATTCGTCACAATATTGAAGATGGGGGAGTTCAACTGGCGGACCATTATCAACAAAATACACCGATCG
GTGACGGACCGGTGTTGTTGCCGGACAACCACTATTTATCATATCAGAGCGCGTTATTTAAGGACCCGAACGAGAAGCGTGACCACATGGTC
CTGTTGGAATTTCTGACAGCCGCTGGTATTACTGAGGGTATGAATGAGCTTTACAAATAAGGAAACACAGAAAAAAGCCCGCACCTGACAGTG
CGGGCTTTTTTTTTCGACCAAAGGGGCTTTTTTGTTATTTCTGCAG

**pSB1A3-AD064**

pTetO, ribozyme, RBS, LacI, degradation tag, mScarletI (RFP), terminator

plasmid map: https://benchling.com/s/seq-KQsGFrAIjH4Iy89qRSD7

GAATTCTACTCCACCGTTGGCTTTTTTCCCTATCAGTGATAGAGATTGACATCCCTATCAGTGATAGAGATAATGAGCACCTGAGGGGTCAGTT
GATGTGCTTTCAACTCTGATGAGTCAGTGATGACGAAACCCCCTCTACAAATAATTTTGTTTAACCCCCCGAGGAGTAGCACATGAAACCAGTA
ACGTTATACGATGTCGCAGAGTATGCCGGTGTCTCTTATCAGACCGTTTCCCGCGTGGTGAACCAGGCCAGCCACGTTTCTGCGAAAACGCG
GGAAAAAGTGGAAGCGGCGATGGCGGAGCTGAATTACATTCCCAACCGCGTGGCACAACAACTGGCGGGCAAACAGTCGTTGCTGATTGGC
GTTGCCACCTCCAGTCTGGCCCTGCACGCGCCGTCGCAAATTGTCGCGGCGATTAAATCTCGCGCCGATCAACTGGGTGCCAGCGTGGTGG
TGTCGATGGTAGAACGAAGCGGCGTCGAAGCCTGTAAAGCGGCGGTGCACAATCTTCTCGCGCAACGCGTCAGTGGGCTGATCATTAACTAT
CCGCTGGATGACCAGGATGCCATTGCTGTGGAAGCTGCCTGCACTAATGTTCCGGCGTTATTTCTTGATGTCTCTGACCAGACACCCATCAAC
AGTATTATTTTCTCCCATGAGGACGGTACGCGACTGGGCGTGGAGCATCTGGTCGCATTGGGTCACCAGCAAATCGCGCTGTTAGCGGGCCC
ATTAAGTTCTGTCTCGGCGCGTCTGCGTCTGGCTGGCTGGCATAAATATCTCACTCGCAATCAAATTCAGCCGATAGCGGAACGGGAAGGCG
ACTGGAGTGCCATGTCCGGTTTTCAACAAACCATGCAAATGCTGAATGAGGGCATCGTTCCCACTGCGATGCTGGTTGCCAACGATCAGATGG
CGCTGGGCGCAATGCGCGCCATTACCGAGTCCGGGCTGCGCGTTGGTGCGGATATCTCGGTAGTGGGATACGACGATACCGAAGATAGCTC
ATGTTATATCCCGCCGTTAACCACCATCAAACAGGATTTTCGCCTGCTGGGGCAAACCAGCGTGGACCGCTTGCTGCAACTCTCTCAGGGCCA
GGCGGTGAAGGGCAATCAGCTGTTGCCAGTCTCACTGGTGAAAAGAAAAACCACCCTGGCGCCCAATACGCAAACCGCCTCTCCCCGCGCG
TTGGCCGATTCATTAATGCAGCTGGCACGACAGGTTTCCCGACTGGAAAGCGGGCAGCGTCCCGCAGCGAATGATGAGAATTACGCAGCCG
CCGTTTGATAATAAGCTCCGGCAAGCACCCATGGATATCTGAGCTGTCACCGGATGTGCTTTCCGGTCTGATGAGTCCGTGAGGACGAAACA
GCGAGATCAGATAATTCTGACGTACAATTTACAAGTCATTAGGAGGATCTAAAATGGTGTCAAAGGGAGAGGCGGTTATCAAGGAATTTATGC
GCTTTAAAGTCCACATGGAGGGCAGCATGAACGGGCACGAGTTTGAAATTGAGGGGGAGGGGGAGGGCCGTCCTTATGAAGGTACTCAGAC
TGCTAAACTGAAGGTGACAAAAGGTGGCCCCTTGCCTTTCTCGTGGGACATCCTGTCGCCACAATTCATGTACGGGAGCCGCGCCTTTATCAA
ACATCCCGCAGATATTCCTGATTACTATAAACAATCTTTCCCGGAAGGTTTCAAATGGGAACGCGTCATGAATTTTGAGGACGGGGGCGCTGT
CACAGTTACTCAGGACACCTCCTTGGAAGACGGCACATTGATTTACAAGGTTAAGTTGCGCGGCACAAACTTCCCCCCTGACGGCCAGTAAT
GCAAAAGAAAACTATGGGTTGGGAGGCGTCTACAGAACGTTTATACCCCGAAGACGGGGTGCTGAAAGGTGACATTAAGATGGCCCTGCGCC
TGAAGGACGGCGGTCGCTATCTTGCCGACTTTAAAACTACTTATAAGGCTAAAAAACCAGTCCAGATGCCAGGCGCCTATAATGTTGACCGCA
AGTTAGACATCACCTCACATAATGAAGACTATACCGTTGTAGAACAATACGAGCGCAGCGAGGGTCGTCACAGTACCGGGGGGATGGATGAA
TTATACAAACGCCCGGCGGCGAACGATGAAAACTATGCGGCAGCGGTTTAATTCAGCCAAAAAACTTAAGACCGCCGGTCTTGTCCACTACCT
TGCAGTAATGCGGTGGACAGGATCGGCGGTTTTCTTTTCTCTTCTCAACTGCAG

**pSB1A3-AD084**

pTetO, ribozyme, RBS, LacI, mScarletI (RFP), degradation tag, terminator

plasmid map: https://benchling.com/s/seq-QEuQ3FexjvMwOcQ27tIV

GAATTCTACTCCACCGTTGGCTTTTTTCCCTATCAGTGATAGAGATTGACATCCCTATCAGTGATAGAGATAATGAGCACCTGAGGGGTCAGTT
GATGTGCTTTCAACTCTGATGAGTCAGTGATGACGAAACCCCCTCTACAAATAATTTTGTTTAACCCCCCGAGGAGTAGCACATGAAACCAGTA
ACGTTATACGATGTCGCAGAGTATGCCGGTGTCTCTTATCAGACCGTTTCCCGCGTGGTGAACCAGGCCAGCCACGTTTCTGCGAAAACGCG
GGAAAAAGTGGAAGCGGCGATGGCGGAGCTGAATTACATTCCCAACCGCGTGGCACAACAACTGGCGGGCAAACAGTCGTTGCTGATTGGC
GTTGCCACCTCCAGTCTGGCCCTGCACGCGCCGTCGCAAATTGTCGCGGCGATTAAATCTCGCGCCGATCAACTGGGTGCCAGCGTGGTGG
TGTCGATGGTAGAACGAAGCGGCGTCGAAGCCTGTAAAGCGGCGGTGCACAATCTTCTCGCGCAACGCGTCAGTGGGCTGATCATTAACTAT
CCGCTGGATGACCAGGATGCCATTGCTGTGGAAGCTGCCTGCACTAATGTTCCGGCGTTATTTCTTGATGTCTCTGACCAGACACCCATCAAC
AGTATTATTTTCTCCCATGAGGACGGTACGCGACTGGGCGTGGAGCATCTGGTCGCATTGGGTCACCAGCAAATCGCGCTGTTAGCGGGCCC
ATTAAGTTCTGTCTCGGCGCGTCTGCGTCTGGCTGGCTGGCATAAATATCTCACTCGCAATCAAATTCAGCCGATAGCGGAACGGGAAGGCG
ACTGGAGTGCCATGTCCGGTTTTCAACAAACCATGCAAATGCTGAATGAGGGCATCGTTCCCACTGCGATGCTGGTTGCCAACGATCAGATGG
CGCTGGGCGCAATGCGCGCCATTACCGAGTCCGGGCTGCGCGTTGGTGCGGATATCTCGGTAGTGGGATACGACGATACCGAAGATAGCTC
ATGTTATATCCCGCCGTTAACCACCATCAAACAGGATTTTCGCCTGCTGGGGCAAACCAGCGTGGACCGCTTGCTGCAACTCTCTCAGGGCCA
GGCGGTGAAGGGCAATCAGCTGTTGCCAGTCTCACTGGTGAAAAGAAAAACCACCCTGGCGCCCAATACGCAAACCGCCTCTCCCCGCGCG
TTGGCCGATTCATTAATGCAGCTGGCACGACAGGTTTCCCGACTGGAAAGCGGGCAGTGATAATAAGCTCCGGCAAGCACCCATGGATATCT
GAGCTGTCACCGGATGTGCTTTCCGGTCTGATGAGTCCGTGAGGACGAAACAGCGAGATCAGATAATTCTGACGTACAATTTACAAGTCATTA
GGAGGATCTAAAATGGTGTCAAAGGGAGAGGCGGTTATCAAGGAATTTATGCGCTTTAAAGTCCACATGGAGGGCAGCATGAACGGGCACGA
GTTTGAAATTGAGGGGGAGGGGGAGGGCCGTCCTTATGAAGGTACTCAGACTGCTAAACTGAAGGTGACAAAAGGTGGCCCCTTGCCTTTCT
CGTGGGACATCCTGTCGCCACAATTCATGTACGGGAGCCGCGCCTTTATCAAACATCCCGCAGATATTCCTGATTACTATAAACAATCTTTCCC
GGAAGGTTTCAAATGGGAACGCGTCATGAATTTTGAGGACGGGGGCGCTGTCACAGTTACTCAGGACACCTCCTTGGAAGACGGCACATTGA
TTTACAAGGTTAAGTTGCGCGGCACAAACTTCCCCCCTGACGGCCAGTAATGCAAAAGAAAACTATGGGTTGGGAGGCGTCTACAGAACGTT
TATACCCCGAAGACGGGGTGCTGAAAGGTGACATTAAGATGGCCCTGCGCCTGAAGGACGGCGGTCGCTATCTTGCCGACTTTAAAACTACT
TATAAGGCTAAAAAACCAGTCCAGATGCCAGGCGCCTATAATGTTGACCGCAAGTTAGACATCACCTCACATAATGAAGACTATACCGTTGTAG
AACAATACGAGCGCAGCGAGGGTCGTCACAGTACCGGGGGGATGGATGAATTATACAAACGCCCGGCGGCGAACGATGAAAACTATGCGGC
AGCGGTTTAATTCAGCCAAAAAACTTAAGACCGCCGGTCTTGTCCACTACCTTGCAGTAATGCGGTGGACAGGATCGGCGGTTTTCTTTTCTCT
TCTCAACTGCAG

**pSB1C3-pLacO-TetR-GFP**

pLacO, ribozyme, RBS, TetR, GFPmut3 (BBa_E0040), terminator

plasmid map: https://benchling.com/s/YyqP4114

GAATTCATTCTGATAAATGTGAGCGGATAACATTGACATTGTGAGCGGATAACAAGATACTGAGCACCTGAGCGCTCAACGGGTGTGCTTCCC
GTTCTGATGAGTCCGTGAGGACGAAAGCGCCTCTACAAATAATTTTGTTTAACCCCCCGAGGAGTAGCACATGTCCAGATTAGATAAAAGTAAA
GTGATTAACAGCGCATTAGAGCTGCTTAATGAGGTCGGAATCGAAGGTTTAACAACCCGTAAACTCGCCCAGAAGCTAGGTGTAGAGCAGCCT
ACATTGTATTGGCATGTAAAAAATAAGCGGGCTTTGCTCGACGCCTTAGCCATTGAGATGTTAGATAGGCACCATACTCACTTTTGCCCTTTAG
AAGGGGAAAGCTGGCAAGATTTTTTACGTAATAACGCTAAAAGTTTTAGATGTGCTTTACTAAGTCATCGCGATGGAGCAAAAGTACATTTAGG
TACACGGCCTACAGAAAAACAGTATGAAACTCTCGAAAATCAATTAGCCTTTTTATGCCAACAAGGTTTTTCACTAGAGAATGCATTATATGCAC
TCAGCGCTGTGGGGCATTTTACTTTAGGTTGCGTATTGGAAGATCAAGAGCATCAAGTCGCTAAAGAAGAAAGGGAAACACCTACTACTGATA
GTATGCCGCCATTATTACGACAAGCTATCGAATTATTTGATCACCAAGGTGCAGAGCCAGCCTTCTTATTCGGCCTTGAATTGATCATATGCGG
ATTAGAAAAACAACTTAAATGTGAAAGTGGGTCCTAATAAGCTCCGGCAAGCAATATCTGAGTAGTCACCGGCTGTGCTTGCCGGTCTGATGA
GCCTGTGAAGGCGAAACTACCTCTACAAATAATTTTGTTTAACTATGGACTATGTTTCACATACGAGGGGGATTAGATGCGTAAAGGAGAAGA
ACTTTTCACTGGAGTTGTCCCAATTCTTGTTGAATTAGATGGTGATGTTAATGGGCACAAATTTTCTGTCAGTGGAGAGGGTGAAGGTGATGCA
ACATACGGAAAACTTACCCTTAAATTTATTTGCACTACTGGAAAACTACCTGTTCCATGGCCAACACTTGTCACTACTTTCGGTTATGGTGTTCA
ATGCTTTGCGAGATACCCAGATCATATGAAACAGCATGACTTTTTCAAGAGTGCCATGCCCGAAGGTTATGTACAGGAAAGAACTATATTTTTC
AAAGATGACGGGAACTACAAGACACGTGCTGAAGTCAAGTTTGAAGGTGATACCCTTGTTAATAGAATCGAGTTAAAAGGTATTGATTTTAAAG
AAGATGGAAACATTCTTGGACACAAATTGGAATACAACTATAACTCACACAATGTATACATCATGGCAGACAAACAAAAGAATGGAATCAAAGTT
AACTTCAAAATTAGACACAACATTGAAGATGGAAGCGTTCAACTAGCAGACCATTATCAACAAAATACTCCAATTGGCGATGGCCCTGTCCTTT
TACCAGACAACCATTACCTGTCCACACAATCTGCCCTTTCGAAAGATCCCAACGAAAAGAGAGACCACATGGTCCTTCTTGAGTTTGTAACAGC
TGCTGGGATTACACATGGCATGGATGAACTATACAAATAAGGAAACACAGAAAAAAGCCCGCACCTGACAGTGCGGGCTTTTTTTTTCGACCA
AAGGGGCTTTTTTGTTATTTCTGCAG

**pSB1A3-pTetO-LacI-mCherry**

pTetO, ribozyme, RBS, LacI, mCherry, degradation tag, terminator

plasmid map: https://benchling.com/s/seq-N5gMAp7aZ0pYflaixmlS

GAATTCTACTCCACCGTTGGCTTTTTTCCCTATCAGTGATAGAGATTGACATCCCTATCAGTGATAGAGATAATGAGCACCTGAGGGGTCAGTT
GATGTGCTTTCAACTCTGATGAGTCAGTGATGACGAAACCCCCTCTACAAATAATTTTGTTTAACCCCCCGAGGAGTAGCACATGAAACCAGTA
ACGTTATACGATGTCGCAGAGTATGCCGGTGTCTCTTATCAGACCGTTTCCCGCGTGGTGAACCAGGCCAGCCACGTTTCTGCGAAAACGCG
GGAAAAAGTGGAAGCGGCGATGGCGGAGCTGAATTACATTCCCAACCGCGTGGCACAACAACTGGCGGGCAAACAGTCGTTGCTGATTGGC
GTTGCCACCTCCAGTCTGGCCCTGCACGCGCCGTCGCAAATTGTCGCGGCGATTAAATCTCGCGCCGATCAACTGGGTGCCAGCGTGGTGG
TGTCGATGGTAGAACGAAGCGGCGTCGAAGCCTGTAAAGCGGCGGTGCACAATCTTCTCGCGCAACGCGTCAGTGGGCTGATCATTAACTAT
CCGCTGGATGACCAGGATGCCATTGCTGTGGAAGCTGCCTGCACTAATGTTCCGGCGTTATTTCTTGATGTCTCTGACCAGACACCCATCAAC
AGTATTATTTTCTCCCATGAGGACGGTACGCGACTGGGCGTGGAGCATCTGGTCGCATTGGGTCACCAGCAAATCGCGCTGTTAGCGGGCCC
ATTAAGTTCTGTCTCGGCGCGTCTGCGTCTGGCTGGCTGGCATAAATATCTCACTCGCAATCAAATTCAGCCGATAGCGGAACGGGAAGGCG
ACTGGAGTGCCATGTCCGGTTTTCAACAAACCATGCAAATGCTGAATGAGGGCATCGTTCCCACTGCGATGCTGGTTGCCAACGATCAGATGG
CGCTGGGCGCAATGCGCGCCATTACCGAGTCCGGGCTGCGCGTTGGTGCGGATATCTCGGTAGTGGGATACGACGATACCGAAGATAGCTC
ATGTTATATCCCGCCGTTAACCACCATCAAACAGGATTTTCGCCTGCTGGGGCAAACCAGCGTGGACCGCTTGCTGCAACTCTCTCAGGGCCA
GGCGGTGAAGGGCAATCAGCTGTTGCCAGTCTCACTGGTGAAAAGAAAAACCACCCTGGCGCCCAATACGCAAACCGCCTCTCCCCGCGCG
TTGGCCGATTCATTAATGCAGCTGGCACGACAGGTTTCCCGACTGGAAAGCGGGCAGTGATAATAAGCTCCGGCAAGCACCCATGGATATCT
GAGAAGTCAATTAATGTGCTTTTAATTCTGATGAGTCGGTGACGACGAAACTTCCTCTACAAATAATTTTGTTTAACTATGGACTATGTTTTAACT
ACTAGATGGTCAGCAAGGGCGAGGAGGATAACATGGCCATCATCAAGGAGTTCATGCGCTTCAAGGTGCACATGGAGGGCTCCGTGAACGG
CCACGAGTTCGAGATCGAGGGCGAGGGCGAGGGCCGCCCCTACGAGGGCACCCAGACCGCCAAGCTGAAGGTGACCAAGGGTGGCCCCC
TGCCCTTCGCCTGGGACATCCTGTCCCCTCAGTTCATGTACGGCTCCAAGGCCTACGTGAAGCACCCCGCCGACATCCCCGACTACTTGAAG
CTGTCCTTCCCCGAGGGCTTCAAGTGGGAGCGCGTGATGAACTTCGAGGACGGCGGCGTGGTGACCGTGACCCAGGACTCCTCCCTGCAAG
ACGGCGAGTTCATCTACAAGGTGAAGCTGCGCGGCACCAACTTCCCCTCCGACGGCCCCGTAATGCAGAAGAAGACTATGGGCTGGGAGGC
CTCCTCCGAGCGGATGTACCCCGAGGACGGCGCCCTGAAGGGCGAGATCAAGCAGAGGCTGAAGCTGAAGGACGGCGGCCACTACGACGC
TGAGGTCAAGACCACCTACAAGGCCAAGAAGCCCGTGCAGCTGCCCGGCGCCTACAACGTCAACATCAAGTTGGACATCACCTCCCACAACG
AGGACTACACCATCGTGGAACAGTACGAACGCGCCGAGGGCCGCCACTCCACCGGCGGCATGGACGAGCTGTACAAGCGCCCGGCGGCGA
ACGATGAAAACTATGCGGCAGCGGTTTAATTCAGCCAAAAAACTTAAGACCGCCGGTCTTGTCCACTACCTTGCAGTAATGCGGTGGACAGGA
TCGGCGGTTTTCTTTTCTCTTCTCAACTGCAG
